# Bioorthogonal chemical labelling of endogenous neurotransmitter receptors in living mouse brains

**DOI:** 10.1101/2023.01.16.524180

**Authors:** Hiroshi Nonaka, Seiji Sakamoto, Kazuki Shiraiwa, Mamoru Ishikawa, Tomonori Tamura, Kyohei Okuno, Shigeki Kiyonaka, Etsuo A. Susaki, Chika Shimizu, Hiroki R. Ueda, Wataru Kakegawa, Itaru Arai, Michisuke Yuzaki, Itaru Hamachi

**Affiliations:** Department of Synthetic Chemistry and Biological Chemistry, Graduate School of Engineering, Kyoto University, Nishikyo-ku, Kyoto, 615-8510, Japan; JST-ERATO, Hamachi Innovative Molecular Technology for Neuroscience, Kyoto, Japan; Department of Biomolecular Engineering, Graduate School of Engineering, Nagoya University, Nagoya, 464-8603, Japan; Department of Biochemistry and Systems Biomedicine, Juntendo University Graduate School of Medicine, Tokyo, 113-8421, Japan; Department of Systems Pharmacology, Graduate School of Medicine, The University of Tokyo, Tokyo, 113-0033, Japan; Laboratory for Synthetic Biology, RIKEN Center for Biosystems Dynamics Research, Osaka, Japan; Department of Neurophysiology, Keio University School of Medicine, Tokyo, 160-8582, Japan

## Abstract

Neurotransmitter receptors are essential components of synapses for communication between neurons in the brain. Because the spatiotemporal expression profiles and dynamics of neurotransmitter receptors involved in many functions are delicately governed in the brain, *in vivo* research tools with high spatiotemporal resolution for receptors in intact brains are highly desirable. Covalent chemical labelling of proteins without genetic manipulation is now a powerful method for analyzing receptors *in vitro*. However, selective target receptor labelling in the brain has not yet been achieved. This study shows that ligand-directed alkoxyacylimidazole (LDAI) chemistry can be used to selectively tether synthetic probes to target endogenous receptors in living mouse brains. The reactive LDAI reagents with negative charges were found to diffuse well over the whole brain and could selectively label target endogenous receptors, including AMPAR, NMDAR, mGlu1, and GABA_A_R. This simple and robust labelling protocol was then used for various applications: three-dimensional spatial mapping of endogenous receptors in the brains of healthy and disease-model mice; multi-colour receptor imaging; and pulse-chase analysis of the receptor dynamics in postnatal mouse brains. Here, results demonstrated that bioorthogonal receptor modification in living animal brains may provide innovative molecular tools that contribute to the in-depth understanding of complicated brain functions.

## Main text

The brain is arguably the most complex system in the body. Each neuron in the brain is connected by nanoscale synapses to other neurons forming elaborate cell-to-cell networks. In the brain neural circuits, neurotransmitter receptors on postsynaptic membranes are the starting points of signalling cascades and control synapse formation, maintenance, plasticity, and function. The subcellular localization and cell surface densities of different receptors are dynamically altered in response to changes in neuronal activity during development and are involved in high-order brain functions.^1^ Most of the current knowledge of receptor dynamics and localization has been accumulated from *ex vivo* experiments (i.e., dissociated neurons and brain tissue slices) because the methods available for *in vivo* analysis are limited. However, given that receptor dynamics and functions are delicately governed by the complex, three-dimensionally connected network of the brain, receptors in the living brains of animals should be directly studied.^2^

Various approaches have been recently investigated to examine the dynamics of neurotransmitter receptors in animal brains.^3^ For example, the virus-mediated expression of fluorescent protein-fused receptors has been widely used to investigate the nanoscale organization of receptors in living mouse brain tissues.^4^ However, transient transduction is now recognized to possibly cause overexpression of the modified receptors, resulting in mistargeting and dysregulation.^5^ Recent advances in genetic engineering, especially in CRISPR gene editing, have facilitated the labelling of endogenous proteins.^6,7^ However, the exogenous reporter tag may perturb the physiologically balanced (hetero)oligomeric structure, function, and trafficking of the receptor, as well as the stability of its mRNA.^8^This perturbation is also an issue for chemogenetic labelling methods, such as Halo-tag and SNAP-tag.^9^

Covalent protein labelling can be used to attach desired functional probes to endogenous receptors without genetic manipulation.^10^ Despite some successful examples *in vitro* and *ex vivo*,^11–14^ no chemical labelling of neurotransmitter receptors *in vivo* has been achieved to date. This lack of chemical labelling *in vivo* is mainly because (i) general concerns regarding the target protein selectivity of chemical reactions in the highly complicated brain that contains numerous non-target molecules; (ii) the lack of an established method for efficiently delivering reactive molecules into the brain; and (iii) poor information regarding the appropriate physico-chemical properties (e.g., diffusibility, distribution properties, and reaction kinetics) of chemically reactive small molecules for the covalent labelling of receptors in the brain. If these obstacles can be removed, the covalent tagging of endogenous receptors in the brain would have valuable applications in neuroscience.

Here, we describe a ligand-directed acyl substitution reaction that enabled the selective chemical labelling of a target receptor in a living mouse brain (Figure 1a). The present study revealed that direct injection protocols similar to conventional virus injections were useful for the efficient delivery of reactive small molecules to the mouse brain. We found that the diffusibility in the brain greatly depended on the anionic charge of the reagent. These findings provided valuable guidelines for the design of chemical labelling reagents for use in the brain, and the selective labelling of various neurotransmitter receptors, a-amino-3-hydroxy-5-methyl-4-isoxazolepropionic acid receptors (AMPAR), *N*-methyl-D-aspartate receptors (NMDAR), a metabotropic glutamate receptor 1 (mGlu1), and ionotropic g-aminobutyric acid receptors (GABA_A_R), was achieved (Figure 1b, Extended Data Figure 1). The in-brain ligand-directed (LD) chemistry enabled various applications, including the visualization of the three-dimensional (3D) distribution of endogenous receptors, not only in the brain of a normal mouse, but also in a transgenic mouse model of Alzheimer’s disease, and the multi-colour imaging of different receptors. A detailed reaction kinetics study was conducted to determine both the labelling (acyl transfer reaction) rate and the lifetimes of functionally active receptors in living brains. Finally, we established a protocol for pulse-chase analysis of the endogenous AMPAR dynamics in cerebellar Purkinje cells (PCs) in the brains of developing mice. This technique was used to demonstrate that some of the AMPARs in the soma of PCs migrated to parallel fibre (PF) synapses generated at the distal dendrites during the development of neural circuits in the cerebellum.

**Figure 1.**
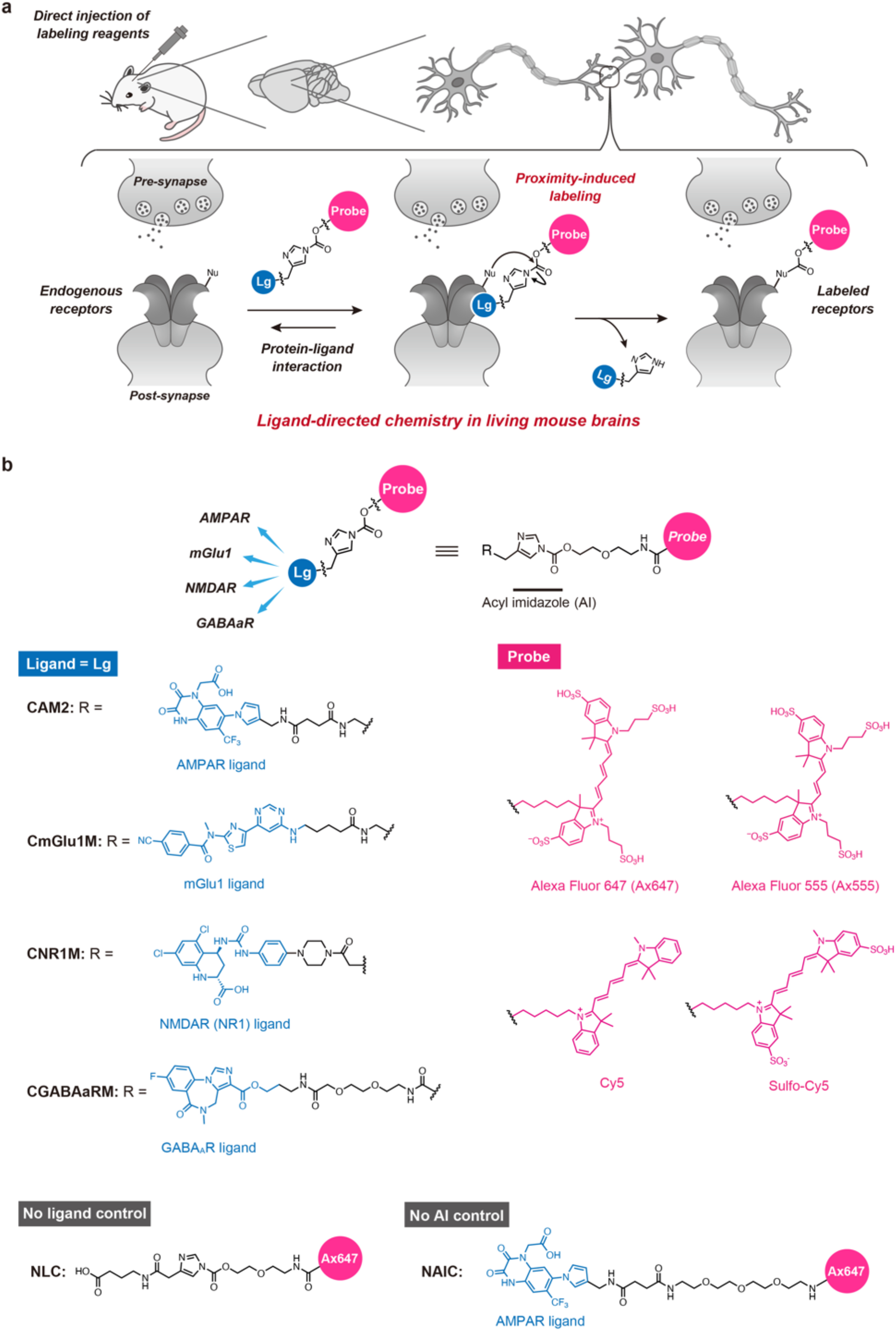
Chemical labelling of neurotransmitter receptors in the living mouse brain. **a**, Schematic illustration of the ligand-directed chemistry in the living mouse brain. Nu, nucleophilic amino-acid residue. Lg, selective ligand for each receptor. **b**, Chemical structures of labeling reagents for AMPAR, mGlu1, NMDAR, and GABA_A_R. Alexa Fluor 647 (Ax647) and Alexa Fluor 555 (Ax555) in this paper are putative structures. **NLC** and **NAIC** are control molecules without ligand or acyl imidazole moiety.

## Results

### Chemical labelling of AMPARs by acyl transfer reactions in living mouse brains

We chose AMPARs in the cerebellum as the initial target receptors to test our strategy for *in vivo* LD chemistry. AMPARs, which are heteromeric tetramers of GluA1–4,^15,16^ are abundantly expressed in the central nervous system and have a characteristic distribution in the cerebellar molecular layer, which can be used as a good indicator to evaluate the efficiency and specificity of the chemical labelling using imaging techniques. We have previously developed a series of ligand-directed alkoxyacylimidazole (LDAI)-based reagents,^10–13^ and **CAM2-Ax647** was selected for the present study because, among the developed compounds, this reagent had the least background signal derived from the autofluorescence of tissues. In addition, we expected that the hydrophilic and cell-impermeable features of this reagent would suppress nonspecific adsorption to hydrophobic materials in tissues and promote the labelling of only cell-surface (functional) receptors. For the delivery of **CAM2-Ax647** to living brains, we used a direct injection protocol that is widely used for virus injections. Initially, we injected the reagent (50 μM, 4.5 μL) into the cerebellum (Cb) region of live C57BL/6N mice (5-weeks old) under anaesthesia (Figure 2a). At 20 h after the injection of **CAM2-Ax647**, the mice appeared to behave normally (Supplementary Video 1). In western blot (WB) analysis of the Cb homogenates, a strong band corresponding to the labelled AMPAR (100 kDa, a GluA subunit) was detected using an anti-Ax647 antibody (Figure 2b). No bands were detected with a control compound lacking the ligand moiety (**NLC**) or vehicle (DMSO) control, indicating that the labelling was driven by selective ligand-receptor recognition. We also confirmed the high stability of the formed covalent bond (no degradation was observed after 8 days in Neurobasal medium, Supplementary Figure 1). Confocal laser scanning microscopy (CLSM) imaging of the Cb slices, which were prepared from a mouse injected with **CAM2-Ax647**, showed obvious and selective fluorescence signals of Ax647 from the molecular layer and almost negligible fluorescence from the granular layer (Figure 2c,d), which was consistent with the localization of endogenous AMPARs in the Cb.^17^ In contrast, only dim fluorescence was observed in the slices prepared from mice brains injected with **NLC** or DMSO. Whole-cell patch clamp recordings from PCs in the Cb acute slices were performed to examine the influence of the injection of **CAM2-Ax647** on the electrophysiological properties of AMPAR-mediated synaptic responses, such as PF- and climbing fibres (CF)-mediated excitatory postsynaptic current (PF- and CF-EPSC, respectively) amplitude and kinetics and the paired-pulse ratio of each EPSC, which reflect a presynaptic neurotransmitter release property (Figure 2e–p). Both the CF- and PF-EPSCs were identical between samples with and without **CAM2-Ax647** treatment. This result indicated that the detrimental impacts of the chemical labelling on the characteristics of AMPARs were minimal or negligible because the ligand moiety is cleaved upon labelling in the LD chemistry.

**Figure 2.**
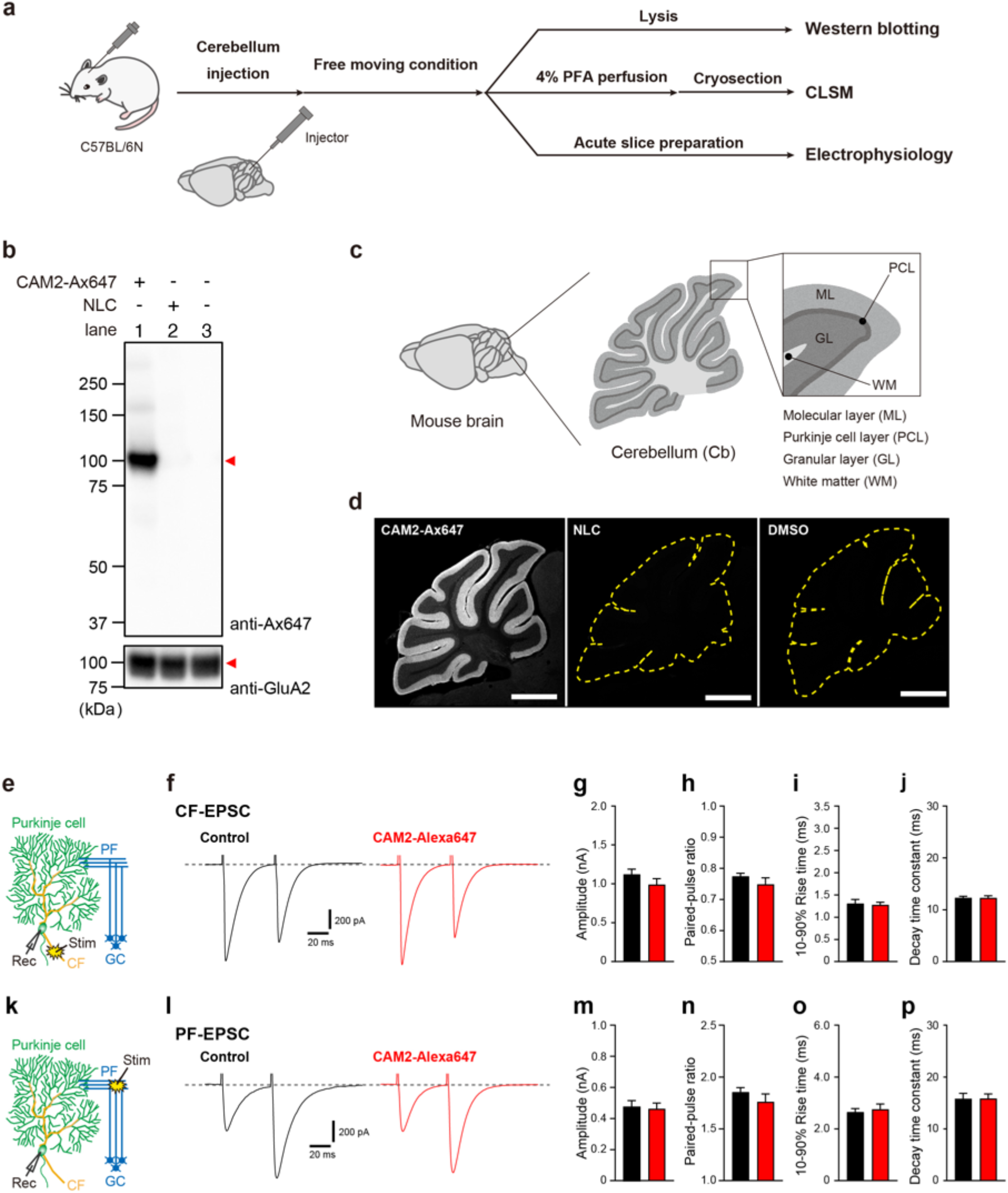
Chemical labelling of AMPA receptors in the live cerebellum. **a**, Experimental workflow in Figure 2. **b**, WB analysis of cerebellum homogenate administered with **CAM2-Ax647**, **NLC**, and DMSO. PBS(–) containing 50 μM of **CAM2-Ax647** (4.5 μL of 50 μM **CAM2-Ax647** which is assumed to be ca. 3.6 μM (C_cere_) on the basis of the cerebellum volume^50^) was injected into cerebellum of the mouse. At 24 h after injection, the mouse was sacrificed. The labelled brain was isolated, lysed with RIPA buffer, and subjected to WB analysis. **c**, Region names in the cerebellar sagittal section. **d**, Fluorescence images of labelled brains with **CAM2-Ax647**, **NLC**, or DMSO. 24 h after injection, the mouse was transcardially perfused with 4% PFA. The brain was isolated and sectioned by cryostat (50-μm thick). Imaging was performed using a CLSM equipped with a 5× objective (633 nm excitation for Ax647). Scale bar: 500 μm. **e-p**, *In vivo* chemical labelling with **CAM2-Ax647** does not affect AMPAR-mediated EPSCs in cerebellar slices. (**e**) An orientation of stimulus and recording electrodes to evoke CF-EPSCs. The cells were clamped at V_h_ = −10 mV. (**f**) Representative CF-EPSC traces recorded from cerebellar Purkinje cells in **CAM2-Ax647**-injected (right) and its control (left) mice. (**g** to **j**) Quantification of peak amplitude (**g**), paired-pulse ratio (**h**), 10–90% rise time (**i**) and decay time constant (**j**) of CF-EPSCs. (**k**) An orientation of stimulus and recording electrodes to evoke PF-EPSCs. The cells were clamped at V_h_ = −80 mV. (**l**) Representative PF-EPSC traces recorded from Purkinje cells in **CAM2-Ax647**-injected (right) and its control (left) mice. (**m** to **p**) Quantification of peak amplitude (**m**), paired-pulse ratio (**n**), 10–90% rise time (**o**) and decay time constant (**p**) of PF-EPSCs. Paired-pulse ratio of EPSCs was defined as the amplitude of 2^nd^ EPSC divided by that of 1^st^ EPSC. These results indicate that both CF-EPSC and PF-EPSC are not affected by the chemical labelling. n = 20 cells in each group for CF-EPSCs, and n =20 cells in each group for PF-EPSCs. Data are represented as mean ± SEM.

We next investigated lateral ventricle (LV) injection with the aim of labelling AMPARs over a wide brain region (Figure 3a). The mice administered **CAM2-Ax647** by LV injection showed no behavioural abnormalities (Supplementary Video 2). After labelling, the mouse brains were separated into Cb and non-Cb regions including the cerebral cortex, hippocampus, and striatum, and analyzed by WB. A single band corresponding to the labelled AMPARs was clearly observed at approximately 100 kDa (Figure 3b) from both Cb and non-Cb regions. Immunoprecipitation with anti-Ax647 followed by mass analysis (immunoprecipitation mass spectrometry: IP-MS) predominantly resulted in the detection of the four subunits of the AMPAR (GluA1–4), which had the highest enrichment scores, demonstrating the selective labelling of AMPARs with **CAM2-Ax647** over a wide brain region (Figure 3c and Supplementary Data 1). CLSM imaging of whole sagittal brain slices showed strong fluorescence signals of Ax647 from the hippocampus and the molecular layer of Cb and relatively weak but clear fluorescence from the cortex and striatum, indicating that reagents injected into the LV were well distributed over the whole brain tissues via the cerebrospinal fluid (CSF) flow.^18^ In contrast, these fluorescent signals were not observed in slices with the **NLC** or the non-reactive control reagent (**NAIC)** (Figure 3d). These negative control experiments indicated that the fluorescence observed with **CAM2-Ax647** injection came from covalently labelled AMPARs.

**Figure 3.**
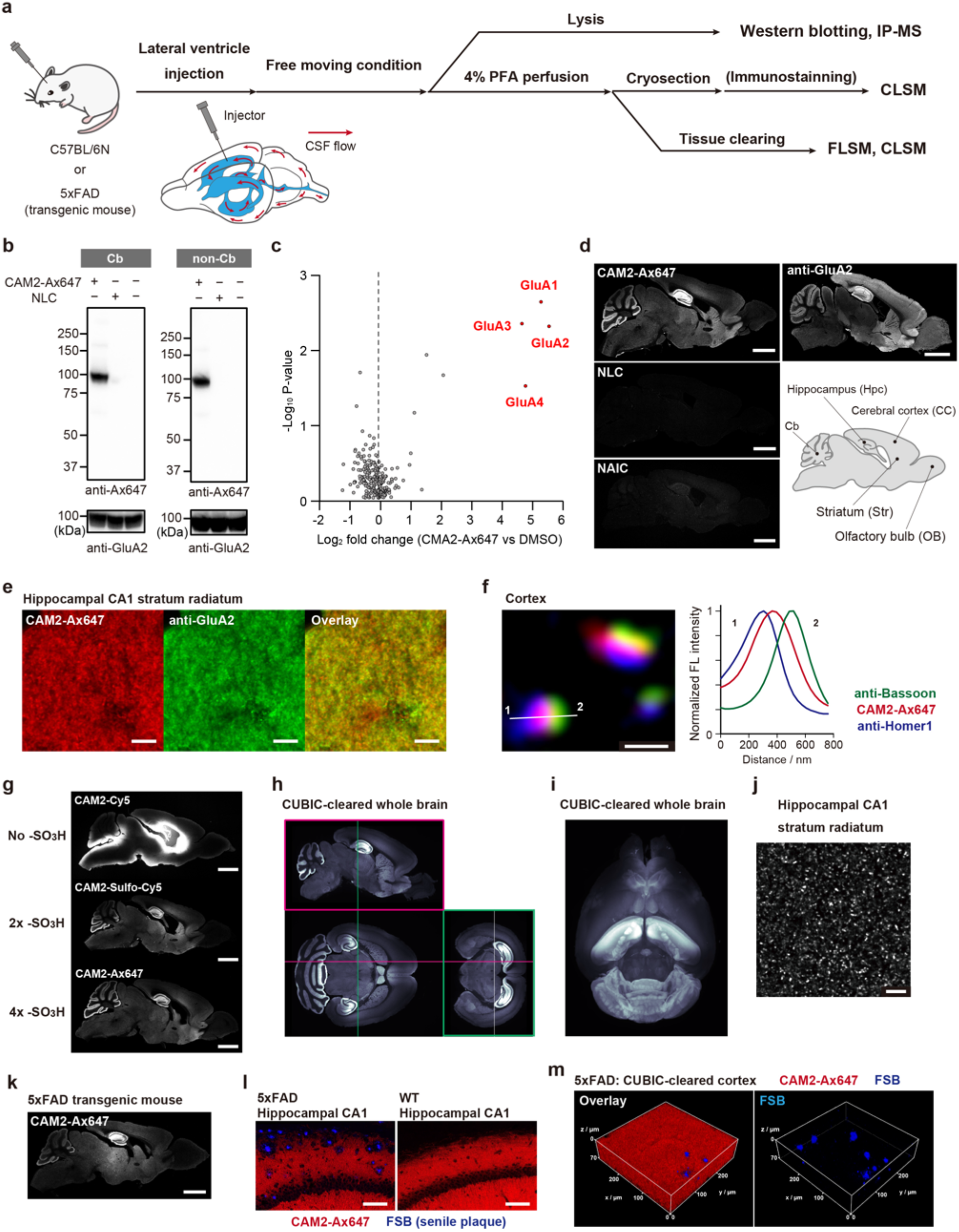
Chemical labelling of AMPA receptors in the living mouse brain. **a**, Experimental workflow in Figure 3. **b**, WB analysis of brain homogenate administered with **CAM2-Ax647** or **NLC**. PBS(–) containing 80 μM of **CAM2-Ax647** (4.5 μL: 4.5 μL of 80 μM **CAM2-Ax647** which is assumed to be ca. 0.8 μM (C_brain_) on the basis of the brain volume^50^) was injected into right LV of the mouse. 24 h after injection, the mouse was sacrificed. The labelled brain was isolated, lysed with RIPA buffer, and subjected to WB analysis. **c**, Volcano plot based on label-free quantification values for the proteins identified in the labelling experiment by LV injection with **CAM2-Ax647**. The dot representing GluA1–4 is labelled in red. **d**, Fluorescence images of labelled brains with **CAM2-Ax647**, **NLC**, or **NAIC** and immunostaining image with anti-GluA2. 24 h after LV injection, the mouse was transcardially perfused with 4% PFA/PBS(–). The brain was isolated and sectioned by cryostat (50-μm thick). Imaging was performed using a CLSM equipped with a 5× objective. Scale bar: 2 mm. **e**, High-resolution confocal image of labelled hippocampus with 100× lens. Scale bar: 5 μm. **f**, Localization analysis of labelling signals by using pre-and postsynaptic markers with a 100× objective and Leica Lightning deconvolution. Scale bar: 500 nm. **g**, Fluorescence images of labelled brains with **CAM2-Cy5**, **Sulfo-Cy5**, or **Ax647**. Labelling conditions are the same as Figure 3d. Scale bar: 2 mm. **h**, 3D fluorescence imaging of a tissue-cleared brain labelled with **CAM2-Ax647** by a LSFM. **i**, 3D rendering of Figure 3h. **j**, Super resolution image of hippocampus region. Fluorescence imaging was performed using a CLSM equipped with a 100× objective and Lightning deconvolution. Scale bar: 5 μm. **k**-**m**, Fluorescence images of labelled 5xFAD mouse brains with **CAM2-Ax647**. Labelling conditions are the same as Figure 3d. **k**. 5× objective. Scale bar: 2 mm. **l**. 100× objective. Scale bar: 100 μm. **m**, 3D fluorescence imaging of a tissue-cleared brain labelled with **CAM2-Ax647** by a CLSM with 40× objective. Colors: **CAM2-Ax647** (red) and FSB (for senile plaques staining, blue).

The AMPAR distribution visualized by Ax647 labelling in the whole brain was in good accordance with that observed with anti-GluA2 antibody staining (Figure 3d). The high-resolution images of the hippocampal CA1 region showed abundant fluorescent puncta of Ax647 and most of these puncta were colocalized with anti-GluA2 antibody signals (Figure 3e). These puncta were located in between pre- and post-synapses stained with anti-Bassoon and anti-Homer1a antibodies, respectively (Figure 3f). These data demonstrated that endogenous AMPARs accumulated in the synaptic cleft could be labelled and visualized with a synthetic Ax647 fluorophore. Collectively, these results indicated that AMPARs were labelled with high selectivity after **CAM2-Ax647** administration into the brain.

To obtain structure–activity insights into the in-brain LD receptor labelling, we additionally prepared two compounds, **CAM2-Cy5** and **CAM2-SulfoCy5,** which have no and two negatively charged sulfonate groups in the Cy5 fluorophore, respectively, and another labelling reagent (**CAM2-Ax555**) with a Cy3-based fluorophore (which has a different emission wavelength) bearing four sulfonate groups. We then investigated the distribution properties of these compounds in the brain compared with **CAM2-Ax647**. Figure 3g shows that the unsulfonated **CAM2-Cy5** scarcely spreads from the injection site, while the more negatively charged **CAM2-SulfoCy5** (divalent anion) and **CAM2-Ax647** (tetravalent anion) are less adsorbed at the injection site and are widely diffused throughout the brain. In particular, **CAM2-Ax647** showed excellent diffusion, reaching areas of the Cb far from the injection site and had the highest signal-to-noise ratio. Similar high diffusibility was also observed for **CAM2-Ax555** (tetravalent anion), which showed almost the same labelling pattern as **CAM2-Ax647** (Supplementary Figure 2). These results indicated that the negative (net) charge of these reactive small molecules had a considerable impact on the diffusion in the live brain.

We subsequently conducted CUBIC-based tissue clearing^19^ of the whole brain after **CAM2-Ax647** LV injection. Light-sheet fluorescence microscopy (LSFM) showed stronger fluorescence signals in the regions of the hippocampus and cerebellar molecular layer, which were in good accordance with the high expression areas of AMPARs (Figure 3h, 3i, Supplementary Video 3)^17^. In CLSM imaging of this transparent brain sample, many small punctate signals (less than 1 μm in diameter), presumably originating from dendritic spines, were observed (Figure 3j). We also performed whole brain AMPAR labelling in the established 5xFAD transgenic mouse model of Alzheimer’s disease. The distribution of the labelled AMPARs detected in the wideview images was similar to that in normal mice (Figure 3k); however, labelled AMPARs were not observed around senile plaques, which are extracellular deposits of highly neurotoxic Aβ proteins, in the CA1 of the hippocampus area, probably because of a loss of excitatory neurons (Figure 3l).^20^ 3D analysis using a transparent brain sample showed that the density of AMPAR puncta in the regions containing senile plaques was less than in other areas, 0.295 ± 0.008 and 0.316 ± 0.005 spots/μm^3^, respectively (Figure 3m, Supplementary Video 4). These results demonstrated that our chemical labelling method enabled the visualization of the spatial distribution of endogenous AMPARs in the whole brains of genetically intact, transgenic, and disease model mice without requiring the preparation of numerous tissue slices. Notably, the receptor labelling was completed before fixation/clearing processes in our method, unlike antibody-based staining, which enabled snapshot images of the target endogenous receptors under more physiologically relevant conditions of the brain to be obtained.

### Acyl transfer-based chemical labelling of other receptors

Owing to the modular features of LDAI reagents, this chemistry can be readily extended for targeting other receptors by changing the ligand moiety. To demonstrate the robustness of our in-brain LD chemistry, we targeted several receptors with different structures and functions, namely, mGlu1,^21^ a class-C G protein-coupled (metabotropic glutamate) receptor; NMDAR,^22^ an ionotropic glutamate receptor involved in excitatory synaptic transmission together with AMPAR; GABA_A_R,^23^ a major inhibitory neurotransmitter receptor. Leveraging on the vast knowledge obtained from past drug discovery research on these receptors, we employed CNITM,^24^ L-689,560,^22^ and flumazenil^23^ as the ligand moieties for labelling reagents targeting mGlu1, NMDAR (NR1 subunit), and GABA_A_R, respectively (Figure 1b, Extended Data Figure 1). Although previously we have used gabazine and a benzodiazepine derivative as the ligand moiety in labelling reagents for GABA_A_R in model HEK293T cells,^13^ flumazenil was employed in the present live brain study because of its minimal toxicity. The corresponding ligand and the tetrasulfonated Ax647 were connected by an acyl imidazole group through a linker with an appropriate length as optimized in *in vitro* experiments (Supplementary Figure 3). We then injected these LDAI reagents into the LVs of a mouse brain and confirmed that there were no serious effects on the mouse behaviour. Note that a previously reported GABAAR-labelling reagent with a gabazine ligand (orthosteric antagonist) caused markedly intense seizures in mice, indicating the importance of appropriate ligand selection and dosage in in-brain LD chemistry. After mGlu1 labelling with **CmGlu1M**, the WB of the Cb homogenate exhibited three bands (300, 150, and 80 kDa) corresponding to aggregated, monomeric, and partially truncated mGlu1, respectively (Figure 4a). These bands were not detected in a mGlu1 knockout mouse,^25^ clearly indicating the specific labelling of mGlu1 in the brain (Figure 4a). Similarly, a strong band corresponding to the labelled NMDAR (110 kDa, NR1 subunit) and GABA_A_R (45 kDa, g2 subunit) was predominantly detected with **CNR1M** and **CGABAaRM**, respectively, while no obvious bands were detected with **NLC**. The IP-MS data also indicated the high target selectivity of these reagents as shown in Supplementary Data 2–4. In the CLSM imaging of the brain sections, fluorescence signals derived from the labelled receptors were observed from the particular brain regions where the corresponding endogenous receptors are reported to be abundantly expressed (Figure 4b). These fluorescence signals were well merged with those from immunostaining using anti-mGlu1,^26^ anti-NR1A,^27^ and anti-GABA_A_R-α1^28^ antibodies (Figure 4c, Extended Data Figure 2). The labelling was compatible with tissue clearing, which facilitated the 3D mapping of mGlu1, NMDAR, and GABA_A_R endogenously expressed in the brain (Supplementary Figure 4).

**Figure 4.**
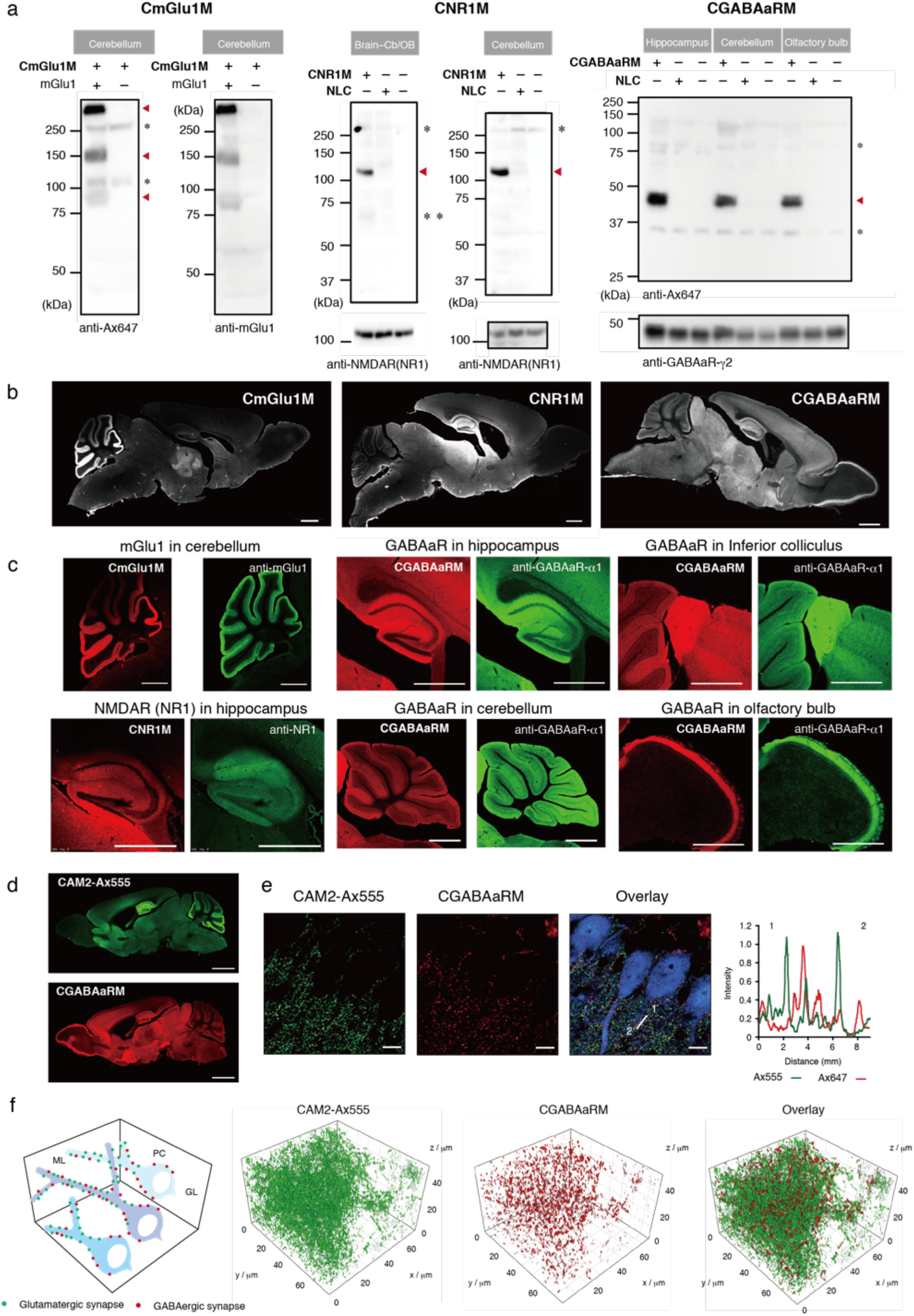
Chemical labelling of endogenous receptors (mGlu1, NMDAR, and GABA_A_R) in mouse brains. **a**, WB analysis of brain homogenates administered with **CmGlu1M**, **CNR1M**, **CGABAaRM** and **NLC**. Red triangle indicates specific labelling to a target receptor. * indicates non-specific bands derived from antibody. ** indicates non-specific labelling to serum albumin included in the brain. **b**, CLSM analysis of labelled whole brain slices. Left: **CmGlu1M** 100 μM × 4.5 μL, center: **CNR1M** 100 μM × 4.5 μL × 2, right: **CGABAaRM** 100 μM × 4.5 μL × 2. scale bar: 1 mm. **c**, Fluorescence images of sagittal sections of labelled brain with **CmGlu1M**, **CNR1M,** and **CGABAaRM**. Brain sections were coimmunostained with anti-mGlu1, anti-NR1 or anti-GABA_A_R-α1, respectively. Scale bar: 1 mm. **d**, Multiplex imaging of endogenous AMPAR and GABAaR by simultaneous injection of **CAM2-Ax555** and **CGABAaRM**. **CAM2-Ax555** (40 μM) and **CGABAaRM** (60 μM) dissolved in PBS(–) (4.5 μL) were simultaneously injected into LVs in both sides of mouse brain. 21 h after injection of labelling reagents, the mouse was transcardially perfused with 4% PFA/PBS(–). The imaging was performed using a CLSM equipped with a 5× objective. Scale bar: 2 mm. **e**, High-resolution confocal images of co-labelled cerebellum area with **CAM2-Ax555** and **CGABAaRM**, 63× objective, Zeiss Airy scan mode. Purukinje cells were stained with anti-calbindin (blue). Fluorescence intensities of Ax555 and Ax647 signals were analyzed by line plots. Scale bar: 10 μm. **f**, High-resolution 3D confocal images of cerebellum area cleared by CUBIC protocol after labelling with **CAM2-Ax555** and **CGABAaRM**, 63× lens, Zeiss Airy scan Z-stack mode.

We then sought to simultaneously label and visualize two different target receptors in an individual mouse. **CAM2-Ax555** and **CGABAaRM** were mixed and injected into the mouse brain. As shown in Figure 4d, strong fluorescence signals were observed from brain regions identical to the individual (single) labelling of AMPARs and GABA_A_Rs, indicating negligible interference between these reagents (Figure 4e). The high-resolution images of Cb regions exhibited a number of punctate signals derived from Ax555 or Ax647. These puncta were never overlapped with each other and were distinguishable in not only 2D, but also 3D images (Figure 4f). We counted these bright spots in the 1000 μm^3^ region near cerebellar Purkinje cells, which revealed that the 3D density of AMPAR puncta and GABAAR puncta are 0.751 ± 0.131 and 0.216 ± 0.069 /μm^3^, respectively (Figure 4f, Supplementary Video 5). Note that the value for the AMPAR density was similar to the density of excitatory synapses in the molecular layer of rat Cb (0.817 /μm^3^) previously determined by a stereological method using electron microscopy.^29^ In addition, our finding that AMPAR puncta exhibit a higher density than GABAAR puncta is consistent with previous reports that excitatory synapses are more abundant than inhibitory synapses.^30^ These results obtained by quantitative imaging analysis clearly highlight the power of our *in vivo* LD chemistry coupled with tissue clearing techniques.

### Labelling kinetics and lifetime studies of receptors in living mice brains

To apply our chemical labelling to more intricate biological experiments, determining how long it takes to complete the reaction and how long the labelled receptors can be followed in the living brain is important. We thus quantitatively characterized the reaction kinetics and the lifetimes of the labelled receptors (Figure 5a). After injection of **CAM2-Ax647**, followed by various incubation times prior to dissection, the homogenates of mice brains were subjected to SDS-PAGE in-gel fluorescence analysis. As shown in Figure 5b and Supplementary Figure 5a–e, the labelling band intensity increased for the initial 12 h and thereafter the band intensity decreased over several days. This biphasic profile can be fitted to a typical stepwise reaction comprising two distinct processes that presumably are the chemical labelling process (the 1st increasing step) and the following degradation of the labelled AMPARs (the 2nd decay step). Given that the CSF exchanges every 1.8 h in the mouse brain^31^, it is conceivable that most of the **CAM2-Ax647** was extruded a few hours after injection without interacting with AMPARs, halting further AMPAR labelling. Curve fitting analyses provided two kinetic values, 2.2 ± 0.6 h for the half-life of the first step (*T*_1/2, label_) and 97.2 ± 15.8 h for the second step (*T*_1/2, degradation_) in the Cb, and *T*_1/2, label_ and *T*_1/2, degradation_ values of 3.9 ± 1.9 and 92.6 ± 18.2 h, respectively, in the brain–Cb/olfactory bulb (OB). The labelling rates (*T*_1/2_ = 2–4 h) were sufficiently faster than the subsequent degradation rates (approximately 4 days). We observed similar biphasic kinetics for receptor labelling in HEK293T cells with a labelling rate almost comparable to that of in-brain labelling, i.e., 2.2–3.9 h in mice brains and 3.7 ± 1.4 h in HEK293T cells (Supplementary Figure 5e). Interestingly, these data imply that the labelling step (i.e., the acyl transfer reaction) is the rate-determining step rather than the inbrain diffusion of the LDAI reagent in the case of LV injection. Almost the same trends were observed for the other receptors (mGlu1, NMDAR, and GABA_A_R) in the brain and HEK cells as shown in Figure 5c and Supplementary Figure 5e. A signal decay curve of the labelled receptors gave lifetimes of ca. 40–95 h (*T*_1/2, degradation_), which is slightly shorter than the literature values determined by mass spectroscopy coupled with stable isotope labelling.^32,33^ Given that the LD chemistry relies on receptor-ligand recognition and our labelling reagents are cell impermeable, the obtained values may reflect the lifetime of the surface-exposed (functional) receptors under live brain conditions.

**Figure 5.**
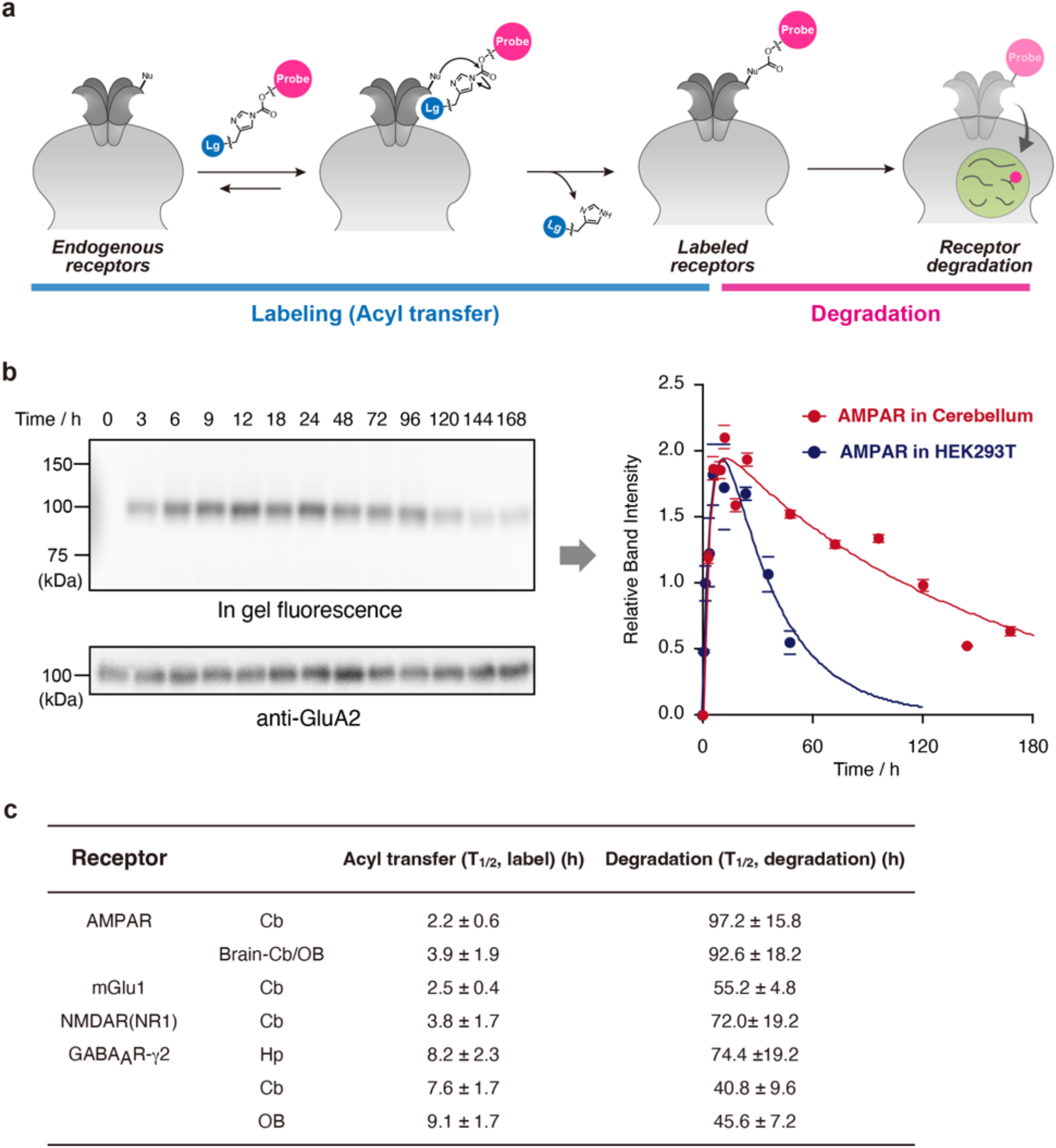
Time course analysis of acyl-transfer reaction of receptors and double chemical labelling of endogenous AMPAR and GABA_A_Rs. **a**, Schematic illustration of time course analysis of labelling reaction and labelled receptor degradation. **b**, Time course analysis of acyl-transfer reaction to AMPAR and degradation of labelled AMPAR. The band intensity of labelled receptor observed by in-gel fluorescence analysis were plotted with increasing the labelling reaction time. Data are presented as mean ± SEM. **c**, Kinetic parameters for acyl-transfer reaction to target receptors and degradation of labelled receptors. Data are presented as mean ± SEM.

### Chasing analysis of AMPAR dynamics in the mouse brain during development

We finally applied this labelling technique to *in vivo* pulse-chase analysis (Figure 6a). We investigated the dynamics of AMPARs during cerebellar development because synaptogenesis and synaptic pruning in the neonatal cerebellum have dynamically occurred,^34–37^ but the detailed molecular basis remains yet to be explored; for example, from where, how, and when are the synaptic proteins synthesized, degraded, and transported. Prior to the pulse-chase experiment, we confirmed that the selective labelling of cerebellar AMPARs in postnatal mice of various ages (at days P4, P7, and P18) could be performed by injection of **CAM2-Ax647** (Extended Data Figure 3a–f, Supplementary Videos 6–8). The snapshot imaging data showed that the Ax647-tagged AMPARs were localized in the areas surrounding PC soma in the P4 mouse, and the expression pattern changed from the soma to the dendrites of PCs from P7 to P18, which was consistent with previous reports using immunostaining (Extended Data Figure 3d–f).

**Figure 6.**
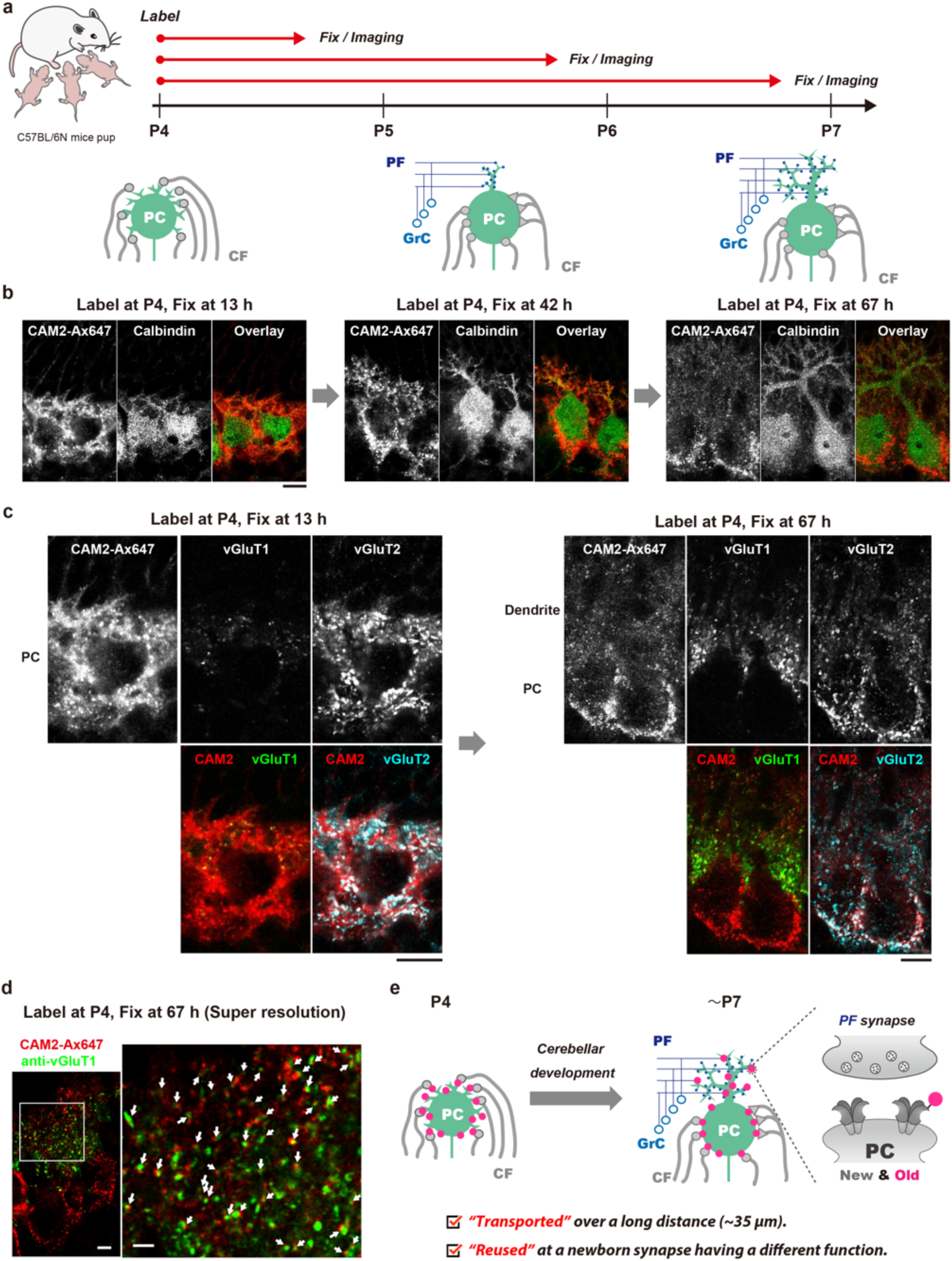
Pulse-chase analysis of AMPARs translocation during postnatal development. **a**, Schematic representation of pulse-chase experiments in this study. PC: Purkinje cells, CF: Climbing fibers, PF: Parallel fibers, GrC: Granule cells. Cerebellar PCs slowly develop dendrites and numerous synapses are formed on the dendrites. **b-c,** Pulse-chase analysis of P4 mice cerebellum labelled with **CAM2-Ax647**. PBS(–) containing 80 μM of **CAM2-Ax647** (2.0 μL) was injected into P4 mice brains. 13, 42, and 67 h after injection, the mice were transcardially perfused with 4% PFA. The brain was isolated and sectioned by cryostat (50-μm thick). The labelled sections were immunostained with anti-vGluT1, vGluT2, and calbindin. Imaging was performed using a CLSM equipped with a 100× objective. Scale bar: 10 μm. **d,** High-resolution fluorescence imaging of the labelled section at 67 hours after direct injection. Fluorescence image was acquired by using a CLSM equipped with a 100× objective and Lightning deconvolution. Colors: **CAM2-Ax647** (red) and anti-vGluT1 (green). Scale bars: 5 μm (left), 2 μm (right). **e,** A summary of the experimental results revealed in Figure 6.

We then proceeded to perform pulse-chase labelling in the neonatal mouse brain. Given the labelling kinetics and the lifetime of AMPARs as determined above, pulse-chase analysis in the live brain could conceivably be performed over a 3–4 day range with a dead time of ca. 12 h after injection (Figures 5 and 6a, Supplementary Figure 6). Thus, we injected **CAM2-Ax647** into P4 mice (pulse) and maintained the mice for 13 to 67 h (chase), followed by imaging of the cerebellar tissues. As shown in Figure 6b, the labelled AMPARs were located homogeneously around the soma of PCs at the P4 stage (chase at 13 h). Notably, at chase after 42 and 67 h, abundant fluorescence punctate signals were observed in the dendrites of PCs and the distribution of the fluorescence that remained in the soma after 13 h became heterogenous. A PC will receive inputs from CFs on soma in the early developmental stage and the PC will extend its dendrites toward the molecular layer to form new synapses with PFs in the distal area (PF synapses) during the period from P4 to P7 (Figure 6a). Given our findings obtained from the pulse-chase analysis and the established model of synaptogenesis in PCs, we suspected that some of the AMPARs expressed on the surface of PCs at P4 gradually translocate to the PF synapses in dendrites during functional neural circuit formation. To investigate this hypothesis, we conducted colocalization analysis by immunostaining with anti-vGluT1 and anti-vGluT2 antibodies, which are conventional PF- and CF-presynapse markers, respectively (Figure 6c, Extended Data Figures 4–6). Although vGluT2 is expressed in both CF and PF at the early developmental stage, CF synapses could be identified by their larger puncta size.^38^ High-resolution images of PCs showed that the labelled AMPAR signals were distributed over the cell surfaces, and some signals were observed from CF synapses at P4, while PF synapses (vGluT1-positive puncta) were barely detected at this time (Figure 6c, Extended Data Figure 4). At P7 (chase 67 h), numerous puncta from Ax647 signals along the dendrites were observed adjacent to the fluorescent spots derived from vGluT1, indicating that the nascent PF synapses in the dendrites contained AMPARs that were once present in the soma (Figure 6c and 6d, Extended Data Figures 5 and 6). Overall, these results validated our hypothesis and provided new insights into AMPAR dynamics during synaptogenesis, i.e., some early expressed (old) AMPARs in the developing mouse brain may be transported over a long distance (~35 μm) and become part of new synapses rather than a simple scrap-and-build scenario (Figure 6e). Although the physiological significance and mechanism of the distal transport of old AMPARs is beyond the scope of this study, the obtained results highlight the power of our *in vivo* pulse-chase labelling method using LDAI chemistry for the analysis of previously unknown behaviours of endogenous neurotransmitter receptors in their natural contexts.

## Discussion

Tremendous efforts in the chemical biology field have resulted in the development of a variety of sophisticated strategies for protein bioconjugation over the past few decades.^10,39–41^ However, there are as yet limited methods to achieve target-selective covalent protein modification *in vivo*. Although bioorthogonal chemistry has been shown to work *in vivo*, this method requires the incorporation of unnatural substrates into a protein for a chemoselective reaction.^42^ While the metabolic incorporation of such substrates has often been performed, this method does not show high selectivity for a particular protein.^43–45^ Although genetic code expansion technology allows the site-specific incorporation of noncanonical amino acids bearing a bioorthogonal reaction handle into a target protein in animals, the expression levels of the target protein are heavily suppressed because of the severe competition with engineered suppressor tRNAs with release factors for the stop codon.^46^ Activity-based probes have been used to modify native enzymes *in vivo* but while powerful for chemoproteomic research, this method generally results in the loss of the original activity of the enzyme and is therefore not suitable for functional analysis.^47,48^ In the present study, we have demonstrated LD chemistry that was bioorthogonal even under live brain conditions, enabling, for the first time, remarkably selective labelling of target neurotransmitter receptors in living mouse brains without genetic manipulation. The resultant whole brain imaging indicated that LD chemistry could be used to design labelling reagents with high targetability and good distribution features. Furthermore, we performed the kinetic analysis of a chemical labelling reaction in a live brain, which revealed that the affinity-based acyl transfer reaction proceeded with a half-life of a few hours. The present study should pave the way for *in vivo* organic chemistry methods targeting various biofunctional molecules. Furthermore, given that many synthetic chemical biology probes are now available, we envisioned that a target receptor may be directly functionalized with such probes in the brain for advanced studies. Our ligand-directed approach is simple and compatible with the conventional genetic and chemogenetic labelling methods, leading to the potential for a rational combination of methods for multiplex analysis, such as multi-colour imaging and the opto/chemogenetic functional regulation of live animals.^9,49^ Such efforts are expected to contribute to elucidating the functions and dynamics of neurotransmitter receptors in the complicated neural circuits of the live brain in detail.

## Methods

### Synthesis

All synthesis procedures and characterizations are described in the Supplementary Information.

### Animal experiments

C57BL6/N mice were purchased from Japan SLC, Inc (Shizuoka, Japan). The animals were housed in a controlled environment (23 °C, 12 h light/dark cycle) and had free access to food and water, according to the regulations of the Guidance for Proper Conduct of Animal Experiments by the Ministry of Education, Culture, Sports, Science, and Technology of Japan. All experimental procedures were performed in accordance with the National Institute of Health Guide for the Care and Use of Laboratory Animals, and were approved by the Institutional Animal Use Committees of Kyoto University and Keio University.

Experimental details for injection of the labeling reagents, sample preparation, electrophysiology, and fluorescence imaging are described in the Supplementary Information.

## Acknowledgments

The authors thank Dr. Hideki Nakamura for discussions and Dr. Muneo Tsujikawa, Ms. Kumiko Nishizawa, and Ms. Tomoko Gonda for technical support of biological experiments. The authors also appreciate the support of The IRCN Imaging Core, The University of Tokyo Institutes for Advanced Studies, for the usage of the LSFM system developed by Masafumi Kuroda (The University of Tokyo). The authors also thank Victoria Muir, PhD, from Edanz (https://jp.edanz.com/ac) for editing a draft of this manuscript. This work was supported by the Japan Science and Technology Agency (JST) ERATO (grant number JPMJER1802 to I.H. and JPMJER2001 to H.R.U.), the Science and Technology Platform Program for Advanced Biological Medicine (AMED/MEXT, grant number JP22am0401006 to H.N., grant number JP22am0401011 to H.R.U.), MEXT/JSPS KAKENHI Grant-in-Aid for JSPS Fellows (grant number 21J15773 to K.S.), Grant-in-Aid for Scientific Research on Innovative Areas “Integrated Bio-metal Science” (grant number 19H05764 to T.T.), JSPS KAKENHI grant-in-aid for scientific research (S) (grant number JP18H05270 to H.R.U.), MEXT Quantum Leap Flagship Program (MEXT QLEAP) (grant number JPMXS0120330644 to H.R.U.,), JSPS KAKENHI grant-in-aid for scientific research (B) (grant number 22H02824 to E.A.S.), AMED-PRIME (grant number JP22gm6210027 to E.A.S.), Grants-in-Aid from the Takeda Science Foundation (to E.A.S., W.K., and I.A.), Nakatani foundation for advancement of measuring technologies in biomedical engineering (to E.A.S.), and Mochida Memorial Foundation for Medical and Pharmaceutical Research (to E.A.S.).

## Data availability

The authors declare that the data supporting the findings of this study are available with the paper and its Supplementary information files. The data that support the findings of this study are available from the corresponding author upon reasonable request.

## Contributions

H. N. and I.H. initiated and designed the project. H.N., S.S., K.S., M.I., T.T., K.O., and S.K. performed synthesis and chemical labeling experiments in HEK293T cells and brains with the help of W.K. and M.Y. W.K., I.A., and M.Y. performed electrophysiological experiments. E.A.S., C.S., and H.R.U. performed LSFM experiments of CUBIC-cleared brains. H.N., S.S, K.S., T.T., S.K., W.K., I.A., and I. H. prepared the manuscript with contributions from the other authors.

## Corresponding author

Correspondence to Itaru Hamachi.

---

**Extended Data Figure 1.**
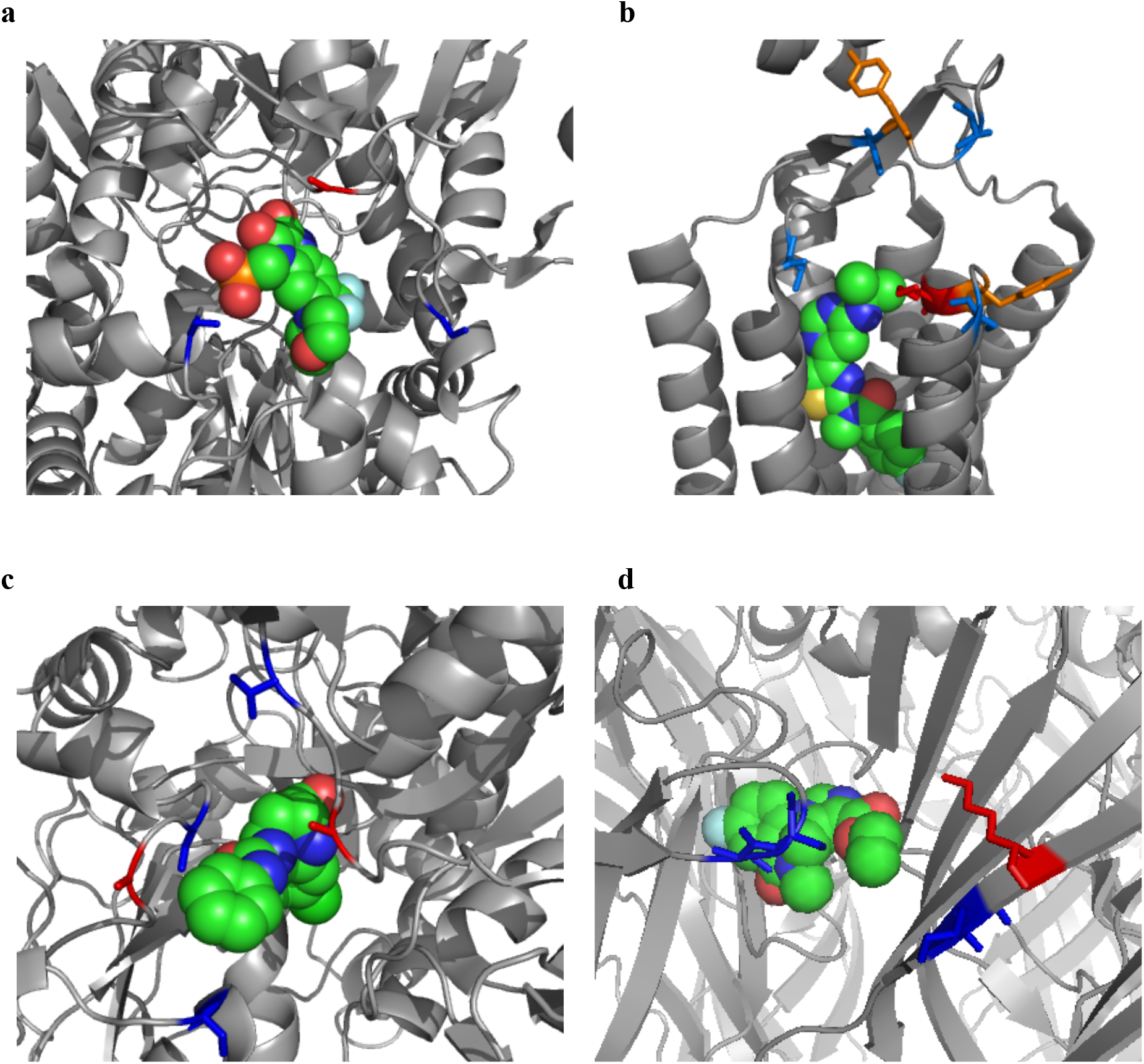
Design of new labelling reagents for AMPAR, mGlu1, NMDAR, and GABAaR. The labelling reagents were designed based on the information of these co-crystal structures. Space-filling drawing of ligands, ZK200775 (PFQX analogue) (a) FITM (CNITM analog) (b), L689,560 (c) and Flumazenil (d) bound to AMPAR, mGlu1, NMDAR (NR1) and GABAaR, respectively. Side chains of nucleophilic amino acid residues are drawn as stick. Red = Lys, Blue = Ser or Thr, Orange = Tyr. PDB ID = 3KG2 (AMPAR), 4OR2 (mGlu1), 6WHY (NR1), 6D6U (GABAaR).

**Extended Data Figure 2.**
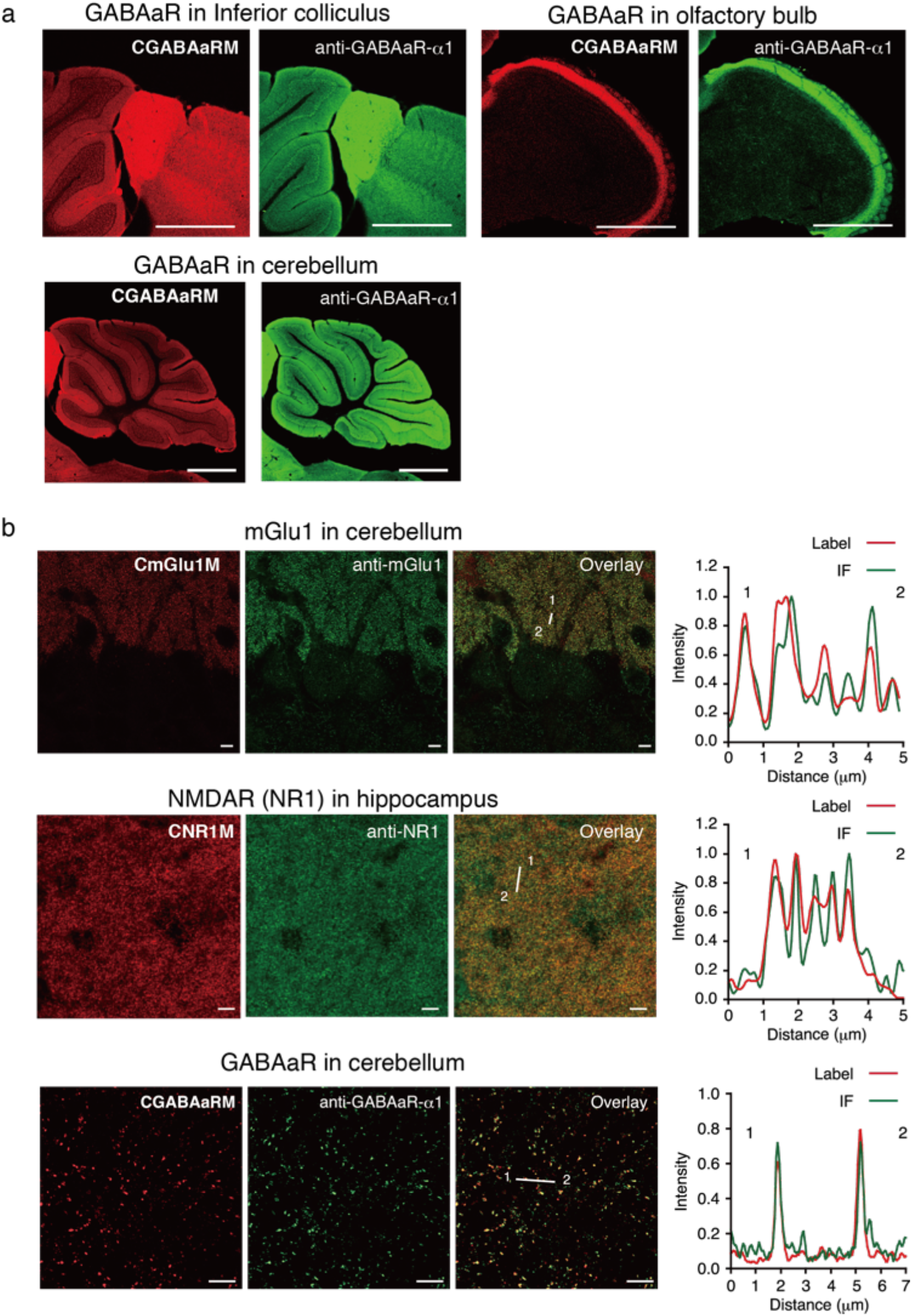
CLSM images of immunostained mouse brain slices labelled with CmGlu1M, CNR1M, and CGABAaRM. Brain slices were prepared as described in Figure 4c. **a**, Fluorescence images of sagittal sections of labelled brain with **CGABAaRM**. Brain sections were coimmunostained with anti-GABAAR-a1. Scale bar: 1 mm. **b**, High-resolution confocal images of cerebellum and hippocampus area labelled with **CmGlu1M** (100× lens, Leica Lightning deconvolution), **CNR1M** (100× lens, Leica Lightning deconvolution) and l **CGABAaRM** (63× lens, Zeiss Airy scan). Fluorescence intensities of labelling and immunostaining were analyzed by line plots. Scale bar: 5 μm.

**Extended Data Figure 3.**
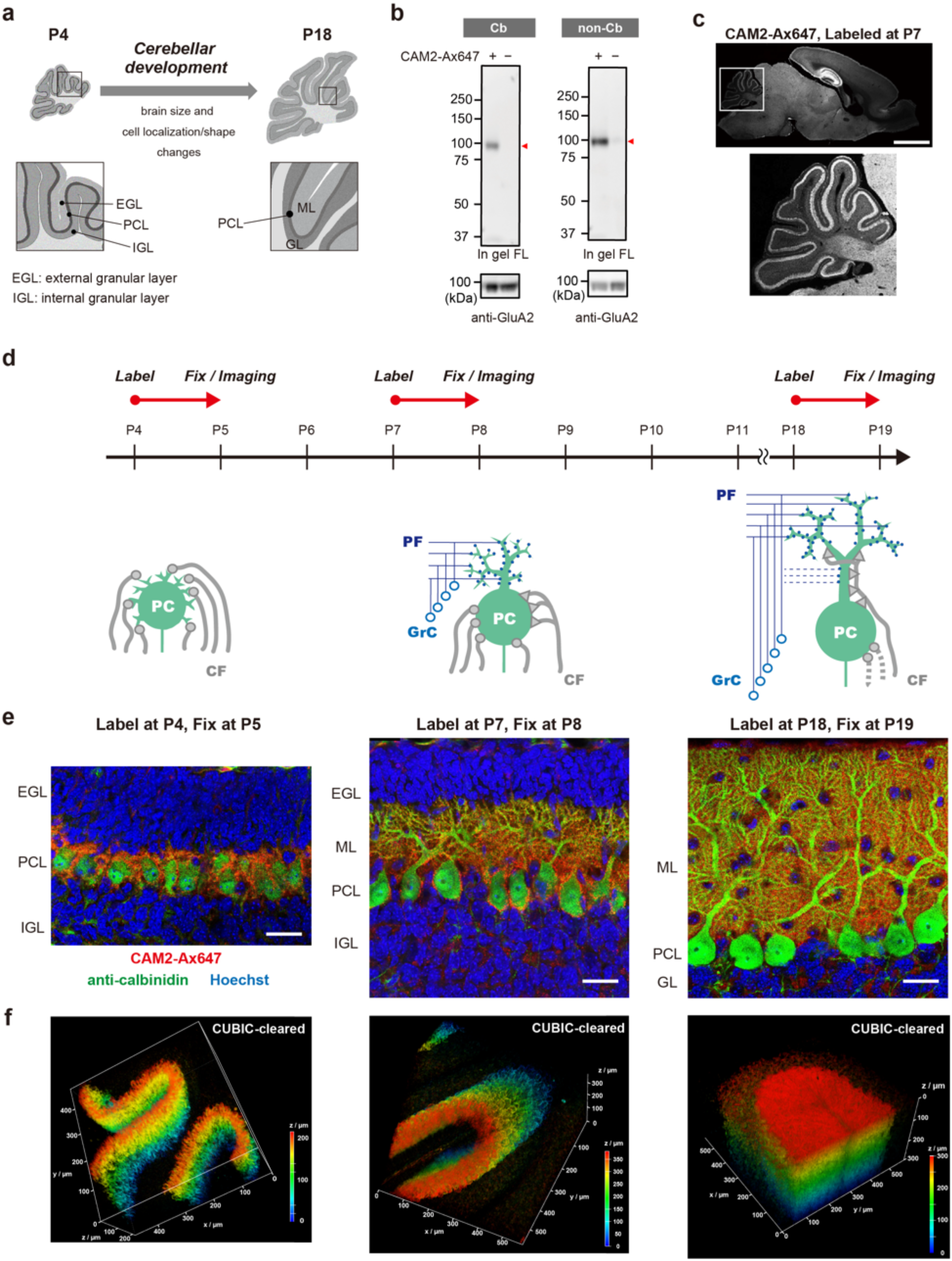
Chemical labelling of endogenous AMPARs in neonatal mouse cerebellum. **a**, Schematic illustration of cerebellar development. The developing process of Cb involves proliferation of granule cells in the external granular layer (EGL) and subsequent cell migration into the internal granular layer (IGL). **b**, In gel fluorescence analysis of brain homogenates administered with **CAM2-Ax647** (80 μM, 2.0 μL) to P7 mouse brain. Red triangle indicates specific labelling to a target receptor. **c**, Fluorescence image of a sagittal brain section labelled with **CAM2-Ax647** at P7. 24 h after injection with **CAM2-Ax647** (80 μM, 2.0 μL), the mouse was transcardially perfused with 4% PFA. The brain was isolated and sectioned by cryostat (50-μm thick). Imaging was performed using a CLSM equipped with a 5× objective (633 nm excitation for Ax647). Scale bar: 2 mm. **d,** Schematic illustration of Purkinje cells during cerebellar development and snapshot-type labelling experiments in this Figure. PC: Purkinje cells, CF: Climbing fibers, PF: Parallel fibers, GrC: Granule cells. Cerebellar PCs slowly develop dendrites over several postnatal weeks, and numerous synapses are formed on the dendrites. **e,** Snapshot analysis of post-natal mice cerebellum labelled with **CAM2-Ax647** (80 μM, 2 μL, PBS(–)) by CLSM. Colors: **CAM2-Ax647** (red), anti-calbindin (green), Hoechst (blue). Fluorescence imaging using a CLSM equipped with a 100× objective. Scale bar: 25 μm. **f,** 3D fluorescence imaging of a tissue-cleared cerebellum in Extended Data Figure 3e. Fluorescence imaging using a CLSM equipped with a 20× objective.

**Extended Data Figure 4.**
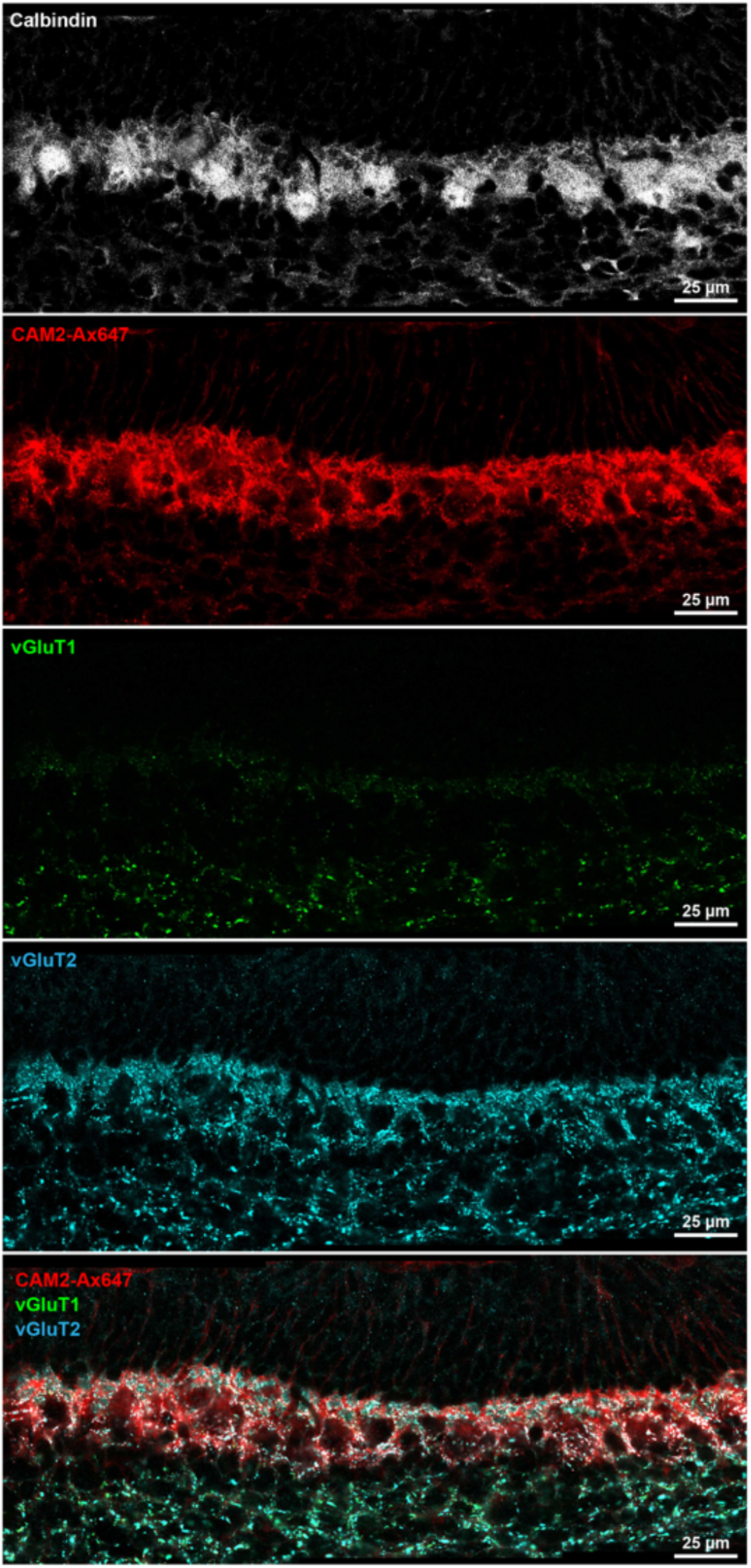
Fluorescence images around cerebellar Purkinje cells at 13 hours after injection. P4 mouse was labelled with **CAM2-Ax647** (80 μM, 2 μL, PBS(–)). The mouse was transcardially perfused with cold 4% PFA/PBS(–) at 13 h after injection. After cryosection of the isolated brain, immunostaining was conducted with anti-vGluT1 (Frontier institute, VGluT1-Rb-Af500, rabbit), anti-vGluT2 (Frontier institute, VGluT2-Gp-Af810, guinea pig), and anti-calbindin (abcam, mouse, ab82812). Fluorescence image was obtained with a CLSM equipped with a 100× objective. Colors: **CAM2-Ax647** (red), anti-vGluT1 (green), anti-vGluT2 (cyan), and anti-calbindin (gray).

**Extended Data Figure 5.**
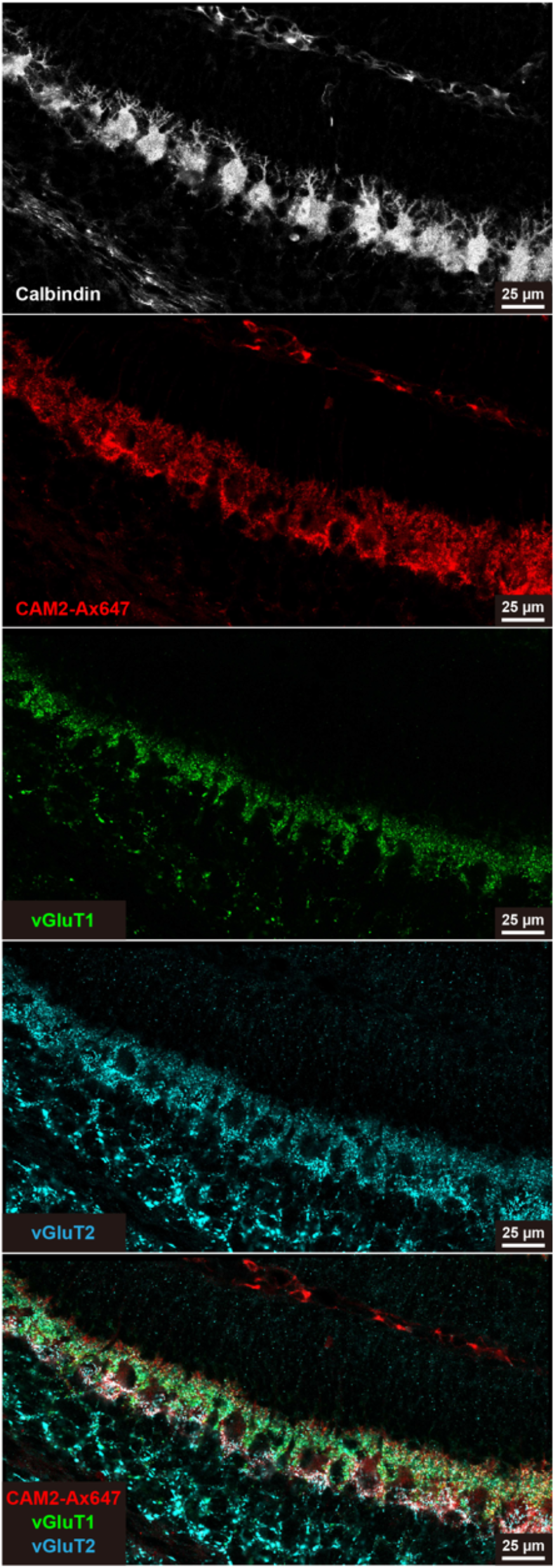
Fluorescence images around cerebellar Purkinje cells at 42 hours after injection. P4 mouse was labelled with **CAM2-Ax647** (80 μM, 2 μL, PBS(–)). The mouse was transcardially perfused with cold 4% PFA/PBS(–) at 42 h after injection. All procedures were same as the sample at 13 h after injection.

**Extended Data Figure 6.**
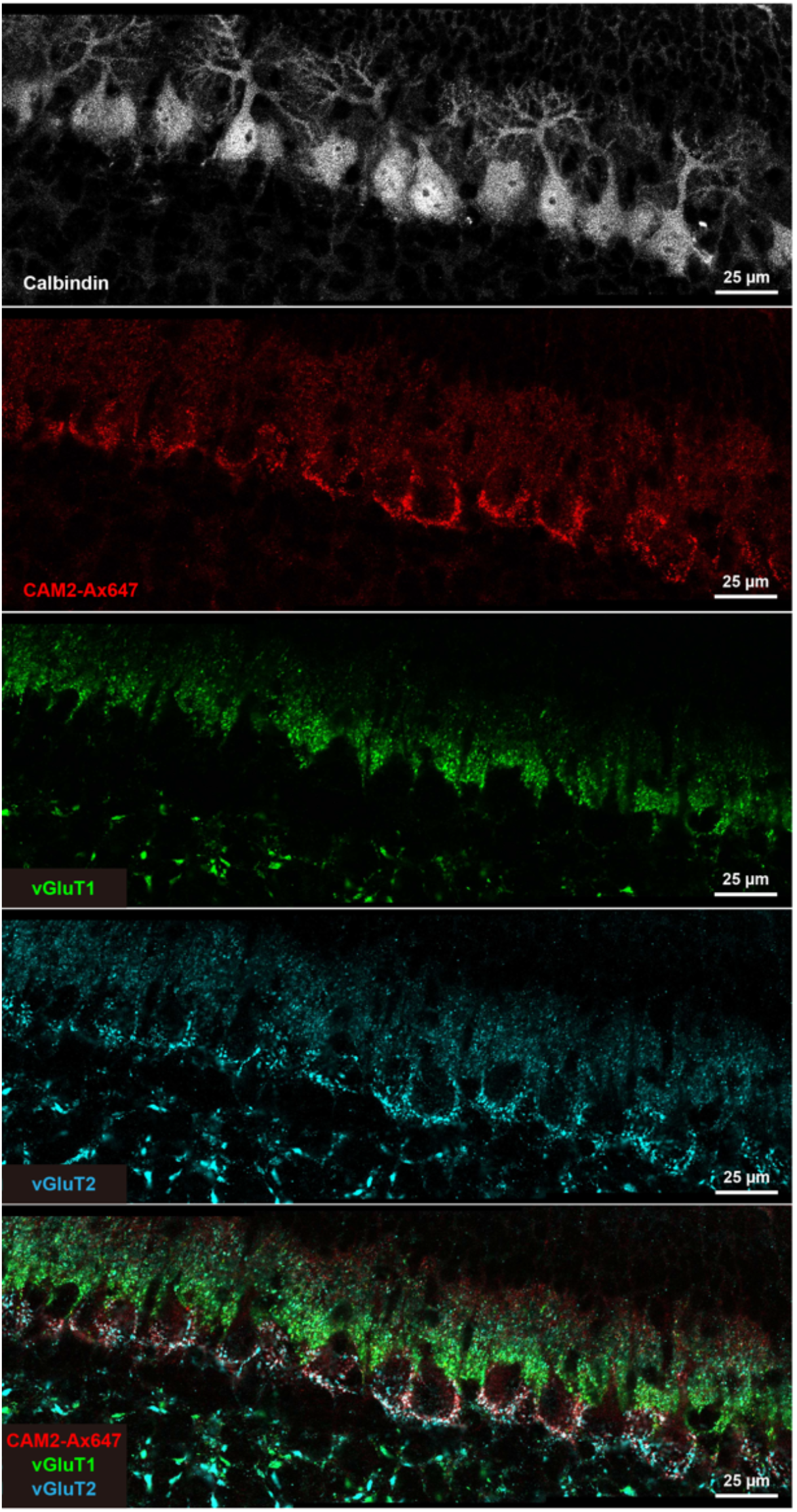
Fluorescence images around cerebellar Purkinje cells at 67 hours after injection. P4 mouse was labelled with **CAM2-Ax647** (80 μM, 2 μL, PBS(–)). The mouse was transcardially perfused with cold 4% PFA/PBS(–) at 67 h after injection. All procedures were same as the sample at 13 h after injection.

