## Supplementary Information for "Bioorthogonal chemical labelling of endogenous neurotransmitter receptors in living mouse brains"

### Supplementary Figures

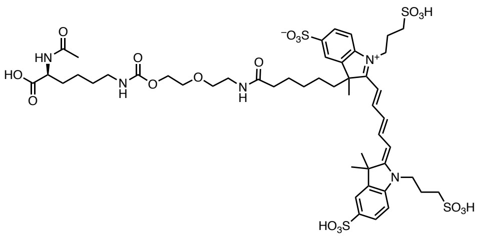

**Lys-labeled product**

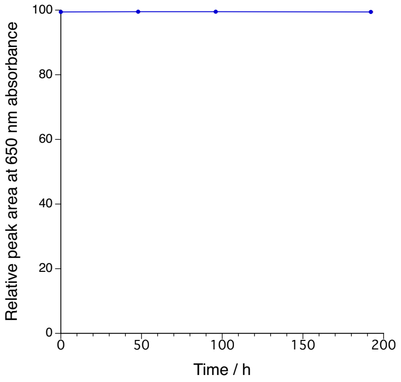


**Supplementary Figure 1 | Stability of the covalent bond formed between LDAI labelling reagents and Lys side chains.** The solution of **Lys-labeled product** (1 μM) was prepared with Neurobasal Medium (Gibco™) that meets the special cell culture requirements of pre-natal and embryonic neuronal cells. After 48, 96, and 192 h incubation at 37ºC, the mixture was analyzed by RP-HPLC. RP-HPLC analysis was conducted on a Shimadzu Nexera system equipped with a fluorescent detector (640/660 nm (Ex/Em)) with a linear gradient of 0-45% $CH_3CN$/10 mM $NH_4OAc$ aq. (45 min). Data are presented as mean ± SEM.

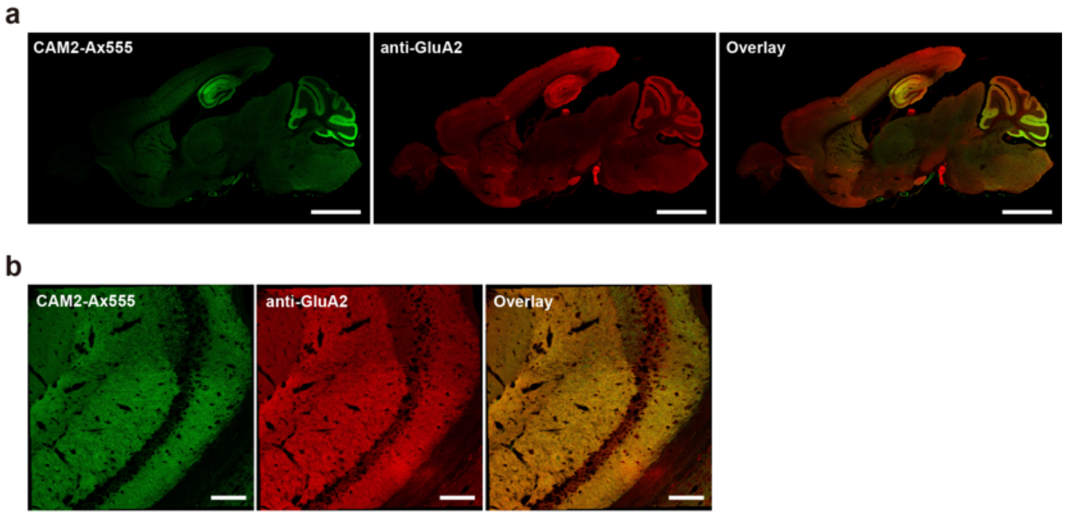


**Supplementary Figure 2 | Fluorescence images of labelled brains with CAM2-Ax555.** PBS(−) containing 100 μM of **CAM2-Ax555** (4.5 μL, $C_{brain}$ = ca. 1 μM ) was injected into the lateral ventricle (LV) of mice. 24 h after injection, the mouse was transcardially perfused with 4% PFA/PBS(−). The brain was isolated and sectioned by cryostat (20-μm thick). After immunostaining with anti-GluA2 (1:500, MAB397, Merk) and Anti-Ms IgG Ax647 (1:200, ab150115, abcam), imaging was performed using a CLSM equipped with GaAsP detector (561 nm excitation for Ax555 and 633 nm excitation for Ax647. (a) 5× objective, Scale bar: 2 mm. (b) 10× objective, Scale bar: 100 μm. The fluorescence images of labelled brain were well merged with these of Anti-GluA2.

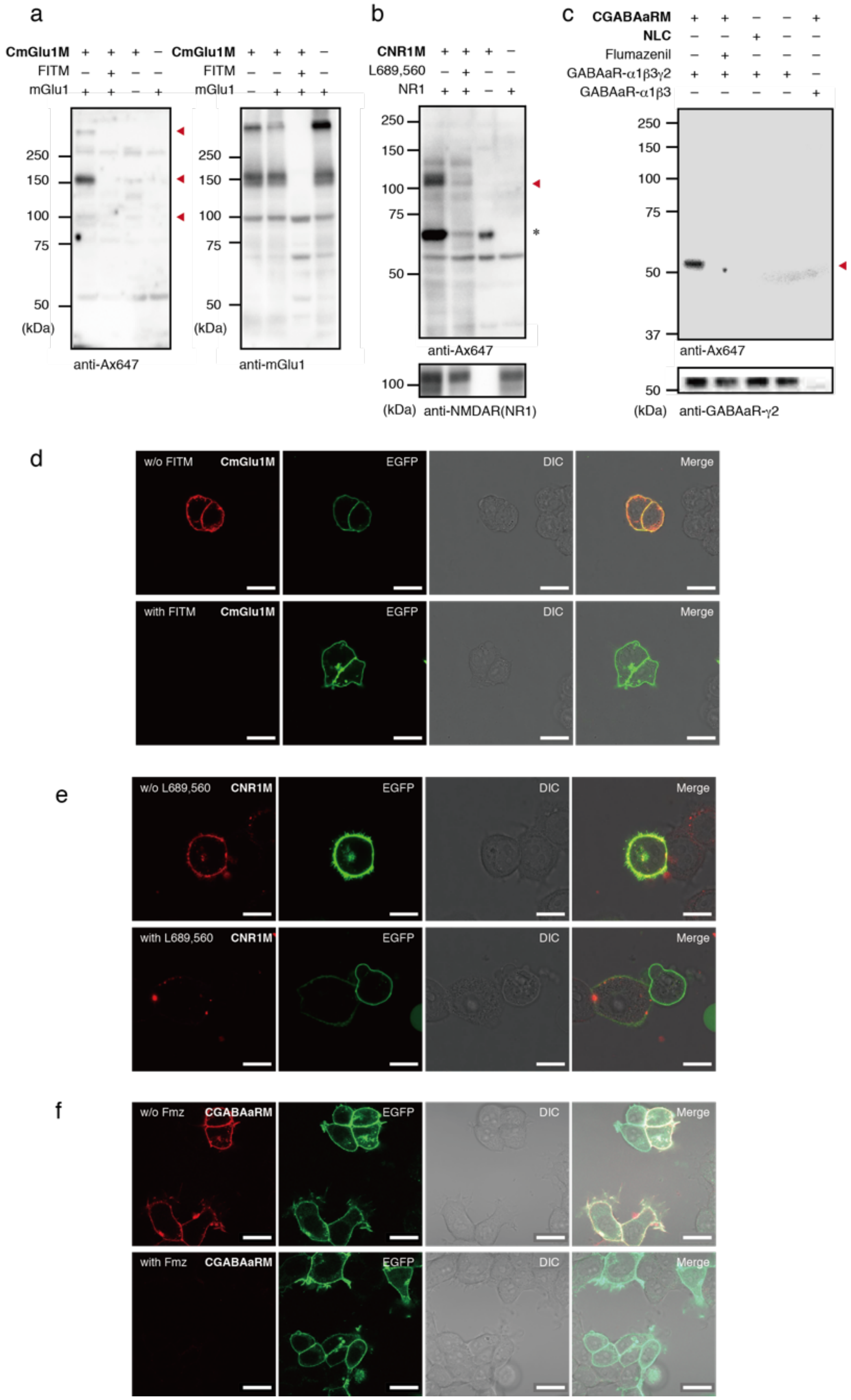
a
CmGlu1M + + + −
FITM − + − −
mGlu1 + + − +
250
150
100
75
50
(kDa)
anti-Ax647
CmGlu1M + + + −
FITM − − + −
mGlu1 − + + +
anti-mGlu1
b
CNR1M + + + −
L689,560 − + − −
NR1 + + − +
anti-Ax647
anti-NMDAR(NR1)
c
CGABAaRM + + − − +
NLC − − + − −
Flumazenil − + − − −
GABAaR-α1β3γ2 + + + + −
GABAaR-α1β3 − − − − +
37
anti-GABAaR-γ2
d
w/o FITM
with FITM
CmGlu1M
EGFP
DIC
Merge
e
w/o L689,560
with L689,560
CNR1M
f
w/o Fmz
with Fmz
CGABAaRM

**Supplementary Figure 3 | Chemical labelling of mGlu1, NMDAR, and GABA$_A$R expressed in HEK293T cells.** a–c, Western blot (WB) analysis of HEK293T cells labelled with **CmGlu1M** (a), **CNR1M** (b), **CGABAaRM**, and **NLC** (c). HEK293T cells transfected with mGlu1, NMDAR (NR1: a subunit of NMDAR), GABA$_A$R-α1β3γ2 (a major subtype in brain synapse possessing a benzodiazepine site), GABA(A)- α1β3, or vector control were treated with 2 μM **CmGlu1M**, **CNR1M**, **CGABAaRM**, or **NLC** in the presence or absence of each competitive inhibitor (FITM, L689,560 or flumazenil, 100 μM) in serum-free at 17°C for 4 h. The cell lysates were analyzed by western blot using anti-Ax647, anti-GluR1, anti-NR1 or anti GABAAR-γ2 antibody. ▲ and * indicate specific labelling and non-specific labelling to bovine serum albumin included in culture medium, respectively. d–f, Confocal live imaging of HEK293T cells labelled with **CmGlu1M** (d), **CNR1M** (e), or **CGABAaRM** (f). Chemical labelling was conducted as described in a–c. EGFP-F was used as a transfection marker.

a

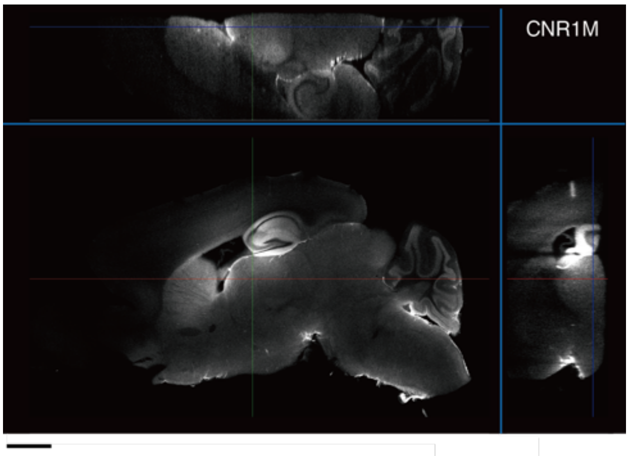


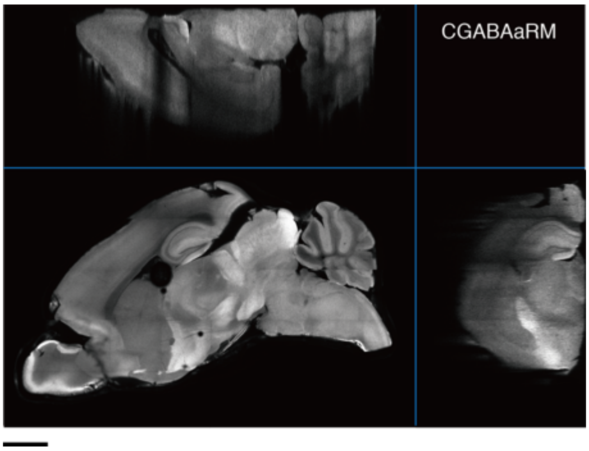


b

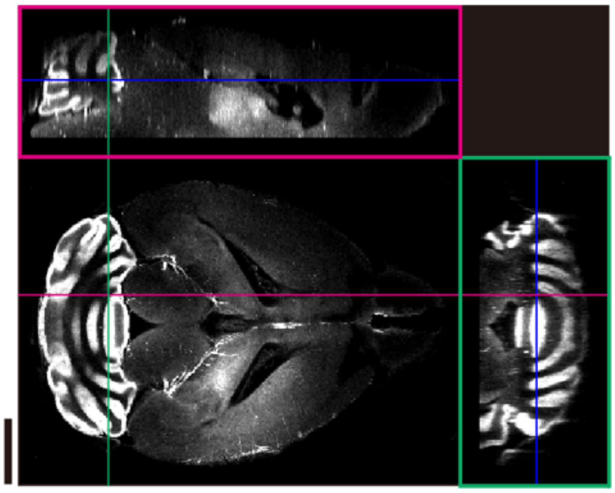

**Supplementary Figure 4 |** a, 3D rendering of the right-side brain cleared by CUBIC protocols after labelling with **CNR1M** and **CGABAaRM**. b, 3D rendering of the whole brain cleared by

3DISCO protocols after labelling with **CmGlu1M**. Scale bar: 2 mm. While the CUBIC protocols strongly attenuated the fluorescence signals of mGlu1 labelled with **CmGlu1M,** probably due to the cleavage of chemical bond formed by the acyl transfer reaction under the basic conditions of CUBIC protocols, the 3DISCO method allowed to prepare the transparent brain tissues fluorescently labelled with **CmGlu1M** and showed the well-matched distribution of native mGlu1.

a

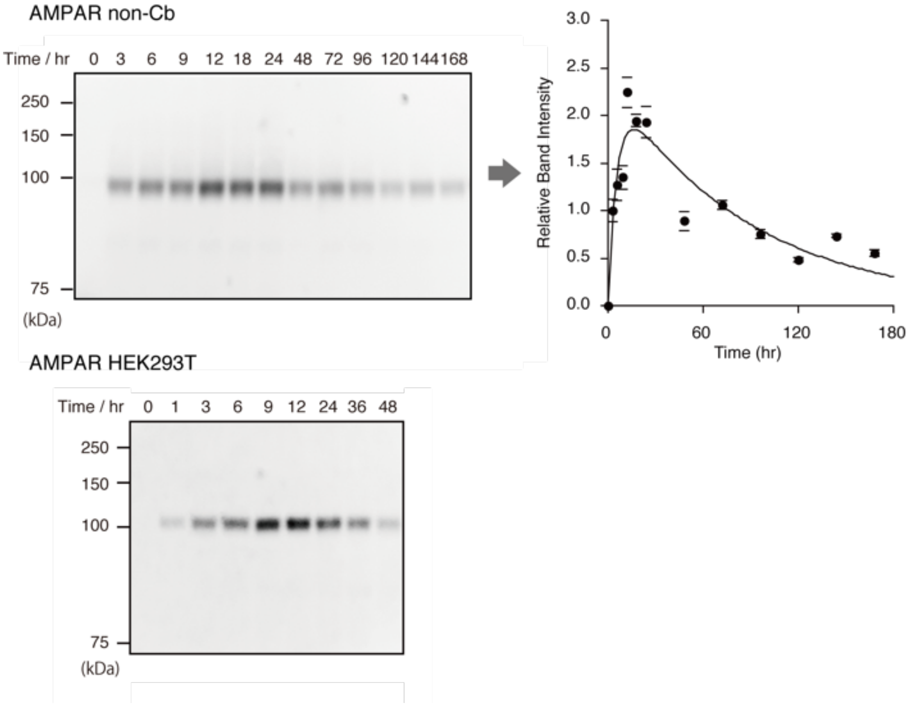
AMPAR non-Cb
Time / hr 0 3 6 9 12 18 24 48 72 96 120 144 168
250
150
100
75
(kDa)
Relative Band Intensity
Time (hr)
AMPAR HEK293T
Time / hr 0 1 3 6 9 12 24 36 48
250
150
100
75
(kDa)

b

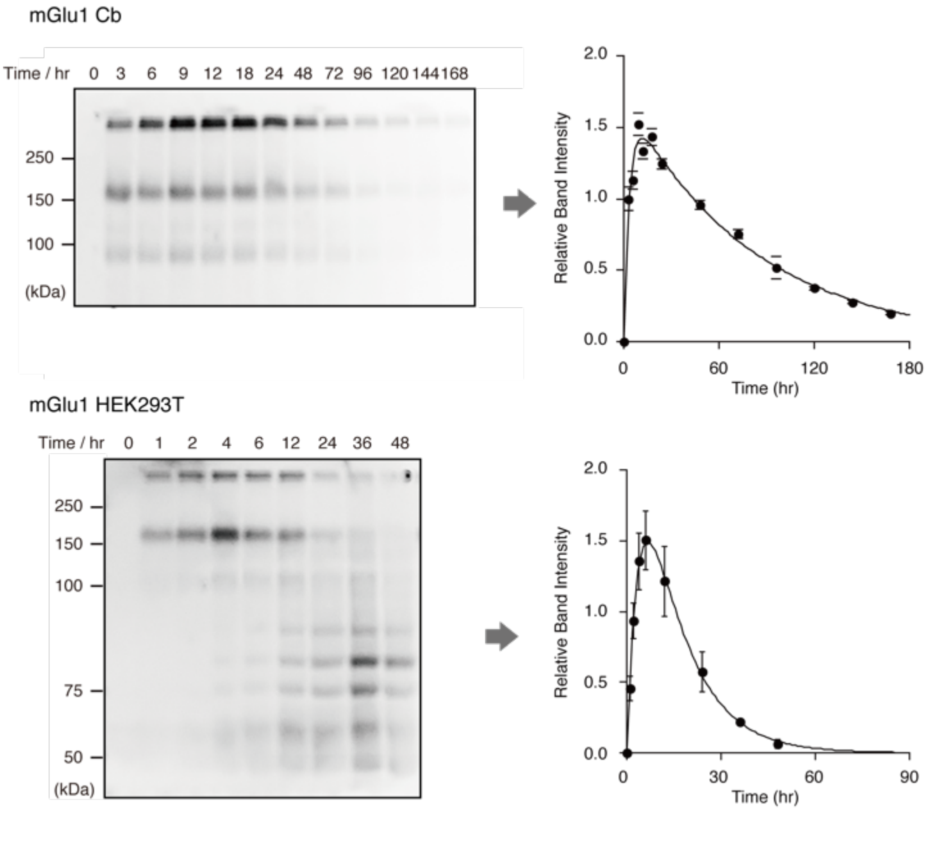
mGlu1 Cb
Time / hr 0 3 6 9 12 18 24 48 72 96 120 144 168
250
150
100
(kDa)
Relative Band Intensity
Time (hr)
mGlu1 HEK293T
Time / hr 0 1 2 4 6 12 24 36 48
250
150
100
75
50
(kDa)
Relative Band Intensity
Time (hr)

**c**

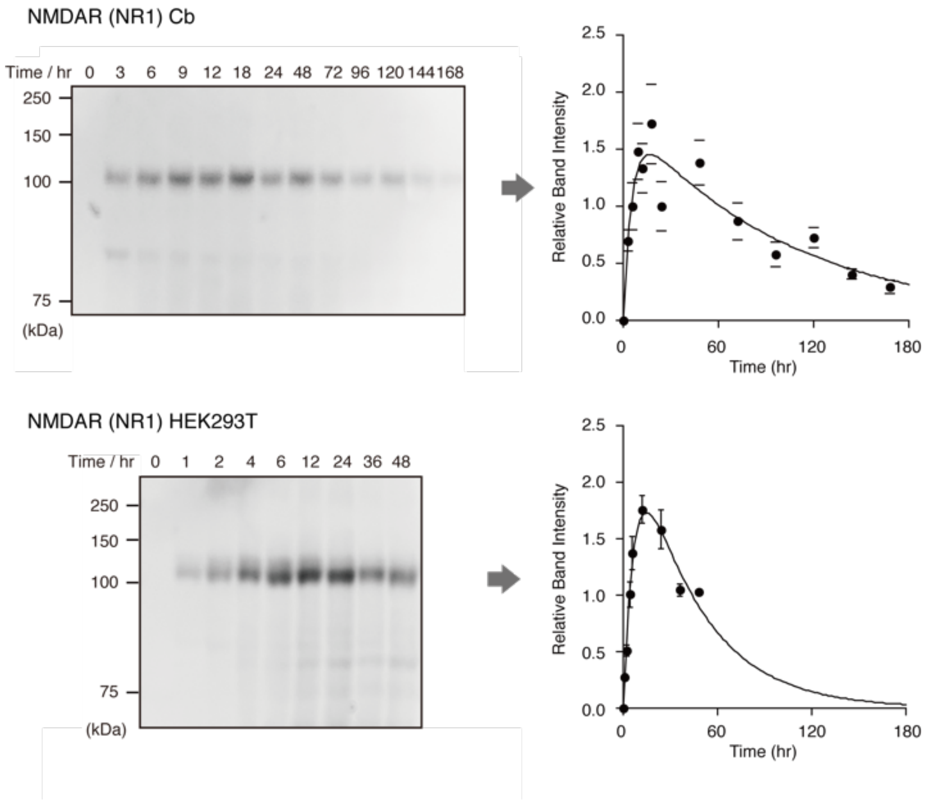
NMDAR (NR1) Cb
Time / hr 0 3 6 9 12 18 24 48 72 96 120 144 168
250
150
100
75
(kDa)
Relative Band Intensity
Time (hr)
NMDAR (NR1) HEK293T
Time / hr 0 1 2 4 6 12 24 36 48
250
150
100
75
(kDa)
Relative Band Intensity
Time (hr)

**d**

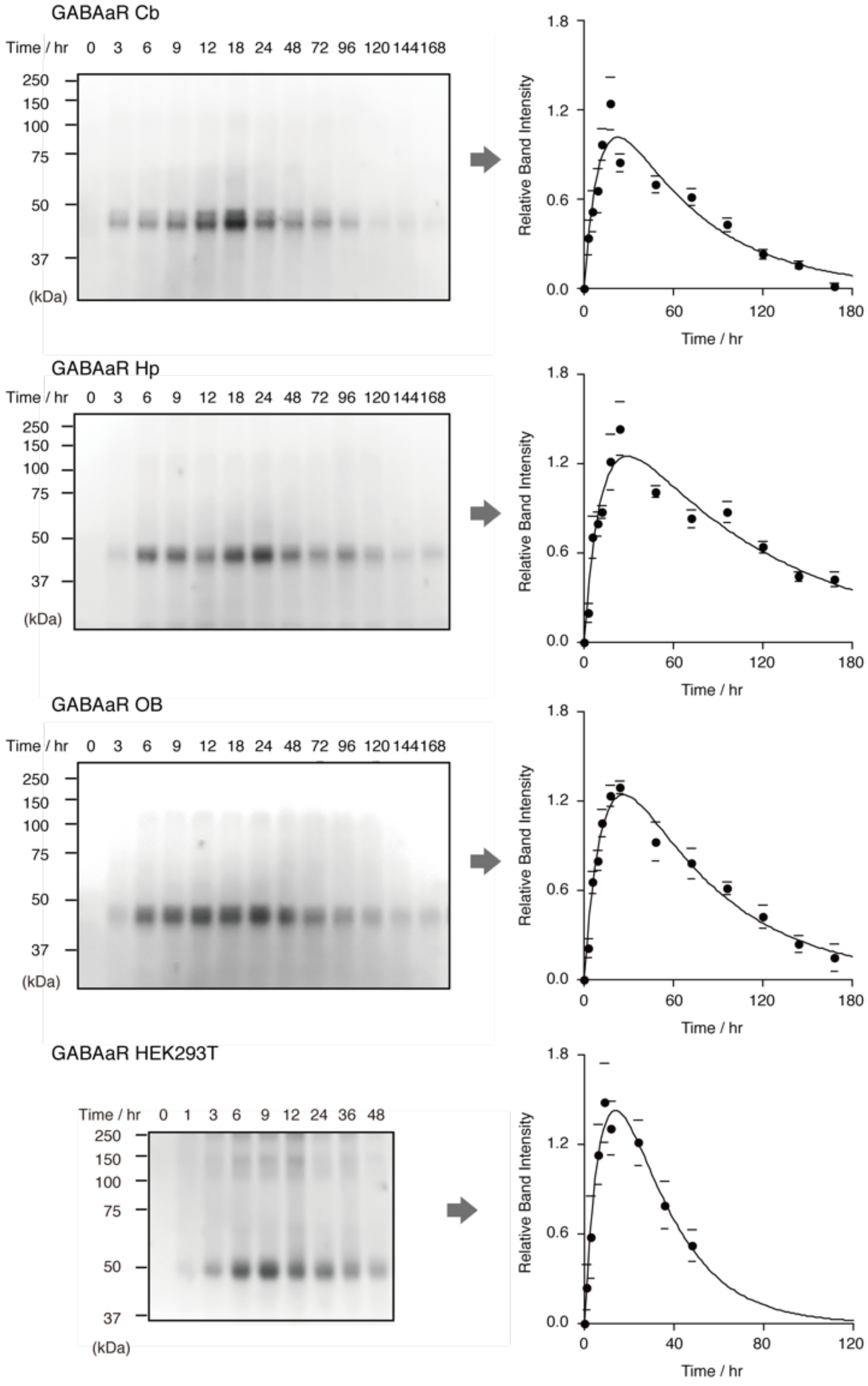
GABAaR Cb
Time / hr 0 3 6 9 12 18 24 48 72 96 120 144 168
250
150
100
75
50
37
(kDa)
Relative Band Intensity
Time / hr
GABAaR Hp
Time / hr 0 3 6 9 12 18 24 48 72 96 120 144 168
GABAaR OB
Time / hr 0 3 6 9 12 18 24 48 72 96 120 144 168
GABAaR HEK293T
Time / hr 0 1 3 6 9 12 24 36 48

**e**

| Receptor | | Acyl transfer ($T_{1/2}$, label) (h) | Degradation ($T_{1/2}$, degradation) (h) |
|---|---|---|---|
| AMPAR | HEK293T | 3.7 ± 1.4 | 20.6 ± 7.4 |
| mGlu1 | HEK293T | 2.8 ± 0.3 | 8.7 ± 0.7 |
| NMDAR(NR1) | HEK293T | 4.9 ± 0.8 | 27.4 ± 5.6 |
| $GABA_AR$-γ2 | HEK293T | 6.4 ± 3.9 | 15.4 ± 9.4 |

**Supplementary Figure 5 | Time course analysis of acyl-transfer reaction to endogenous AMPAR, mGlu1, NMDAR (NR1), and $GABA_A$ receptors and degradation of labelled receptors in the mouse brain.** In gel fluorescence analysis and plots of band intensity versus time in cerebellum (Cb), brain-Cb/olfactory bulb (OB), and HEK293T cells labelled with **CAM2-Ax647** (a), in Cb labelled with **CmGlu1M** (b), in Cb **CNR1M** (c), in hippocampus, Cb, OB, and HEK293T cells labelled with **CGABAaRM** (d). (e) Kinetic parameters for acyl-transfer reaction to target receptors and degradation of labelled receptors. The biphasic processes were fitted by a theoretical equation. Data are presented as mean ± SEM.

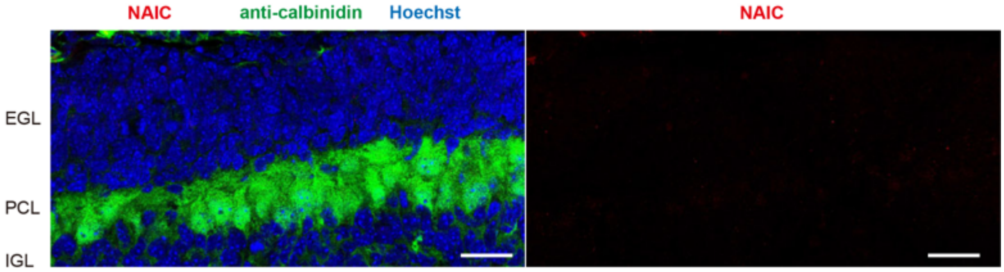


**Supplementary Figure 6 | Fluorescence images of sagittal Cb sections injected with NAIC at postnatal day 4 (P4).** PBS(−) containing 80 µM of **NAIC** (2.0 µL) was injected to the mouse brain at P4. 12 h after the injection, the mouse was transcardially perfused with 4% PFA/PBS(−). The brain was isolated and sectioned by cryostat (50-µm thick). The slices were permeabilized and immunostained with primary antibody anti-calbindin. After that, secondary antibody (Ax488) and Hoechst 33258 staining was conducted in 0.1% Triton X-100/PBS(−). Fluorescence image was acquired by using a CLSM equipped with a 100× objective and GaAsP detector (405 nm excitation for Hoechst, 488 nm excitation for Ax488 and 647 nm excitation for Ax647). Colors: **CAM2-Ax647** (red), anti-calbindin (green), and Hoechst 33258 (blue). Scale bar: 25 µm.

### Supplementary Methods

**General methods for biochemical and biological experiments.**

SDS-PAGE and western blotting (WB) were carried out using a BIO-RAD Mini-Protean III electrophoresis apparatus. Samples were applied to SDS-PAGE and electrotransferred onto polyvinylidene fluoride membranes (BIO-RAD), followed by blocking with 5% nonfat dry milk in Tris-buffered saline containing 0.05% Tween 20. Primary antibody was indicated in each experimental procedure, and anti-rabbit IgG-HRP conjugate (CST, 7074S, 1:3,000) or anti-mouse IgG-HRP conjugate (CST, 7076S, 1:3,000), and anti-sheep IgG-HRP conjugate (abcam, ab6747-1, 1:2000) was utilized as the secondary antibody. Chemiluminescent signals generated with ECL Prime (GE Healthcare) were detected with a Fusion Solo S imaging system (Vilber Lourmat).

**General information for fluorescence imaging experiments.**

Fluorescence imaging was performed using a CLSM (Leica microsystems, Germany, TCS SP-8 or Carl Zeiss, Germany, LSM-800) equipped with 5× objective (numerical aperture (NA) = 0.15 dry objective for TCP SP-8), 5× objective (NA = 0.25 dry objective for LSM-800), 10× objective (NA = 0.40 dry objective), 40× objective (NA = 1.30 oil objective), 63× objective (NA = 1.40 oil objective), 100× objective (NA = 1.40 oil objective), and GaAsP detector. The excitation laser was derived from a 405 nm, 488 nm, 561 nm, 640 nm diode laser or a white laser and was set to an appropriate wavelength depending on the dye. Lightning deconvolution process (LAS X 3.5.5, Leica microsystems, Germany) was used in Figure 3f, 3j (for AMPAR), Extended Figure 2b (for mGlu1 and NR1), and Figure 6d. Airyscan mode (Zeiss, Germany) was used in Figure 4e, 4f and Extended Figure 2b (for $GABA_AR$).

**Expression of receptors in HEK293T cells.**

HEK293T cells (ATCC) were cultured in Dulbecco's modified Eagle's medium (DMEM) supplemented with 10% fetal bovine serum (Sigma Aldrich), penicillin (100 units $mL^{-1}$), streptomycin (100 μg $mL^{-1}$) and amphotericin B (250 ng $mL^{-1}$) and incubated in a 5% $CO_2$ humidified chamber at 37 °C. For expression of each receptor, HEK293T cells (2.0 × $10^5$ cells) plated on a 3.5-cm dish (Corning) were transfected with a plasmid encoding rat GluA2 (GluA2flip(Q))[S1], mGlu1[S2], mouse $GABA_AR$ (α1, β3, and γ2L[S3]), rat GluN1-4[S4], or the control vector pCAGGS (kindly provided by Dr. H. Niwa from RIKEN) using Lipofectamine 2000 (Invitrogen) according to the manufacturer's instructions. For live imaging experiments, HEK293T cells were co-transfected with EGFP-F as a transfection marker. The transfected cells were subjected to labeling experiments after 36–48 h of the transfection. For NMDAR expression, 30 μM MK-801 (Hello Bio Inc.) was added to the culture medium to suppress cell death.

**Chemical labeling of receptors in HEK293T cells (Supplementary Figure 3).**

For the labeling of mGlu1, HEK293T cells transfected with mGlu1 were treated with 0.5 μM **CmGlu1M** in the absence or presence of 1.5 μM FITM in the DMEM Glutamax containing 10 mM HEPES at 17ºC for 4 h. For WB analyses of labeled receptors, labeled cells were washed three times with PBS, lysed with radio immunoprecipitation assay (RIPA) buffer containing 1% protease inhibitor cocktail set III (Millipore, 539134), and mixed with 5× Laemmli sample buffer containing 250 mM DTT. SDS PAGE and WB analyses were performed as described in "General methods for biochemical and biological experiments." mGlu1 was detected using a sheep anti-mGlu1 antibody (R&D systems, MAB48361-SP, 1:3,000).

In the case of labeling of NMDAR, HEK293T cells transfected with GluN1-4 were treated with 3 μM **CNR1M** in the absence or presence of 90 μM L-689,560 (Tocris Bioscience) in the DMEM Glutamax containing 10 mM HEPES at 17°C for 4 h. WB were performed as described above. NR1 was detected using a rabbit anti-GluN1 antibody (CST, 224003, 1:2,000).

In the case of labeling of $GABA_AR$, HEK293T cells transfected with $GABA_AR$-α1, $GABA_AR$-β3 and $GABA_AR$-γ2 were treated with 1 μM **CGABAaRM** in the absence or presence of 100 μM Flumazenil (TCI) in the DMEM Glutamax containing 10 mM HEPES at 17°C for 4 h. WB were performed as described above. $GABA_AR$-γ2 was detected using a rabbit anti-$GABA_AR$-γ2 antibody (Synaptic Systems, 224003, 1:2,000).

**Confocal imaging of labeled receptors in HEK293T cells (Supplementary Figure 3).**

After 24 h of transfection, the cells were dissociated by treating with TrypLE Express (Gibco) and re-seeded on 35 mm glass-bottom dishes (Iwaki) pretreated with poly-L-lysine. After 24 h of the re-seeding, the cells were washed twice with 2 mL of DMEM GlutaMax containing HEPES (Gibco). Confocal live imaging of labeled NMDA and mGlu1 receptors were performed using a CLSM (TCS SP-8, Leica microsystems) equipped with a 63×, NA = 1.40 oil objective, and

GaAsP detector. Fluorescence images were acquired using the 488 nm excitation for EGFP and the 633 nm excitation for Alexa Flour 647 derived from a white laser. Confocal live imaging of labeled $GABA_A$ receptors was performed with a confocal microscope (LSM800, Carl Zeiss) equipped with a 63×, NA = 1.4 oil-immersion objective. Fluorescence images were acquired by excitation at 488 nm for EGFP (transfection marker) and 640 nm for Alexa Flour 647 derived from diode lasers.

**Half-life studies of labeled receptors in HEK293T cells by in-gel fluorescence analysis (Supplementary Figure 5).**

For half-life study of labeled receptors, HEK293T cells transfected with the target-receptor plasmids were treated with 2 μM **CAM2-Ax647** for AMPAR, 1 μM **CmGlu1M** for mGlu1, 3 μM **CNR1M** for NMDAR, or 1 μM **CGABAaRM** for $GABA_AR$ in DMEM at 37°C. The cells were incubated for 0, 1, 2, 4, 6, 12, 24, 36, and 48 h. Labeled cells were washed three times with PBS, lysed with RIPA buffer containing 1% protease inhibitor cocktail set III (Millipore, 539134), and mixed with 5× Laemmli sample buffer containing 250 mM DTT. SDS-PAGE and WB analyses were performed as described in "General methods for biochemical and biological experiments." and "Chemical labeling of receptors in HEK293T cells." GluA2 was detected using a rabbit anti-GluA2 antibody (abcam, ab20673, 1:2,000).

The Ax647 labeled receptors were detected and quantified by in-gel fluorescence analysis on SDS-PAGE gels. The target bands were manually selected, and the intensity were calculated with Fusion software (Vilber Lourmat), background intensity was manually subtracted by cutting the minimal intensity in the selected area. The half-life was calculated by curve fitting using KaleidaGraph and following equation (**1**): $I = a + b(k_1/(k_1 - k_2))$ x $(\exp(-k_2\mathrm{t}) – \exp(k_1\mathrm{t}))$, where $k_1$ and $k_2$ were defined as a pseudo first order rate constant for acyl transfer reaction and degradation of labeled receptor, respectively, and the offset value (a) was set equal to zero. The $t_{1/2}$ was defined as $t_{1/2} = \ln(2)/k$.

**Injection of reagents into the mouse Cb.**

Experiments were conducted according to the literature using 4–5 weeks old mice (male, C57BL/6N strain; body weight 18–23 g). [S5] mGlu1 KO mice were purchased from Laboratory Animal Resource Center (Tsukuba University, Japan).[S6] Under the deep anesthesia, **CAM2-Ax647** (50 µM, 4.5 µL, PBS(–)) was directly injected into the vermis of cerebellar lobules V-VIII (0.5 mm depth from the surface) using a microinjector (Nanoliter 2010, world precision instruments) (600 nL/min).

**Chemical labeling of endogenous AMPAR with Cb injection (Figure 2b and d).**

PBS(−) containing 50 µM of **CAM2-Ax647**, DMSO, or **NLC** (4.5 µL) was injected into mouse Cb. In the case of WB analyses, the mouse was sacrificed under the deep anesthesia with isoflurane at 24 h after the injection. The Cb was isolated, washed with PBS twice, and lysed with RIPA buffer containing 1% protease inhibitor cocktail set III (Millipore, 539134). The lysate was incubated at 4°C for 30 min and centrifuged at 4°C and 14,000 g for 10 min. After mixing with 5× Laemmli sample buffer containing 250 mM DTT, SDS-PAGE and WB analyses were performed as described in "General methods for biochemical and biological experiments". The Ax647-labeled AMPAR was detected using a mouse anti-Alexa647 antibody (Immunology Consultants Laboratory, M567-65A-400, 1:1,000). The GluA2 was was detected using a rabbit anti-GluA2 antibody (abcam, ab206293, 1:3,000).

In the case of CLSM analyses, the mouse was transcardially perfused with 4% PFA/PBS(–) at 24 h after the injection. The brain was isolated and sectioned by cryostat (50-µm thick). Imaging was performed using a CLSM equipped with a 5× objective (633 nm excitation for Ax647).

**Perfusion fixation and brain slices preparation.**

Experiments were conducted according to the literature.[S7] Briefly, under the deep anesthesia with isoflurane, mice were perfused transcardially with ice-colded 4% PFA/PBS(–) (pH 7.4). The mouse brain samples were fixed with 4% PFA at 4°C overnight. After washing with PBS(–) (×3), the brain samples were immersed into 30% sucrose/PBS(–). The brain slices were prepared using a cryostat (Leica, CM-1950).

**Whole-cell patch-clamp recording from Purkinje cells in cerebellar acute slices (Figure 2e–p).**

Parasagittal cerebellar slices (200-μm thick) were prepared from C57BL/6N mice (postnatal day 28–35) in which **CAM2-Ax647** was directly injected into the cerebellum, as described previously.[S8] Whole-cell patch-clamp recordings were made from visually identified Purkinje cells using a 60× water-immersion objective attached to an upright microscope (BX51WI, Olympus) at room temperature. The solution used for recording consisted of the following (in mM): 125 NaCl, 2.5 KCl, 2 $CaCl_2$, 1 $MgCl_2$, 1.25 $NaH_2PO_4$, 26 $NaHCO_3$ and 10 D-glucose, bubbled continuously with a mixture of 95% $O_2$ and 5% $CO_2$. Picrotoxin (100 μM, Sigma-Aldrich) was always present in the saline to block the inhibitory inputs. Intracellular solutions were composed of (in mM): 150 Cs-gluconate, 10 HEPES, 4 $MgCl_2$, 4 $Na_2ATP$, 1 $Na_2GTP$, 0.4 EGTA and 5 lidocaine *N*-ethyl bromide (QX-314) (pH 7.25, 292 mOsm/kg). The patch pipette resistance was 2−4 MΩ when filled with each intracellular solution.

To evoke CF- and PF-EPSCs, square pulses were applied through a stimulating electrode placed on the granular layer (10 μs, 0−200 μA) and the molecular layer (~50 μm away from the pial surface; 10 μs, 0−200 μA), respectively. Selective stimulation of CFs and PFs was confirmed by the paired-pulse depression (PPD) and paired-pulse facilitation (PPF) of EPSC amplitudes at a 50-ms interstimulus interval, respectively. The current responses were recorded using an Axopatch 200B amplifier (Molecular Devices), and the pCLAMP system (version 9.2; Molecular Devices) was used for data acquisition and analysis. The signals were filtered at 1 kHz and digitized at 4 kHz. After current recordings, we always confirmed the effective labeling of **CAM2-Ax647** to the slices by using a confocal microscopy (Olympus, FV1000).

**Injection of reagents into the mouse LV.**

Experiments were conducted according to the literature[S9] using 5 weeks old mice (male, C57BL/6N strain; body weight 18–23 g) or 10–15 months old 5xFAD transgenic mice (Tg6799, The Jackson Laboratory). Under the deep anesthesia, the labeling reagent solution (4.5 μL) was directly injected into the LV using a microinjector (Nanoliter 2010, world precision instruments) (600 nL/min).

**Chemical labeling of endogenous receptors with LV injection.**

The labeling reagent was injected into mouse LV. In the case of WB analyses, the mouse was sacrificed under the deep anesthesia with isoflurane at 20–24 h after the injection. The remaining steps were performed as described in "Chemical labeling of endogenous AMPAR with cerebellum injection." The Ax647-labeled receptor was detected using a mouse anti-Alexa647 antibody (Immunology Consultants Laboratory, M567-65A-400, 1:1,000). The NR1 was detected using a mouse anti-NR1 (Novus biologicals lab, NB300-118, 1:1000).

In the case of CLSM analyses, the mouse was transcardially perfused with 4% PFA/PBS(–) at 20–24 h after the injection. The brain was isolated and sectioned by cryostat. Imaging was performed using a CLSM.

**Half-life studies of labeled receptors in the brain by in-gel fluorescence analysis (Figure 5 and Supplementary Figure 5).**

At the indicated time points after the injection of labeling reagent from lateral ventricle, the mouse was sacrificed under the deep anesthesia with isoflurane. The mouse brain tissue was isolated and lysed with RIPA buffer containing 1% protease inhibitor cocktail set III (Millipore, 539134). After mixing with a quarter volume of 5× Laemmli sample buffer containing 250 mM DTT, Electrophoresis and western blotting analyses were performed as described in "General methods for biochemical and biological experiments". The Ax647 labeled receptors were detected and quantified by in-gel fluorescence analysis on SDS PAGE gels. The target bands were manually selected, and the intensity were calculated with Fusion software (Vilber Lourmat), background intensity was manually subtracted by cutting the minimal intensity in the selected area. The half-life was calculated by curve fitting using KaleidaGraph and following equation (**1**): $I = a + b(k_1/(k_1 - k_2))$ x (exp(-$k_2$t) – exp($k_1$t)), where $k_1$ and $k_2$ were defined as a pseudo first order rate constant for acyl transfer reaction and degradation of labeled receptor, respectively, and the offset value (a) was set equal to zero. The $t_{1/2}$ was defined as $t_{1/2} = \ln(2)/k$.

**In-gel tryptic digestion and extraction**

After 20 hours of the injection of **CAM2-Ax647** or DMSO into the mouse lateral ventricle, the mice were sacrificed under the deep anesthesia with isoflurane. After removing the whole brain on ice, the brain tissue were washed with PBS twice and lysed by homogenization and sonication in RIPA buffer. The lysates were incubated at 4°C for 30 min and centrifuged (4°C, 14000 g, 10 min). The protein concentrations in the supernatant were determined by BCA assay and 1 mg of proteins were applied to immunoprecipitation. A small portion of the supernatant was collected as the input fraction. 10 μL of anti-Alexa647 antibody (Immunology Consultants Laboratory, M567-65A-400) was added to the lysate, and it was rotated for 4 h at 4°C. Subsequently, 100 μL of Protein G Sepharose slurry (50% bed vol.) (GE Healthcare) prewashed with RIPA buffer (x 2) was mixed with the lysate. After incubation at 4°C for another 2 h with rotation, the beads were centrifuged (4°C, 1000 g, 3 min), and the supernatant was collected as a flow through fraction. The beads were washed with RIPA buffer (x 10), and boiled at 95°C in 2 x laemmli buffer containing 100 mM DTT for 5 min. The beads were removed by filtration and the filtrate was kept as an immunoprecipitation fraction.

The immunoprecipitated samples were loaded to a 7.5% SDS-PAGE gel (Mini-PROTEAN TGX Gels) and resolved for about 1 cm by the running gel. The gel containing protein samples was manually cut into slices, and fixed with 45% methanol/water containing 5% acetic acid for 20 min. The fixed gel was washed with 50% methanol aq. and pure water. The excised gels were rinsed with pure water, and dehydrated with acetonitrile. The dehydrated gel was swelled with 200 μL of 10 mM DTT in 100 mM TEAB (triethylamine bicarbonate) buffer (Sigma) and incubated at 56°C for 30 min. The DTT solutions were replaced with 55 mM IAA in 100 mM TEAB buffer and incubated at 37°C for 30 min in the dark. The gel pieces were then dehydrated in acetonitrile, rehydrated in 100 mM TEAB buffer, and dehydrated in acetonitrile again. The gel was swelled in 100 mM TEAB buffer containing 10 ng/μL Sequence Grade Trypsin (Promega) and incubated overnight at 37°C. After the digestion, we transferred the supernatant to a new tube and added 50 μL of an extraction solution (50% acetonitrile, 0.1% TFA) to the gel pieces. The supernatants were collected after 10 min incubation, and this process was repeated another two times. Extracted peptides were concentrated by centrifugal concentrator and purified with Stage-tip (GL science).

**NanoLC–MS/MS analyses**

NanoLC–MS/MS analyses were performed on a Q-Exactive mass spectrometer (Thermo Fisher Scientific) and an Ultimate 3000 nanoLC pump (AMR) as described previously.[S10] Samples were automatically injected using PAL system (CTC analytics, Zwingen, Switzerland) into a peptide L-trap column OSD (5 μm) attached to an injector valve for desalinating and concentrating peptides. After washing the trap with MS-grade water containing 0.1% TFA and 2% acetonitrile, the peptides were loaded into a nano HPLC capillary column (C18 packed with the gel particle size of 3 μm, 0.1 × 125 mm, Nikkyo Technos, Tokyo Japan) by switching the valve. The injection volume was 5 μL and the flow rate was 500 nL/min. The mobile phases consisted of (A) 0.5% acetic acid and (B) 0.5% acetic acid and 80% acetonitrile. A two-step linear gradient of 5–45% B in 60 min, 45–95% B in 1 min, 95% B for 20 min was employed. Spray voltages of 2,000 V were applied. The mass scan ranges were m/z 350–1,800, and top ten precursor ions were selected in each MS scan for subsequent MS/MS scans. The normalized collision energy was set to be 30. The raw MS data files were analyzed by Proteome Discoverer 2.2 (Thermo Fisher Scientific) to create peak lists based on the recorded fragmentation spectra. Peptides and proteins were identified by means of automated database searching using Sequest HT (Thermo Fisher Scientific) against UniprotKB/Swiss-Prot release 2021-03 with a precursor mass tolerance of 10 p.p.m., a fragment ion mass tolerance of 0.02 Da, and trypsin specificity that allows for up to three or two missed cleavages. Methionine oxidation was allowed as a variable modification. A reversed decoy database search was conducted to set false discovery rates (FDRs) of less than 1% both at peptide and protein levels. We performed three independent biological replicates in label free quantification analysis. In each experiment, only proteins that identified and quantified by at least two unique peptides were considered hit proteins. In addition, only proteins that were hit more than twice in triplicate were considered “identified proteins” in which keratins, albumin and IgG components (contaminants) were removed.

**Immunohistochemical staining for Homer1, Bassoon, mGlu1, Calbindin, vGluT1, and vGluT2.**

The brain slices (50 μm thickness) were permeabilized with PBS(–) containing 0.1% triton X-100 for 15 min and blocked with 10% normal goat serum (NGS) in PBS(–) containing 0.1% triton X-100 for 30 min. Then, primary antibody reaction was conducted with the following antibodies in PBS(–) containing 0.1% triton X-100 at 4°C overnight. Secondary antibody reaction was conducted with appropriate antibodies in PBS(–) containing 0.1% triton X-100 at r.t. for 1 h.

For Homer1 and Bassoon staining in Figure 3f, a rabbit anti-Homer1 (abcam, ab184955, 1:1000) and a mouse anti-Bassoon (abcam, ab82958, 1:1000) were used as primary antibodies. Secondary antibody reaction was conducted with a goat anti-rabbit IgG H&L (Alexa Fluor® 405) (abcam, ab175652, 1:200) and a goat anti-mouse IgG H&L (Alexa Fluor® 488) (abcam, ab150113, 1:200).

For mGlu1 staining, rabbit anti-mGlu1 in Figure 4c, (Frontier Institute, MSFR104030, 1:1000) as primary antibody. Secondary antibody reaction was conducted with a goat anti-rabbit IgG H&L (Alexa Fluor® 488) (abcam, ab150077, 1:200).

For Calbindin staining in Figure 4e, mouse anti-Calbindin (abcam, ab82812, 1:1000) was used as primary antibody. Secondary antibody reaction was conducted with a donkey anti-mouse IgG H&L (Alexa Fluor® 405) (abcam, ab175658, 1:200).

For Calbindin and Hoechst staining in Extended Data Figure 3e, mouse anti-Calbindin (abcam, ab82812, 1:1000) was used as primary antibody. Secondary antibody reaction was conducted with a goat anti-mouse IgG H&L (Alexa Fluor® 488) (abcam, ab150113, 1:200) and Hoechst 33258 (Dojindo, H341 -Cellstain®- Hoechst 33258 solution, 1:5000).

For Calbindin, vGluT1, and vGluT2 staining Figure 6b–d, a mouse anti-Calbindin (abcam, ab82812, 1:1000), a rabbit anti-vGluT1 (Frontier Institute, VGluT1-Rb-Af500, 1:1000), and a guinea pig anti-vGluT2 (Frontier Institute, VGluT2-GP-Af810, 1:1000) were used as primary antibody. Secondary antibody reaction was conducted with a goat anti-mouse IgG H&L (Alexa Fluor® 405) (abcam, ab175660, 1:200), a goat anti-rabbit IgG H&L (Alexa Fluor® 488) (abcam, ab150077, 1:200), and a goat anti-guinea pig IgG H&L (Alexa Fluor® 555) (abcam, ab150186, 1:200).

**Immunohistochemical staining for GluA2, GABAAR-α1, and NR1.**

For GluA2 staining, the brain slice (15 μm thickness) was attached to glass slide and activated with antigen retrieval reagent ImmunoSaver (FUJIFILM Wako) at 80ºC for 20 min. After blocking with 10% NGS in PBS(–) containing 0.1% triton X-100 for 30 min, primary antibody reaction was conducted with mouse anti-GluA2 (Merck, MAB397, 1:500). Secondary antibody reaction was conducted with a goat anti-mouse IgG H&L (Alexa Fluor® 594) (abcam, ab150116, 1:200) in PBS(–) containing 0.1% triton X-100 at r.t. for 1 h.

For $GABA_A$R-α1 staining, the brain slices (50 μm thickness) were treated with 0.2 % pepsin (from porcine gastric mucosa, Sigma, P-7012) in a buffer (pH 4.0) at 37 °C for 15 min. Permeabilization and blocking was performed in PBS(–) containing 2% BSA, 2% normal goat serum (NGS) and 0.2% triton X-100 at RT for 30 min. After washing with PBS(–) (x 3), the slices was treated with a rabbit anti-$GABA_A$R-α1 antibody (Millipore, 06-868, 1:300) in PBS(–)

containing 0.1% triton X-100 at 4°C overnight. Secondary antibody reaction was conducted with a goat anti-rabbit IgG H&L (Alexa Fluor 546) (Invitrogen, A11071, 1:1,000) in PBS(–) containing 0.1% triton X-100 at RT for 1 h.

For NR1 staining[S11], we used cryosections. Mice were anaesthetized by isoflurane and decapitated. After removing the whole brain, the whole brain vermis was trimmed on ice, mounted on a tissue freezing medium, and frozen using liquid nitrogen. The frozen vermis was cut into 8 μm slices in a cryostat, mounted on Matsunami Adhesive Slice glasses, dried and fixed with 95% (v/v) ethanol at -20 °C for 30 min, followed by acetone on ice for 8 min. The slices were then washed with PBS and treated with 0.1% Triton-X in PBS on ice for 8 min. Slices were incubated with PBS containing 0.3% BSA, followed by 1 μg/mL rabbit polyclonal anti-GluN1 antibody (Frontier Institute, MSFR102650, 1:100) for 4°C O/N, and washed twice with PBS(–) containing 0.3% BSA. The slices were incubated with a goat anti-rabbit IgG H&L (Alexa Fluor 555) (abcam, ab150078, 1:200) for 1 h, washed twice with PBS containing 0.3% BSA, followed by PBS only.

**Tissue clearing of mouse brain with CUBIC protocol.**

Experiments were conducted according to the literature.[S12]

**Figure 3h and i:** In brief, PBS(–) containing 80 μM of **CAM2-Ax647** (4.5 μL) was injected into right and left LVs of the mouse, respectively. 24 h after injection, the mouse was perfused with 4% PFA/PBS(–). The **CAM2-Ax647**-labeled and PFA-fixed brain was delipidated with CUBIC-L, stained with SYTOX-G (for anatomical reference) with CUBIC-HistoVIsion, cleared with CUBIC-R, and embedded in gel (2 % agarose dissolved in CUBIC-R reagent). The 3D imaging was performed with a custom-built LSFM system. The gel-embedded sample was put on a stage in an observation chamber filled with RI-matched silicone oil. Each optical section was illuminated by horizontally-shaped light sheets from the left and right side sequentially, combined with "axial-sweeping" technology.[S13] The emitted fluorescence signal was captured via an oil-dipped objective lens and upright macro-zoom imaging unit (MVX10, Olympus). The thickness of the light-sheet and the z-step size (9 μm) in the scanning motion of the observation chamber were set to be equal to the x-y pixel size (8.2 μm/pixel for MVPLAPO0.63X and 1.25X of zoom), and the stacked volumetric images performed near-isotropic resolution in 3D. 3D image and movie were reconstructed from z-stacked images using IMARIS software (BITPLANE, Oxford Instruments).

**Extended Data Figure 3f and Supplementary Figure 4:** In brief, the **CAM2-Ax647, CNR1M, or CGABAaRM**-labeled and PFA-fixed brain was delipidated with CUBIC-L, cleared with CUBIC-R, and embedded in CUBIC-R solution. The 3D imaging was performed with CLSM (Leica microsystems, Germany, TCS SP-8 or Carl Zeiss, Germany, LSM-800).

**Tissue clearing and FSB staining of 5xFAD mouse brain (Figure 3m).**

PBS(−) containing 80 μM of **CAM2-Ax647** (4.5 μL) was injected into lateral ventricle of 5xFAD mouse. After 24 h, the mouse was transcardially perfused with 4% PFA/PBS(–). The brain was isolated and immersed with 4% PFA/PBS(–) overnight. After conducting delipidation step by using CUBIC-L, the half-brain was stained with FSB[S14] (Dojindo, 30 μg/mL) in PBS(–) containing 1.5 M NaCl for 3 days at room temperature. The sample was washed with PBS(–), and then RI-matched with CUBIC-R.

**The densitiy calculation of AMPAR punctate signals (Figure 3m).**

The densities of Ax647 punctate signals was determined by counting their numbers in several cubic ROIs (40 × 40 × 40 μm, n = 7) of the CUBIC-cleared cortex with IMARIS software. The value of density was calculated by averaging the observed values with considering the expansion ratio after CUBIC protocol (3.3 times). Data are presented as mean ± SEM

**Tissue clearing of mouse brain with 3DISCO protocol (Supplementary Figure 4).**

Experiments were conducted according to the literature.[S15] The fixed brain sample with 4% PFA/PBS(–) was treated in the mixture of THF/$H_2O$ (vol./vol.) at a series of concentrations (50, 70, 80 (each for 12 h) and 100% (12 h × 3) with shaking at RT on a turning table under the dark. Finally, the brain sample was immersed in dibenzyl ether for refractive index matching for 1–2 day before CLSM imaging.

**3D reconstructed from z-stacked images acquired by a confocal microscope (Supplementary Figure 4).**

3D images were reconstructed from z-stacked images using Leica LAS-X or Zeiss ZEN. Compression in the z-direction caused by spherical aberration was corrected according to the following formula.[S16]

$$d'/d = \{\tan (\sin^{-1} (0.5\ NA/ n_1))\}/ \{\tan (\sin^{-1} (0.5\ NA/ n_2))\}$$

$d'$ is the actual focal position.

$d$ is the expected focal position.

$n_1$ is the refractive index of air (1.00).

$n_2$ is the refractive index of DBE (1.54) or CUBIC-R (1.52).

**The densitiy calculation of AMPAR and GABA$_A$R punctate signals (Figure 4f).**

The densities of Ax555 and Ax647 punctate signals were determined by counting their numbers in several cubic ROIs (10 × 10 × 10 μm, n = 7) in the molecular layer of mouse Cb cleared by CUBIC protocol. The value of density was calculated by averaging the observed values with considering the expansion ratio of Cb after CUBIC protocol (3.3 times). Data are presented as mean ± SEM.

**Intracerebroventricular injection of reagents for the neonatal mouse (Figure 6, Extended Data Figure 3–6).**

Experiments were conducted according to the literature.[S17] Briefly, pups (C57BL/6N strain) were anesthetized by 2–4% isoflurane and clamped in a stereotactic apparatus. After the scalp was opened slightly, a small hole was made in the skull with a 27G needle. The labeling reagent solution (PBS(–) containing 80 μM **CAM2-Ax647**, 2.0 μL) was directly injected using a microinjector (Nanoliter 2010, world precision instruments, 600 nL/min) with a finely pulled glass pipette.

#### Synthesis and Characterization of Compounds

##### General materials and methods for organic synthesis

All chemical reagents and solvents were obtained from commercial suppliers (SIGMA-Aldrich, Tokyo Chemical Industry (TCI), FUJIFILM Wako Pure Chemical Corporation, or Watanabe Chemical Industries) and used without further purification. Thin layer chromatography (TLC) was performed on silica gel 60 $F_{254}$ precoated aluminum sheets and glass plate (Merck) and visualized by fluorescence quenching or ninhydrin staining. Chromatographic purification was conducted by flash column chromatography on silica gel 60N (neutral, 40–50 µm, Kanto Chemical) or Biotage isolera system equipped with a SNAP ultra cartridge. $^{1}$H NMR spectra were recorded in deuterated solvents on a JEOL ECS400 (400 MHz) or JEOL ECZ600R (600 MHz). Chemical shifts were referenced to residual solvent peaks or tetramethylsilane (δ = 0 ppm). Multiplicities are abbreviated as follows: *s* = singlet, *d* = doublet, *t* = triplet, *q* = quartet, *quin* = quintet, *m* = multiplet. MALDI-TOF Mass spectra were measured on UltrafleXtreme (Bruker Daltonics). High resolution mass spectra were measured on an Exactive (Thermo Scientific) equipped with electron spray ionization (ESI). Reversed-phase HPLC (RP-HPLC) was carried out on a Hitachi Chromaster system equipped with a diode array using a Cosmosil 5C18AR2 (Nacalai tesque) and a YMC-Pack ODS-A column (YMC Co. Ltd.).

**Synthesis of CAM2-Ax555**

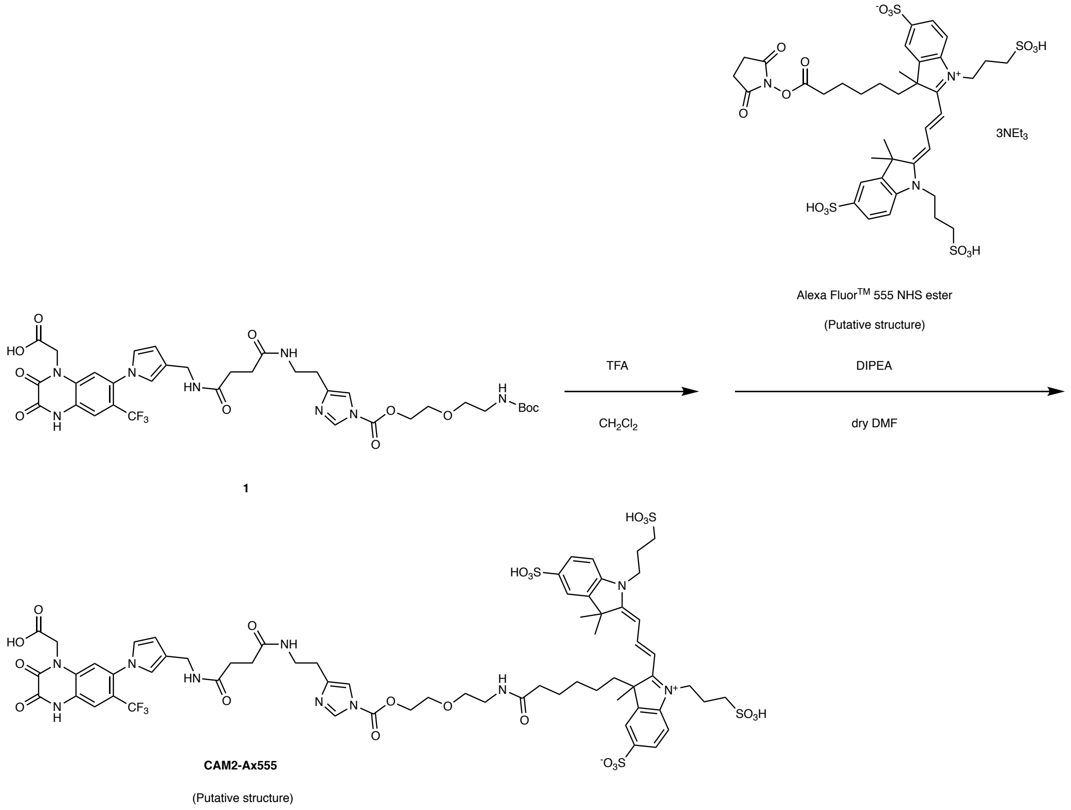


**CAM2-Ax555:** The compound **1** was synthesized according to the previously reported method.[S1] To a solution of **1** (0.6 mg, 0.74 μmol) in dry $CH_2Cl_2$ (1.0 mL), TFA (0.25 mL) was added and the reaction mixture was stirred at room temperature for 1 h. After azeotropically removal of the solvent with $CH_2Cl_2$ (×1) and toluene (×2), the Boc-deprotected **1** was dissolved in dry DMF (0.4 mL). Alexa Fluor™ 555 NHS ester (1.0 mg, 0.81 μmol) and DIPEA (1.5 μL, 8.6 μmol) were added and the mixture was stirred at room temperature for 6 h. The reaction mixture was purified by RP-HPLC (column; Cosmosil 5C18AR2, 250 x 20 mm, mobile phase; $CH_3CN$ : 10 mM $AcONH_4$ aq. = 0 : 100 → 25 : 75 (linear gradient over 70 min), flow rate; 9 mL/min, detection; UV absorbance at 220 nm) to give **CAM2-Ax555** as a red solid (0.27 μmol, 37% yield, determined by measurement of UV-absorbance). HR ESI-MS (calc. for $C_{64}H_{75}F_3N_{10}O_{22}S_4$): $[M-2H]^{2-}$ = 781.1702 (calc. = 781.1693).

#### Synthesis of CAM2-Cy5 and CAM2-SulfoCy5

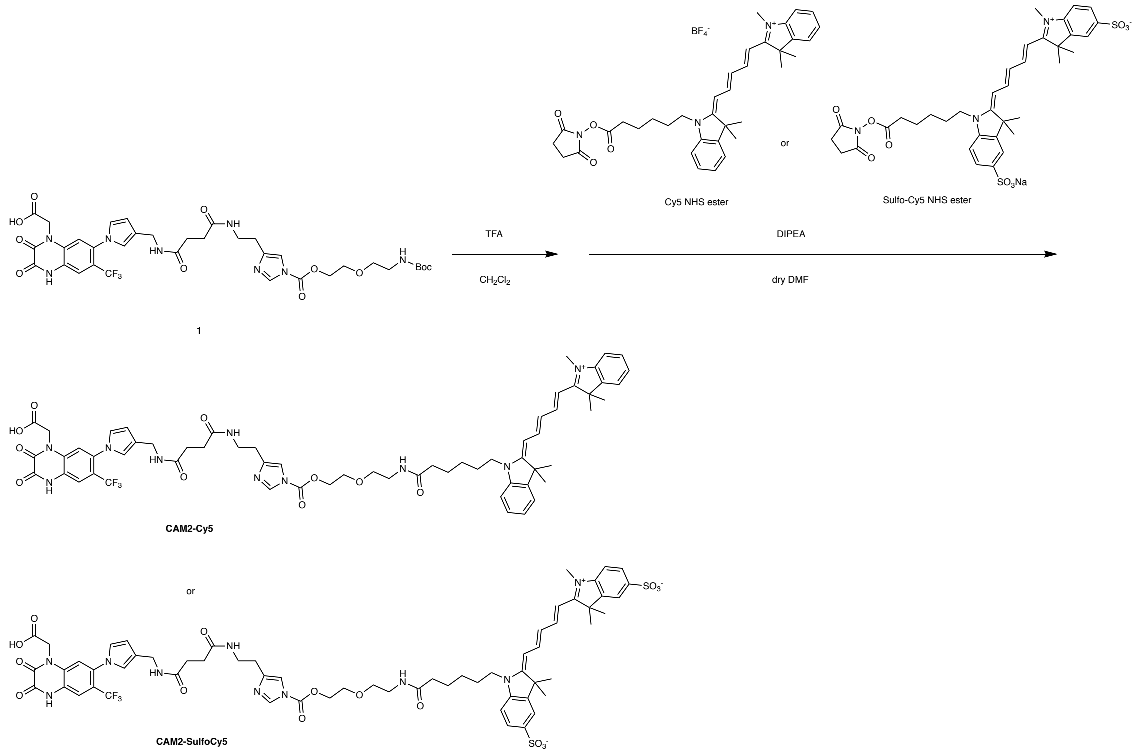


**CAM2-Cy5 and CAM2-SulfoCy5:** These compounds were prepared from compound **1** according to the similar way for the synthesis of **CAM2-Ax555**. HR ESI-MS **CAM2-Cy5** (calc. for $C_{62}H_{70}F_3N_{10}O_{10}$): $[M+H]^+$ = 1171.5212 (calc. = 1171.5223). **CAM2-SulfoCy5** (calc. for $C_{62}H_{67}F_3N_{10}O_{16}S_2$): $[M-2H]^{2-}$ = 664.2080 (calc. =664.2070).

### Synthesis of CmGlu1M

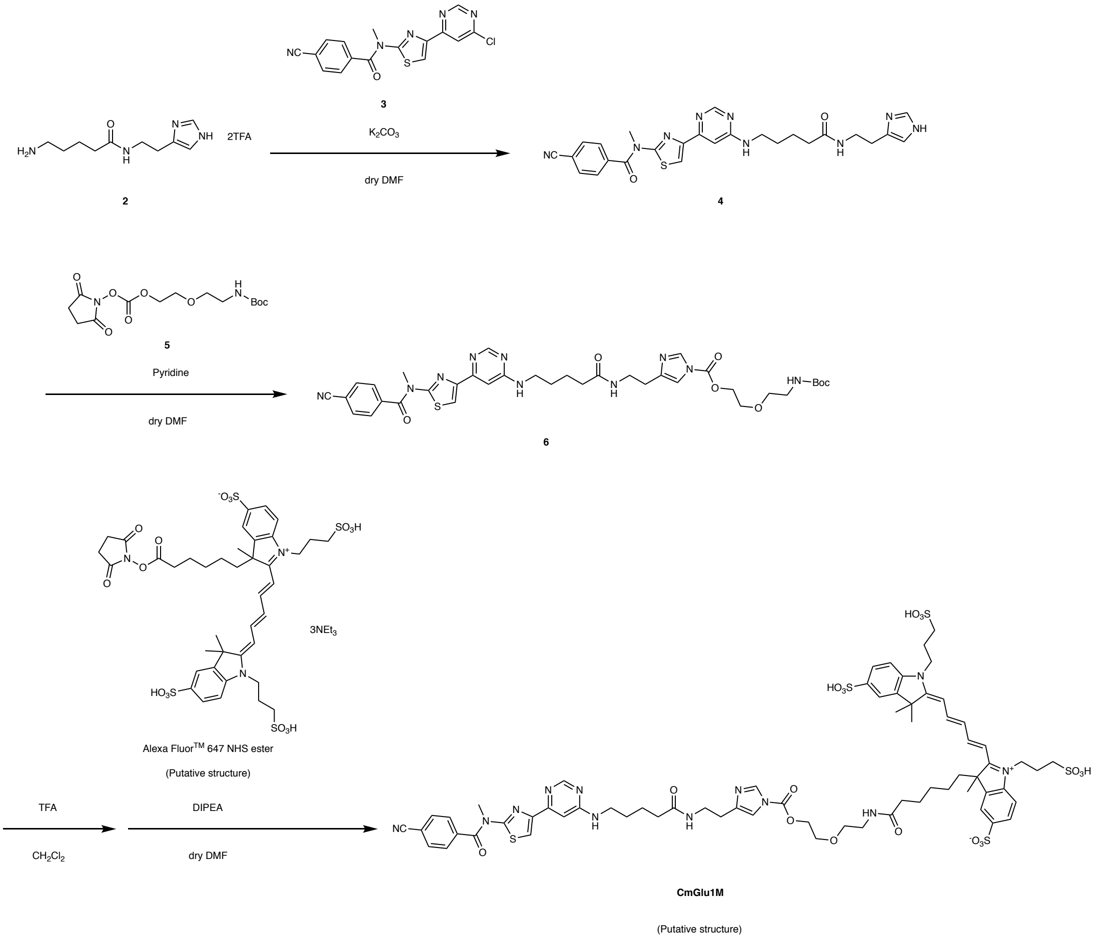

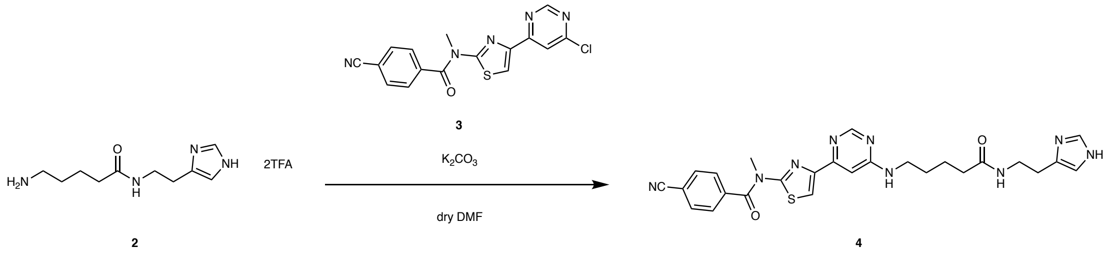


***N*-(4-(6-((5-((2-(1*H*-imidazol-4-yl)ethyl)amino)-5-oxopentyl)amino)pyrimidin-4-yl)thiazol-2-yl)-4-cyano-*N*-methylbenzamide (4):** The compound **2** and **3** were synthesized according to the literature.[S18,S19] To the solution of compound **2** (34 mg, 68 μmol, 1.0 eq.) in dry DMF (0.8 mL) was added compound **3** (24 mg, 68 μmol, 1.0 eq.) and $K_2CO_3$ (55 mg, 400 μmol, 6.0 eq.). After stirring at 75°C for 6 h under argon atmosphere, the reaction mixture was diluted with 2 ml of water and extracted with 20 ml of $CH_2Cl_2$ (×5). The organic layer was dried over $MgSO_4$ and evaporated under reduced pressure. The residue was purified by silica gel chromatography ($CH_2Cl_2$:$CH_3OH$:$NH_3$ aq. = 8:1:0.01). After evaporation, the resulting residue was further purified by reverse-phase HPLC (mobile phase; 0.1% TFA/$CH_3CN$ : 0.1% TFA/$H_2O$ = 20 : 80 → 45 : 55 (linear gradient over 25 min), flow rate = 3 ml/min, detection; UV absorbance at 220 nm) to give a colorless oil (5.0 mg, 14%).$^1$H NMR (DMSO-$d_6$, 600 MHz, 373 K) δ 8.83 (1H, *s*), 8.47 (1H, *s*), 8.04 (1H, *s*), 7.97 (2H, *d*, $J$ = 0.9 Hz), 7.84 (2H, *d*, $J$ =0.9 Hz), 7.33 (1H, *s*), 7.14 (1H, *s*), 3.62 (3H, *s*), 3.36 (4H, *m*), 2.80 (2H, *t*, $J$ = 6.6 Hz), 2.12 (2H, *t*, $J$ = 6.6 Hz), 1.57 (4H, *m*). $^{13}$C NMR (DMSO, 150 MHz, 373 K) δ 172.76, 169.16, 163.81, 160.58, 157.67, 154.405, 146.87, 139.33, 134.19, 133.01, 131.98, 128.82, 118.45, 118.31, 116.99, 116.63, 116.34, 114.05, 100.90, 38.20, 38.03, 35.67, 28.95, 25.26, 23.21. HR ESI-MS (calc. for $C_{26}H_{27}N_9O_2S$): $[M+Na]^+$ = 552.1903 (calc. = 552.1901).

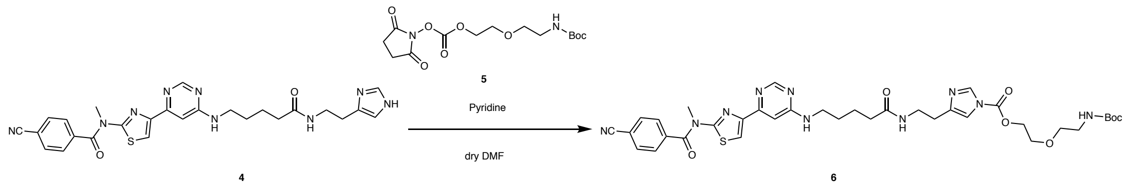


**2-(2-((*tert*-butoxycarbonyl)amino)ethoxy)ethyl 4-(2-(5-((6-(2-(4-cyano-*N*-methylbenzamido) thiazol-4-yl)pyrimidin-4-yl)amino)pentanamido)ethyl)-1*H*-imidazole-1-carboxylate (6):** The compound **5** was synthesized by a previously reported method.[S20] To the solution of compound **4** (34 mg, 64 µmol, 1.0 eq.) and **5** (33 mg, 95 µmol,1.5 eq.) in dry DMF (2 mL) was added dry pyridine (40 µl, 500 µmol, 7.7 eq.). After stirring for 3.5 h at room temperature under argon atmosphere, the solvent was removed by evaporation. The residue was purified by silica gel chromatography with a linear gradient of 1–15% $CH_3OH/CH_2Cl_2$). After evaporation, the residue was purified by reverse-phase HPLC (column; Cosmosil 5C18AR2, 250 x 20 mm, mobile phase; 0.1% TFA/$CH_3CN$ : 0.1% TFA/$H_2O$ = 20 : 80 → 45 : 55 (linear gradient over 25 min), flow rate = 9 ml/min, detection; UV absorbance at 220 nm) to give a colorless solid (42 mg, 84%). $^1$H NMR ($CDCl_3$, 400 MHz) δ 8.52 (1H, *s*), 8.09 (1H, *s*), 7.95 (1H, *s*), 7.82 (2H, *d*, *J* = 0.8 Hz), 7.68 (2H, *d*, *J* =0.8 Hz), 7.20 (1H, *s*), 7.05(1H, *s*), 4.51 (2H, *m*), 3.76 (2H, *m*), 3.71 (3H, *s*), 2.12 (2H, *t*, *J* = 6.6 Hz), 1.57 (2H, *m*) 3.52 (4H, *m*), 3.39 (2H, *br*), 3.31 (4H, *m*), 2.73 (2H, *t*, *J* = 6.0 Hz), 2.24 (2H, *t*, *J* = 7.2 Hz), 1.76 (2H, *m*), 1.67 (2H, *m*).$^{13}$C NMR ($CDCl_3$, 150 MHz) δ 172.58, 168.34, 163.13, 159.68, 158.31, 155.90, 148.01, 141.84, 138.52, 136.93, 132.57, 128.18, 117.72, 116.26, 114.73, 113.66, 79.42, 70.28, 68.28, 66.93, 53.41, 40.22, 38.78, 38.17, 35.96, 28.59, 28.35, 27.35, 22.64, 22,16. HR ESI-MS (calc. for $C_{36}H_{44}N_{10}O_7S$): $[M+Na]^+$ = 783.3005 (calc. = 783.3007).

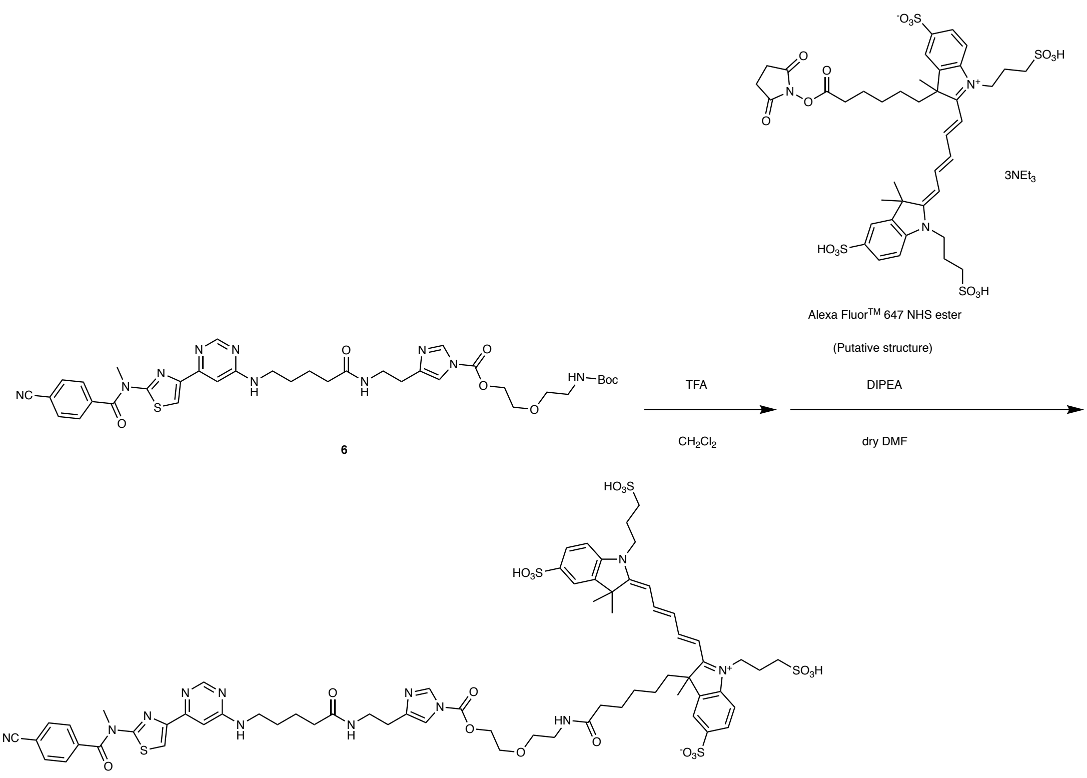


**CmGlu1M**

(Putative structure)

**CmGlu1M:** To a solution of compound **6** (1.0 mg, 1.97 μmol) in dry $CH_2Cl_2$ (2.0 mL), TFA (0.5 mL) was added and the reaction mixture was stirred at room temperature for 1 h under argon atmosphere. After removal of the solvent by evaporation, the residual TFA was azeotropically removed with toluene (×3). The crude product was used for the next step without further purification. To a solution of the Boc-deprotected **6** in dry DMF (0.5 mL), Alexa Fluor™ 647 NHS ester (1.0 mg, 0.80 μmol) and DIPEA (2.4 μL, 14 μmol) were added and stirred at room temperature for 18 h. The reaction mixture was purified by RP-HPLC (Cosmosil 5C18AR2, 250 x 10 mm, mobile phase; $CH_3CN$ : 10 mM $NH_4OAc$ aq. = 15: 85 → 60 : 70 (linear gradient over 45 min), flow rate; 9 mL/min, detection; UV absorbance at 220 nm) to give **CmGlu1M** as a blue solid (0.59 μmol, 75% yield, determined by measurement of UV-absorbance). HR ESI-MS (calc. for $C_{67}H_{80}N_{12}O_{18}S_5$): $[M-3H]^{3-}$ = 499.1365 (calc. = 499.1366).

#### Synthesis of Compound 8

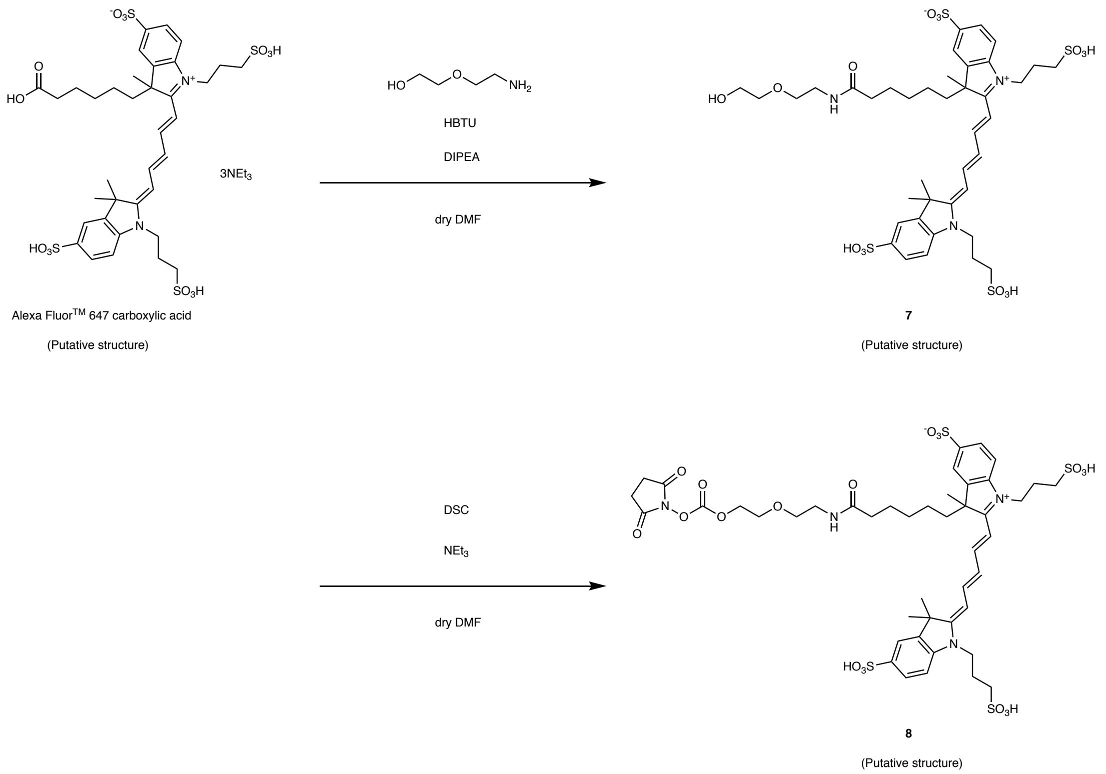


**Compound 7:** To a solution of Alexa Fluor™ 647 carboxylic acid (15.0 mg, 17.4 μmol) in dry DMF (2.0 mL), HBTU (20.0 mg, 52.7 μmol), DIPEA (6.0 μL, 34 μmol), and 2-(2-aminoethoxy)ethanol (7.0 μL, 70 μmol) were added. After stirring the reaction mixture at room temperature under Ar atmosphere for 48 h, the mixture was purified by reverse-phase HPLC (column; Cosmosil 5C18AR2, 250 x 20 mm, mobile phase; 0.1% TFA/$CH_3CN$ : 0.1% TFA/$H_2O$ = 0 min; 0:100 → 5 min; 0:100 → 40 min; 30:70 (linear gradient over 35 min), flow rate = 9 ml/min, detection; UV absorbance at 220 nm) to give a compound **7** as a blue solid (13.4 mg, 81% yield). HR ESI-MS (calc. for $C_{40}H_{55}N_3O_{15}S_4$): $[M+Na-3H]^{2-}$ = 482.6092 (calc. = 482.6095).

**Compound 8:** To a solution of compound **7** (11.8 mg, 12.4 μmol) dissolved in dry DMF (0.8 mL), *N*,*N*'-disuccinimidyl carbonate (DSC) (32 mg, 124 μmol) and triethylamine (16.9 μL, 124 μmol) were added. The reaction mixture was stirred at room temperature under Ar atmosphere for 12 h and purified by RP-HPLC (Cosmosil 5C18AR2, 250 x 10 mm, mobile phase; 0.1% TFA/$CH_3CN$ : 0.1% TFA/$H_2O$ = 0 : 100 → 30 : 70 (linear gradient over 35 min), flow rate; 9

mL/min, detection; UV absorbance at 650 nm) to give a compound **8** as a blue solid (6.4 mg, 47% yield). HR ESI-MS (calc. for $C_{45}H_{58}N_4O_{19}S_4$): $[M-2H]^{2-}$ = 542.1225 (calc. = 542.1216).

#### Synthesis of CNR1M

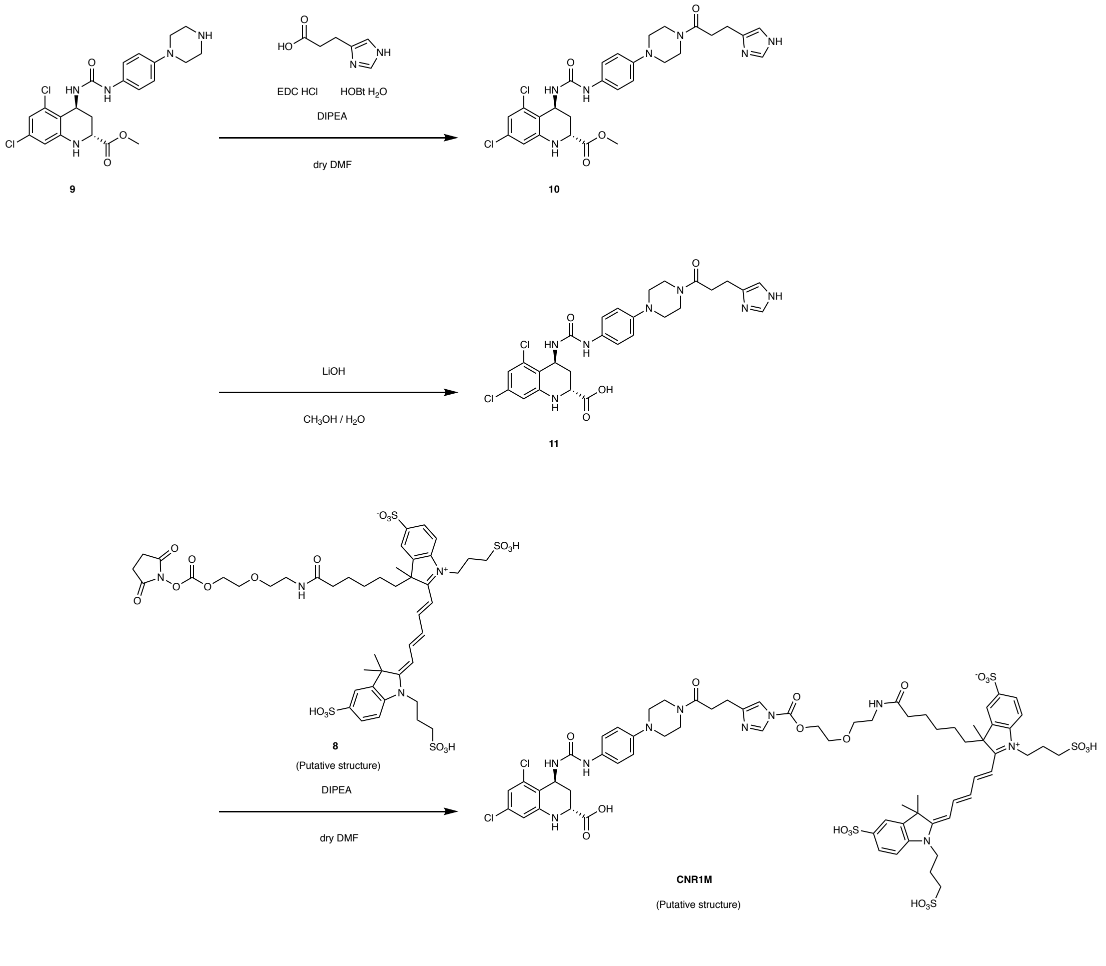


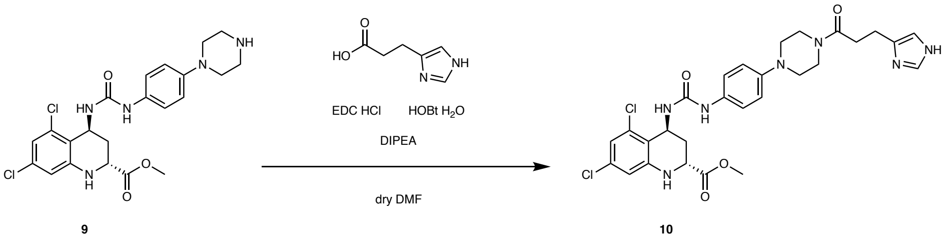


**Methyl (2*R*,4*S*)-4-(3-(4-(4-(3-(1*H*-imidazol-4-yl)propanoyl)piperazin-1-yl)phenyl)ureido)-5,7-dichloro-1,2,3,4-tetrahydroquinoline-2-carboxylate (10):** The compound **9** was synthesized according to the literature.[S21] To a solution of compound **9** (20.2 mg, 42.2 μmol) in dry DMF (1 mL), 3-(1*H*-imidazol-4-yl)propanoic acid (8.0 mg, 50.7 μmol), EDC HCl (10.9 mg, 50.7 μmol), HOBt $H_2O$ (8.7 mg, 50.7 μmol) and DIPEA (23 μL, 126 μmol) were added. The reaction mixture was stirred at room temperature under Ar atmosphere for 40 h. After removal of the solvent by evaporation, the residue was purified by silica gel chromatography ($CHCl_3$:$CH_3OH$:$NH_3$ aq. = 5:1:0.01) to give a compound **10** (17.3 mg, 28.8 μmol, 68% yield) as a white solid. $^1$H NMR (600 MHz, $CD_3OD$): δ 7.84 (1H, *d*, *J* = 1.2 Hz), 7.26 (2H, *dd*, *J* = 7.2 Hz, 2.4 Hz), 6.94 (1H, *d*, *J* = 1.2 Hz), 6.92 (2H, *dd*, *J* = 7.2 Hz, 2.4 Hz), 6.72 (1H, *d*, *J* = 2.4 Hz), 6.68 (1H, *d*, *J* = 2.4 Hz), 5.08 (1H, *t*, *J* = 3.0 Hz), 4.03 (1H, *dd*, *J* = 12.6 Hz, 2.4 Hz), 3.79 (3H, *s*), 3.72 (2H, *t*, *J* = 3.0 Hz), 3.66-3.64 (2H, *m*), 3.02 (4H, *m*), 2.92 (2H, *t*, *J* = 7.2 Hz), 2,77 (2H, *t*, *J* = 7.2 Hz), 2.50 (1H, *dt*, *J* = 13.2 Hz, *J* = 8.4 Hz), 1.70 (1H, *ddd*, *J* = 12.6 Hz, *J* = 2.4 Hz). $^{13}$C NMR (150 MHz, $CD_3OD$): δ 174.15, 172.81, 157.15, 148.32, 147.76, 137.02 (2H), 135.69, 135.62, 134.00, 121.92 (2H), 118.80 (2H), 117,88. 117.40, 116.11, 113.99, 52.94, 51.72, 51.33, 50.48, 46.79, 44.49, 42.93, 33.50, 33.08, 23.15. HR ESI-MS (calc. for $C_{28}H_{31}$ $Cl_2N_7O_4$): $[M+H]^+$ =622.1710 (calc. = 622.1707).

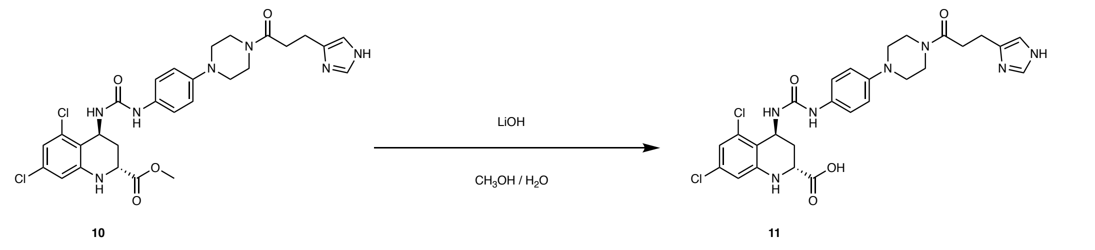


**(2*R*,4*S*)-4-(3-(4-(4-(3-(1*H*-imidazol-4-yl)propanoyl)piperazin-1-yl)phenyl)ureido)-5,7-dichloro-1,2,3,4-tetrahydroquinoline-2-carboxylic acid (11):** To a solution of compound **10** (4.1 mg, 6.8 μmol) in $CH_3OH$ (0.5 mL), 1 M LiOH aq. (0.1 mL, 0.1 mmol) was added and the reaction mixture was stirred at room temperature for 3 h. After neutralization with 1 N HCl aq., the solvent was removed by evaporation to give a compound **11** as a white solid. The compound **11** was used for the next step without further purification.

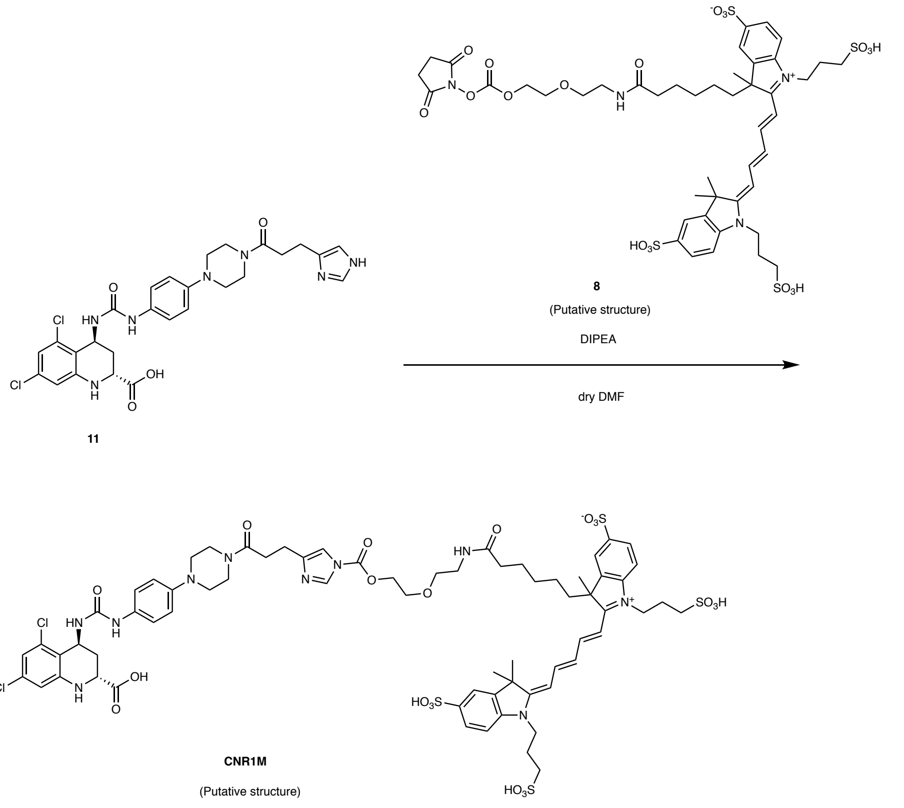


**CNR1M:** To a solution of compound **11** (3.2 mg, 5.4 μmol) in dry DMF (0.7 mL), the compound **8** (6.4 mg, 4.9 μmol) and DIPEA (10 μL, 57.4 μmol) were added and the mixture was allowed to stir at room temperature under $N_2$ atmosphere for 13 h. The reaction solution was purified by RP-HPLC (Cosmosil 5C18AR2, 250 x 10 mm, mobile phase; $CH_3CN$ : 10 mM $NH_4OAc$ aq. = 5 : 95 → 45 : 55 (linear gradient over 40 min), flow rate; 10 mL/min, detection; UV absorbance at 220 nm) to give **CNR1M** as a blue solid (2.2 mg, 27% yield). HR ESI-MS (calc. for $C_{68}H_{82}Cl_{2}N_{10}O_{20}S_4$): $[M-2H]^{2-}$ = 777.1917 (calc. = 777.1911).

### Synthesis of CGABAaRM

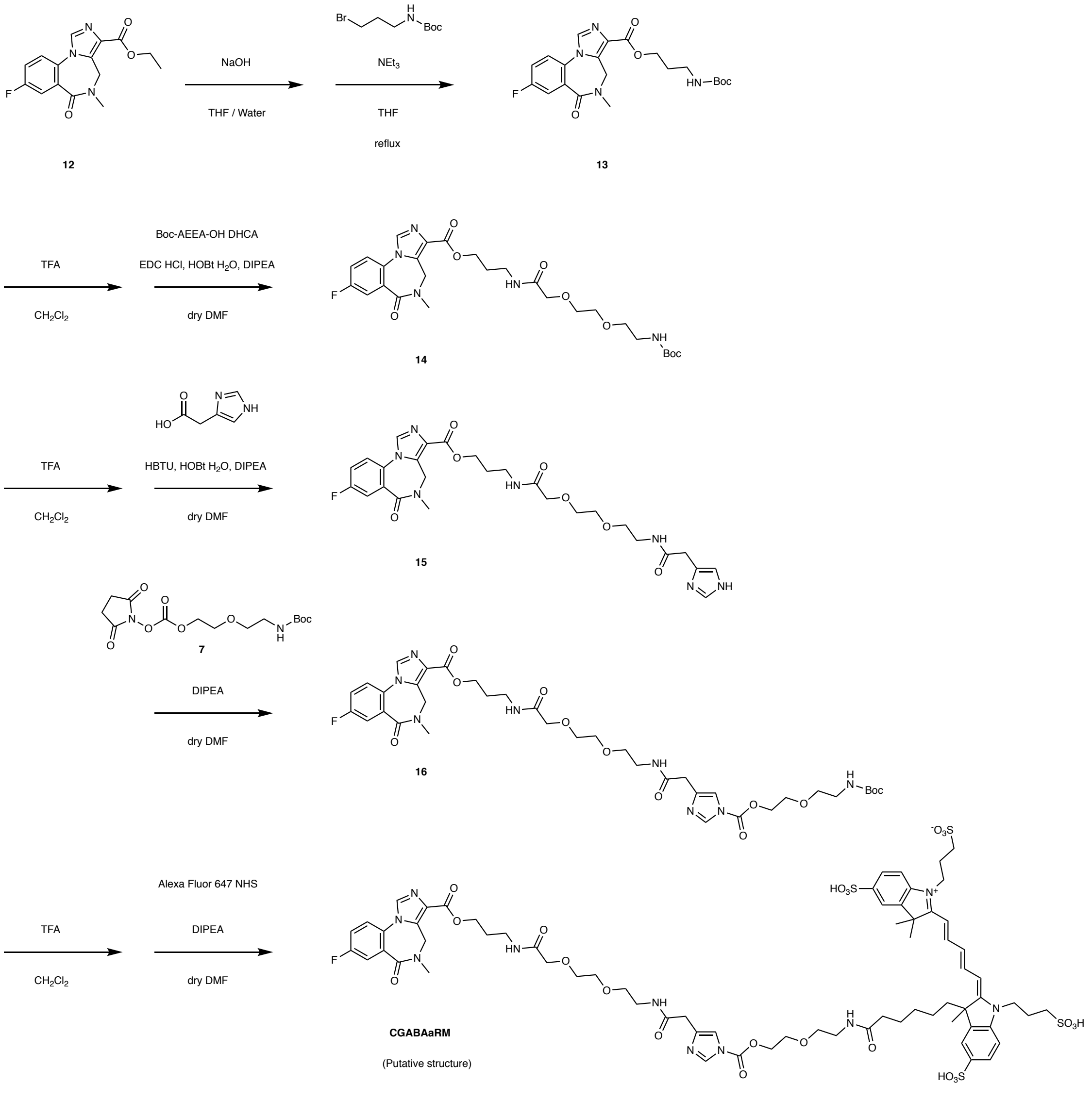

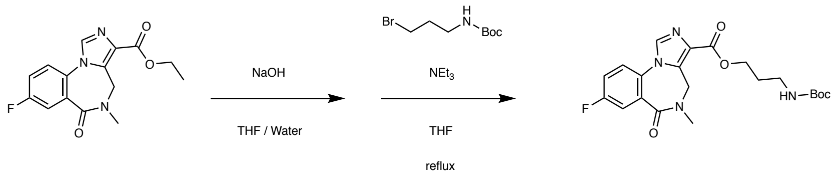


12 13

**3-((*tert*-butoxycarbonyl)amino)propyl 8-fluoro-5-methyl-6-oxo-5,6-dihydro-4*H*-benzo[*f*]imidazo[1,5-*a*][1,4]diazepine-3-carboxylate (13):** Flumazenil (**12**) (188 mg, 0.619 mmol) was suspended in the mixture of THF (3 mL) and $H_2O$ (2 mL). Sodium hydroxide (0.464 mL of a 2 M solution dissolved in water, 0.929 mmol, 1.5 eq.) was added the suspension was stirred at 40 °C for 1.5 h until judged complete by TLC. The solution was neutralized by the addition of 2M HCl (0.464 mL, 0.929 mmol). The solvent was evaporated and the residue was dried under reduced pressure to give a white solid. The obtained product was suspended in dry THF (10 mL). 3-(*tert*-Butoxycarbonylamino)propyl bromide (445 mg, 1.86 mmol, 3 eq.) and triethylamine (0.86 mL, 6.19 mmol, 10 eq.) were added and the mixture was refluxed under Ar atmosphere for 13 h. The reaction mixture was diluted with $CH_2Cl_2$ (50 mL) and washed with brine (50 mL x 1). The aqueous layer was extracted with $CH_2Cl_2$ (50 mL x 1). The combined organic layer was dried over $MgSO_4$ and evaporated under reduced pressure. The residue was purified by silicagel chromatography to give a compound **13** as a clear oil ($CHCl_3$ : $CH_3OH$ = 50 : 1) (313 mg, quant.). $^1$H NMR (600 MHz, $CDCl_3$): δ 7.86 (1H, s), 7.77(1H, *dd*, *J* = 8.4 Hz, *J* = 2.4 Hz), 7.42 (1H, *dd*, *J* = 9.0 Hz, *J* = 4,2 Hz), 7.35 (1H, *m*), 5.22 (1H, *d*, J = 12.0), 4.95 (1H, s), 4.43 (2H, *s*, *J* = 6.4 Hz), 4.36 (1H, d, *J* = 12.0 Hz), 3.28 (2H, *q*, *J* = 6.0 Hz), 3.42 (3H, s), 1.99 (2H, *t*, *J* = 6.4 Hz), 1.43 (9H, s). $^{13}$C NMR (150 MHz, $CDCl_3$): δ 165.17, 162.97, 161.76 (*d*, *J* = 250.5 Hz), 155.94, 135.55, 134.90, 131.21 (*d*, *J* = 7.63 Hz), 128.52, 128.24 (*d*, *J* = 2.54 Hz), 123.84 (*d*, *J* = 7.63 Hz), 120.01 (*d*, *J* = 22.89 Hz), 119.39 (*d*, *J* = 25.43 Hz). 79.20, 62.60, 42.27, 37.47, 35.95, 29.27, 28.36. HR ESI-MS (calc. for $C_{21}H_{25}FN_4O_5$): $[M+H]^+$ = 433.1879 (calc. = 433.1882), $[M+Na]^+$ = 455.1697 (calc. = 455.1701), $[M-H]^-$ = 431.1730 (calc. = 431.1736).

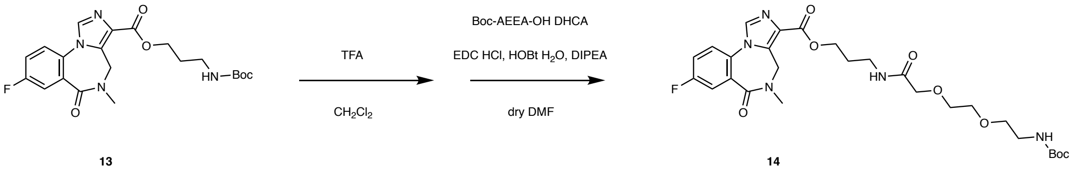


**2,2-dimethyl-4,13-dioxo-3,8,11-trioxa-5,14-diazaheptadecan-17-yl 8-fluoro-5-methyl-6-oxo-5,6-dihydro-4*H*-benzo[*f*]imidazo[1,5-*a*][1,4]diazepine-3-carboxylate (14):** To a stirred solution of compound **13** (82 mg, 0.197 mmol) in $CH_2Cl_2$ (2 mL) was added trifluoroacetic acid (TFA) (1 mL) at room temperature. The solution was allowed to stir for 1 h. TFA was removed by co-evaporation with toluene under reduced pressure (5 mL x 3). The residue was dissolved in dry DMF (5 mL) and Boc-AEEA-OH DCHA (107 mg, 0.24 mmol, 1.2 eq.), EDC HCl (46 mg, 0.24 mmol, 1.2 eq.), HOBt $H_2O$ (37 mg, 0.24 mmol, 1.2 eq.) and DIEPA (350 µL, 2.4 mmol, 20 eq.) were added. The reaction mixture was stirred at room temperature under Ar atmosphere for 16 h. After removal of DMF by evaporation, the residue was dissolved in $CH_2Cl_2$ (30 mL) and washed with 5% citric acid aq. (10 mL x 1), sat. $NaHCO_3$ aq. (10 mL x1), and brine (10 mL x 1). The organic layer was dried over $MgSO_4$ and evaporated under reduced pressure. The residue was purified by silicagel chromatography with a linear gradient of $CH_3OH$–$CHCl_3$, 1%–10%) to give a compound **14** as a clear oil (109 mg, 0.189 mmol, 96% yield). $^1$H NMR (600 MHz, $CDCl_3$): δ 7.87 (1H, *s*), 7.77 (1H, *dd*, *J* = 9.0 Hz, *J* = 3.0 Hz), 7.46 (1H, *dd*, *J* = 9.0 Hz, *J* = 4.2 Hz), 7.35 (1H, *m*), 7.13 (1H, *s*), 5.22 (1H, *bd*, *J* = 12.6 Hz), 5.09 (1H, *s*), 4.43 (1H, *s*), 4.37 (1H, *bd*, *J* = 12.6 Hz), 3.99 (2H, *s*), 3.68 (2H, *m*), 3.64 (2H, *m*), 3.55 (2H, *t*, *J* = 5.4 Hz), 3.47 (2H, *dd*, *J* = 11.4 Hz, *J* = 7.2 Hz), 3.33 (2H, *dd*, *J* = 15.0 Hz, *J* = 4.8 Hz), 3.24 (3H, s), 2.05 (2H, *quin*, *J* = 6.6 Hz), 1.41 (9H, s). $^{13}$C NMR (150 MHz, $CDCl_3$) δ 170.00, 165.16, 162.96, 161.76 (*d*, *J* = 251.77), 155.97, 135.62, 134.96, 131.22 (*d*, *J* = 7.63), 128.46, 128.23 (*d*, *J* = 2.54), 123.84 (*d*, *J* = 7.63), 120.01 (*d*, *J* = 22.89, 119.37 (*d*, *J* = 25.43), 79.31, 70.96, 70.49, 70.34, 69.90, 62.31, 42.29, 40.18, 35.92, 35.5, 28.94, 28.36. HR ESI-MS (calc. for $C_{27}H_{36}FN_5O_8$): $[M+Na]^+$ = 600.2440 (calc. = 600.2440), $[M-H]^-$ = 576.2472 (calc. = 576.2475).

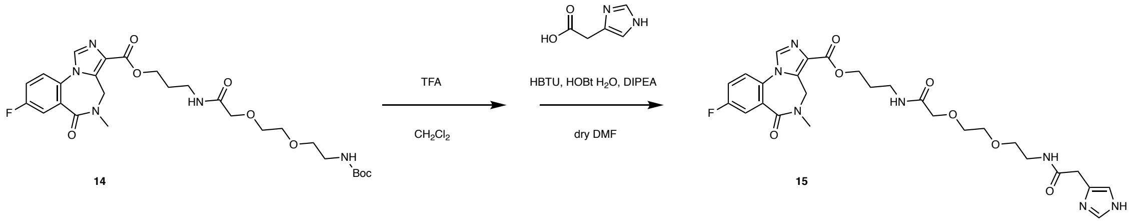

**1-(1*H*-imidazol-4-yl)-2,11-dioxo-6,9-dioxa-3,12-diazapentadecan-15-yl 8-fluoro-5-methyl-6-oxo-5,6-dihydro-4*H*-benzo[*f*]imidazo[1,5-*a*][1,4]diazepine-3-carboxylate (15):** To a stirred solution of compound **14** (44 mg, 0.076 mmol) in dry $CH_2Cl_2$ (2 mL) was added trifluoroacetic acid (TFA) (1 mL) at room temperature. The solution was allowed to stir for 1 h. TFA was removed by co-evaporation with toluene under reduced pressure (5 mL x 3). The residue was dissolved in dry DMF (2 mL) and 4-imidazolecarboxylic acid HCl (35 mg, 0.091 mmol, 1.2 eq.), HBTU (35 mg, 0.091 mmol, 1.2 eq.), and DIEPA (132 µL, 0.76 mmol, 10 eq.) were added. The reaction mixture was stirred at room temperature under Ar atmosphere for 16 h. After removal of DMF by evaporation, the residue was purified by silicagel chromatography with a linear gradient of 0–34% $CH_3OH/CHCl_3$-1% $NH_3$ (12 CV) to give a compound **15** as a clear oil (45 mg, 0.068 mmol, 90%). $^1H$ NMR (600 MHz, $CDCl_3$): δ 7.93 (1H, *s*), 7.79 (1H, *dd*, *J* = 9.0 Hz, *J* = 3.0 Hz), 7.47 (1H, *dd*, *J* = 8.4 Hz, *J* = 4,2 Hz), 7.39–7.36 (1H, *m*), 7.24 (1H, *s*), 6.92 (1H, *s*), 5.23 (1H, *bd*, *J* = 14.4 Hz), 4.40 (3H, *bs*), 3.99 (2H, *s*), 3.70–3.67 (2H, *m*), 3.64–3.61 (2H, *m*), 3.58 (1H, *s*), 3.56 (2H, *t*, *J* = 5.4 Hz), 3.46 (2H, *dd*, *J* = 10.8 Hz, *J* = 5.4 Hz), 3.37 (2H, *dd*, *J* = 8.8 Hz, *J* = 7.2 Hz), 3.26 (3H, *s*), 1.92 (2H, *quin*, *J* = 6.6 Hz). $^{13}C$ NMR (150 MHz, $CDCl_3$): δ 170.68, 170.44, 162.82, 161.89 (*d*, *J* = 251.77 Hz), 135.80, 135.45, 135.06, 132.77, 131.27 (*d*, *J* = 7.63 Hz), 128.27, 128.11, 123.95 (*d*, *J* = 8.9 Hz), 120.16 (*d*, *J* = 24.16 Hz), 119.46 (*d*, *J* = 24.16 Hz), 116.47, 70.97, 70.45, 70.15, 69.85, 62.60, 42.28, 39.13, 36.03, 35.80, 35.11, 28.69. HR ESI-MS (calc. for $C_{27}H_{32}FN_7O_7$): $[M+H]^+$ = 586.2410 (calc. = 586.2420). $[M+Na]^+$ = 608.2231 (calc. = 608.2239).

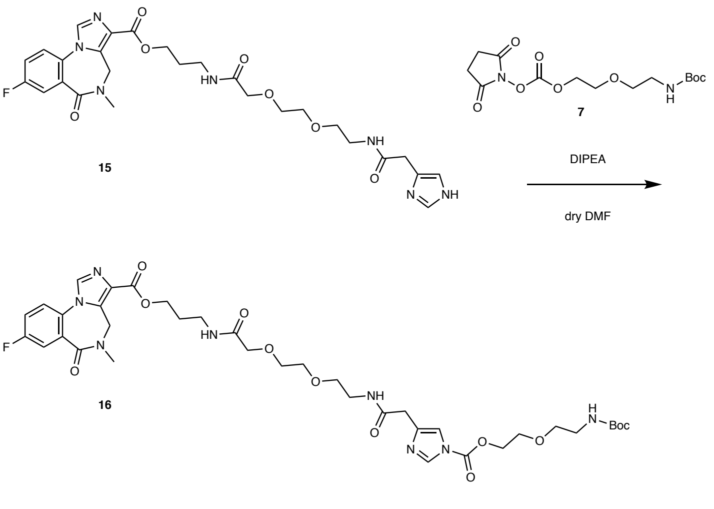


**1-(1-(11,11-dimethyl-9-oxo-2,5,10-trioxa-8-azadodecanoyl)-1*H*-imidazol-4-yl)-2,11-dioxo-6,9-dioxa-3,12-diazapentadecan-15-yl 8-fluoro-5-methyl-6-oxo-5,6-dihydro-4*H*-benzo[*f*]imidazo[1,5-*a*][1,4]diazepine-3-carboxylate (16):** To a solution of compound **15** (40 mg, 0.068 mmol) in dry DMF (1 mL) were added compound **7** (48 mg, 0.137 mmol, 2 eq.) dissolved in dry DMF (1 mL) and DIPEA (0.34 mmol, 60 μL, 5 eq.) at room temperature. The solution was allowed to stir for 2 h under Ar atmosphere. The solvent was evaporated and the residue was purified by silicagel chromatography with a linear gradient of 0–20% $CH_3OH/CHCl_3$ (12 CV) to give a compound **16** as a clear film (31 mg, 0.038 mmol, 56%). $^1$H NMR (600 MHz, $CDCl_3$): δ 8.12 (1H, *s*), 7.91 (1H, *s*), 7.78 (1H, *dd*, *J* = 8.8 Hz, *J* = 2.8 Hz), 7.38–7.34 (1H, *m*), 7.35 (1H, *s*), 7.18 (1H, *s*), 5.23 (1H, *bd*, *J* = 16.4 Hz), 4.87 (1H, *s*), 4.55 (2H, *dd*, *J* = 4.8 Hz, *J* = 2.8 Hz), 4.42 (2H, *t*, *J* = 10.4 Hz), 3.49 (2H, *dd*, *J* = 4.8 Hz, *J* = 2.8 Hz), 3.58–3.56 (3H, *m*), 3.46 (2H, *dd*, *J* = 12.4 Hz, *J* = 8.4 Hz), 3.33 (2H, *dd*, *J* = 9.6 Hz, *J* = 5.2 Hz), 3.25 (3H, *s*), 2.02 (2H, *quin*, *J* = 6.4 Hz), 1.44 (9H, *s*). $^{13}$C NMR (150 MHz, $CDCl_3$): δ 170.17, 169.55, 165.20, 162.93, 162.54, 160.97, 155.94, 148.37, 137.58, 137.16, 135.63, 135.15, 131.22 (*d*, *J* = 7.65 Hz), 128.39, 128.25, 128.24, 123.96 (*d*, *J* = 8.25 Hz), 120.07 (*d*, *J* = 22.80 Hz), 119.39 (*d*, *J* = 24.00 Hz), 114.98, 79.42, 70.98, 70.55, 70.33, 70.02, 69.84, 68.26, 67.12, 62.41, 42.33, 40.23, 39.21, 35.97, 35.69, 35.65, 28.91, 28.39. HR ESI-MS (calc. for $C_{37}H_{49}FN_8O_{12}$): $[M+Na]^+$ = 839.3353 (calc. = 839.3346).

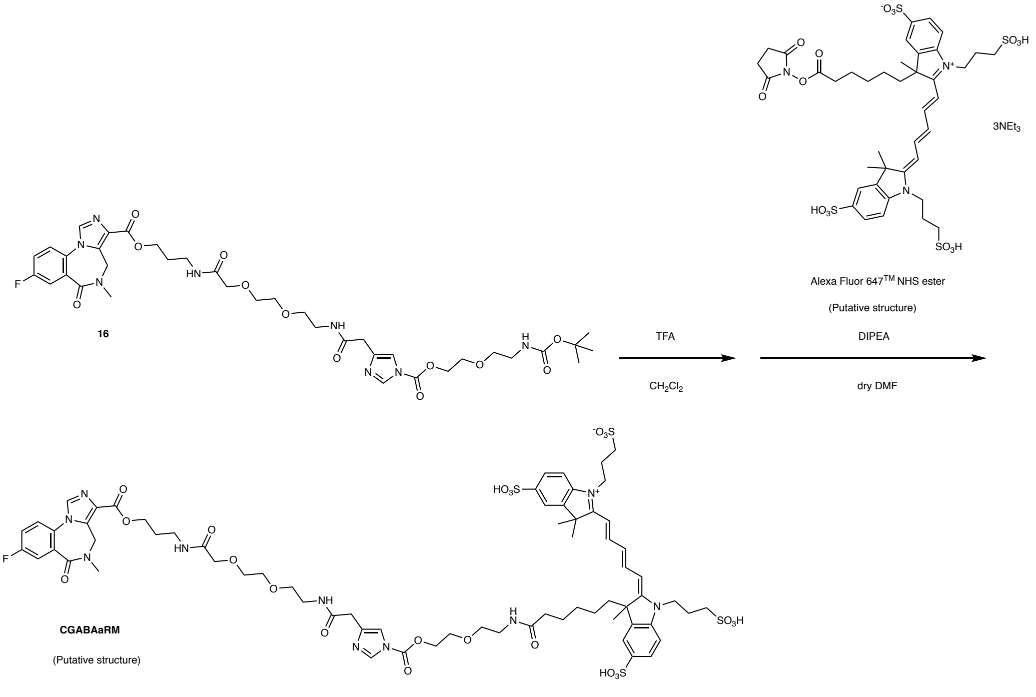


**CGABAaRM:** To a solution of compound **6** (0.65 mg, 0.8 μmol) in dry $CH_2Cl_2$ (1 mL), TFA (0.5 mL) was added and the solution was stirred at room temperature for 1 h. TFA was removed by co-evaporation with toluene under reduced pressure (2 mL x 3). The residue was dissolved in dry DMF (0.5 mL) and Alexa Fluor$^{TM}$ 647 NHS ester (1.0 mg, 0.8 μmol) and DIEPA (5.6 μL, 32 μmol, 40 eq.) were added. The reaction mixture was stirred at room temperature under Ar atmosphere for 2 h. The reaction mixture was diluted with $CH_3CN/H_2O$ (1/9) (2 mL) and purified by RP-HPLC (YMC-pack ODS-A, 250 x 10 mm, mobile phase; $CH_3CN$ : 10 mM $NH_4OAc$ aq. = 10:90 → 50:50 (linear gradient over 40 min), flow rate; 10 mL/min, detection; UV absorbance at 220 and 640 nm) to give **CGABAaRM** as a blue powder (1.1 mg, 0.72 μmol, 90% yield, determined by measurement of UV-absorbance). HR-ESI-MS (calc. for $C_{68}H_{83}FN_{10}O_{23}S_4$): $[M-2H]^{2-}$ = 777.2254 (calc. = 777.2255).

#### Synthesis of NLC

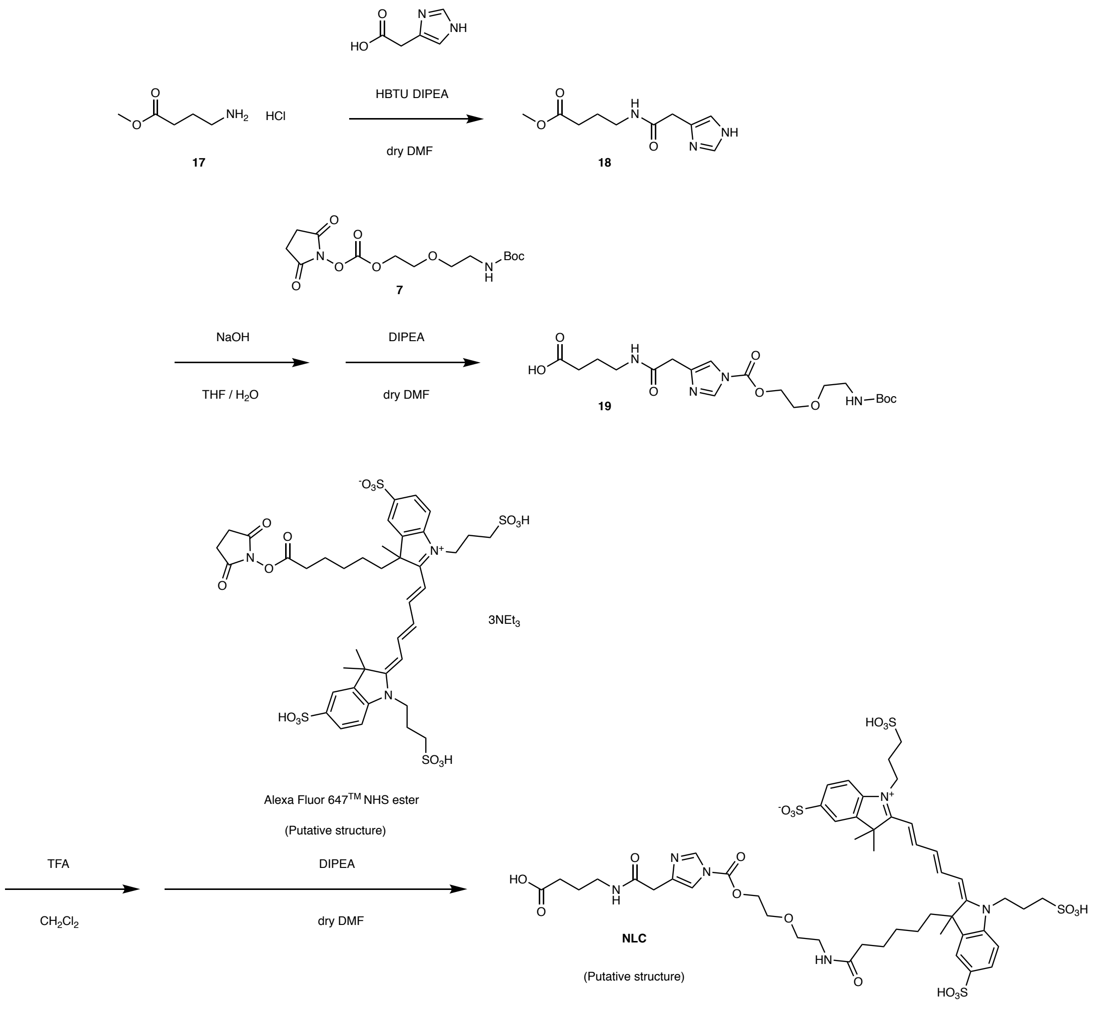


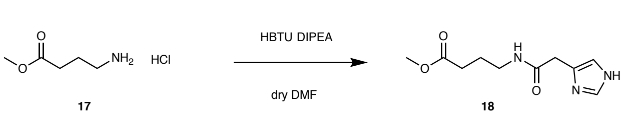


**Methyl 4-(2-(1*H*-imidazol-4-yl)acetamido)butanoate (18)**: To a solution of 4-aminobutyric acid methyl ester (**17**) (46.1 mg, 0.3 mmol) in dry DMF, 4-imidazolecarboxylic acid HCl (59 mg, 0.36 mmol, 1.2 eq.), HBTU (136.5 mg, 0.36 mmol, 1.2 eq.), and DIEPA (260 µL, 1.5 mmol, 5 eq.) were added. The reaction mixture was stirred at room temperature under Ar atmosphere for

16 h. After removal of DMF by evaporation, the residue was purified by silicagel chromatography with a linear gradient of 0–20% $CH_3OH$-1% $NH_3$/$CHCl_3$-1% $NH_3$ (12 CV) to give a compound **18** as a clear film (57.6 mg, 0.256 mmol, 85%). $^1$H NMR (600 MHz, $CDCl_3$): δ 8.42 (1H, *s*), 7.71 (1H, *s*), 7.40 (1H, *t*, *J* = 3.6 Hz), 6.92 (1H, *s*), 3.64 (3H, *s*), 3.57 (2H, *s*), 3.26 (1H, *q*, *J* = 6.0 Hz), 2.33 (2H, *t*, *J* = 7.2 Hz), 1.81 (2H, *quin*, *J* = 7.2 Hz). $^{13}$C NMR (150 MHz, $CDCl_3$): δ 173.798, 170.62, 135.03, 132.20, 116.47, 51.72, 38.93, 34.70, 31.34, 24.60. HR ESI-MS (calc. for $C_{10}H_{15}N_3O_3$): $[M+H]^+$ = 248.1002 (calc. = 248.1006). $[M+Na]^+$ = 248.0741 (calc. = 248.0745).

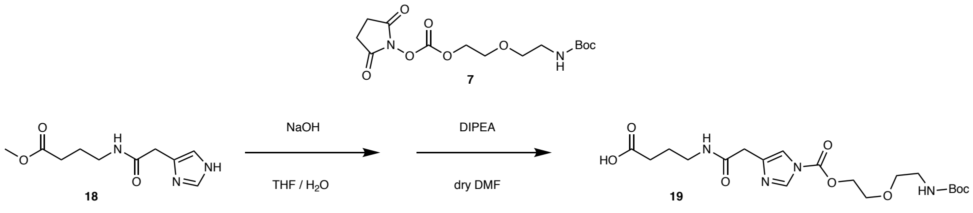


**4-(2-(1-(11,11-dimethyl-9-oxo-2,5,10-trioxa-8-azadodecanoyl)-1*H*-imidazol-4-yl)acetamido)butanoic acid (19):** To a solution of compound 1**8** (39.5 mg, 0.175 mmol) in THF/$H_2O$ (1/1) (4 mL), 1 M NaOH aq. (0.26 mL, 0.26 mmol, 1.5 eq.) was added and the reaction mixture was stirred at 40 °C for 30 min. The solution was neutralized by adding 1 M HCl aq. (0.26 mmol) and the solvent was removed by evaporation. The residue was dissolved in EtOH (2 mL) and filtered to remove NaCl salt. The solvent was evaporated and the residue was dried under reduced pressure to give a white form. After dissolving in dry DMF (0.5 mL), the compound **7** (45 mg, 0.13 mmol) and DIPEA (61 μL, 0.35 mmol, 2.7 eq.) were added. The reaction mixture was stirred under Ar atmosphere at room temperature for 1 h. The reaction mixture was diluted with $CH_3CN$/$H_2O$ (1/1) (2 mL) and purified by RP-HPLC (YMC-pack ODS-A, 250 x 10 mm, mobile phase; $CH_3CN$ : 10 mM $NH_4OAc$ aq. = 20:80 → 50:50 (linear gradient over 30 min), flow rate; 10 mL/min, detection; UV absorbance at 220 nm) to give a compound 1**9** as a clear film (34.3 mg, 0.078 mmol, 60%). $^1$H NMR (600 MHz, $CD_3OD$): δ 8.22 (1H, *s*), 7.46 (1H, *s*), 4.55 (2H, *m*), 3.80 (2H, *m*), 3.54 (2H, *t*, *J* = 6.6 Hz), 3.29–3.20 (6H, *m*), 2.27 (2H, *t*, *J* = 9.0 Hz), 1.78 (2H, *quin*, *J* = 8.4 Hz), 1.41 (9H, *s*). $^{13}$C NMR (150 MHz, $CD_3OD$): δ 178.44, 172.23, 158.49, 149.77, 138.58, 138.47, 116.74, 80.09, 71.04, 69.38, 68.51, 55.78, 41.19, 40.23, 36.05, 33.53, 28.74, 26.21. HR-ESI-MS (calc. for $C_{19}H_{30}N_4O_8$): $[M+H]^+$ = 443.2137 (calc. = 443.2136). $[M+Na]^+$ = 465.1955 (calc. = 465.1956).

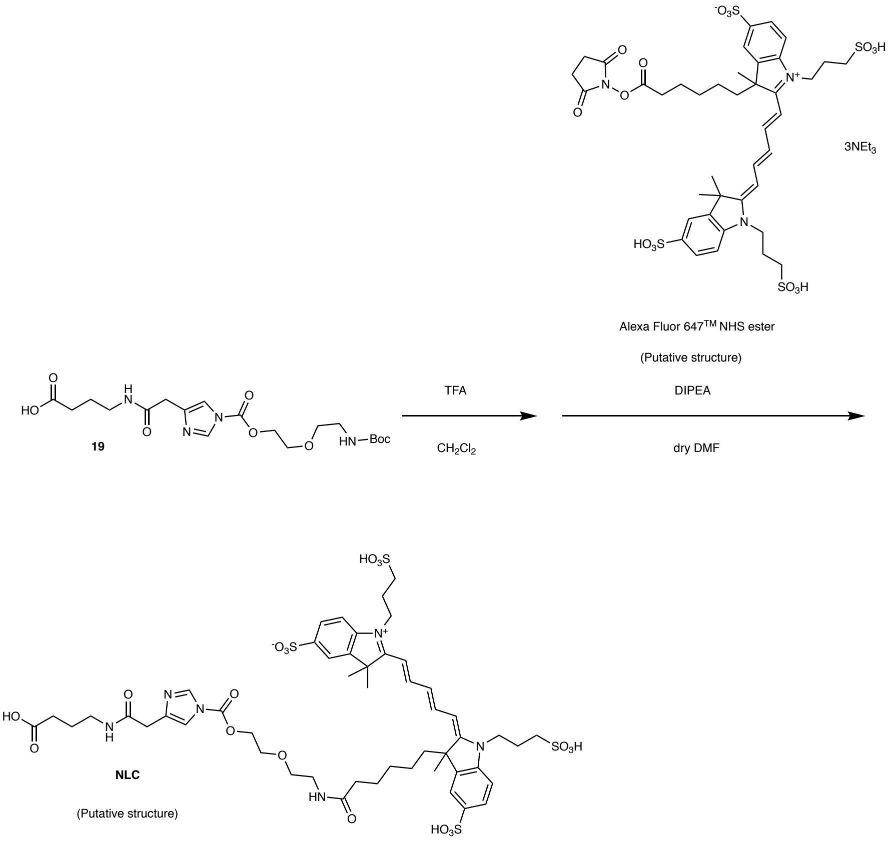


**NLC:** To a solution of compound **19** (0.36 mg, 0.8 μmol) in dry $CH_2Cl_2$ (1 mL), TFA (0.5 mL) was added and the reaction mixture was stirred at room temperature for 1 h. TFA was removed by co-evaporation with toluene under reduced pressure (2 mL x 3). The residue was dissolved in dry DMF (0.5 mL) and Alexa Fluor 647 NHS ester (1.0 mg, 0.8 μmol) and DIEPA (2.8 μL, 16 μmol, 20 eq.) were added. The reaction mixture was stirred at room temperature under Ar atmosphere for 2 h. The reaction mixture was diluted with $CH_3CN/H_2O$ (1/2) (2 mL) and purified by RP-HPLC (YMC-pack ODS-A, 250 x 10 mm, mobile phase; $CH_3CN$ : 10 mM $NH_4OAc$ aq. = 0:100 → 60:40 (linear gradient over 60 min), flow rate; 10 mL/min, detection; UV absorbance at 220 and 640 nm) to give **NLC** as a blue powder (1.1 mg, 0.72 μmol, 90% yield, determined by measurement of UV-absorbance)). HR-ESI-MS (calc. for $C_{50}H_{66}N_6O_{19}S_4$): $[M-3H]^{3-}$ = 393.1019 (calc. = 393.1016).

#### Synthesis of No AI Control (NAIC)

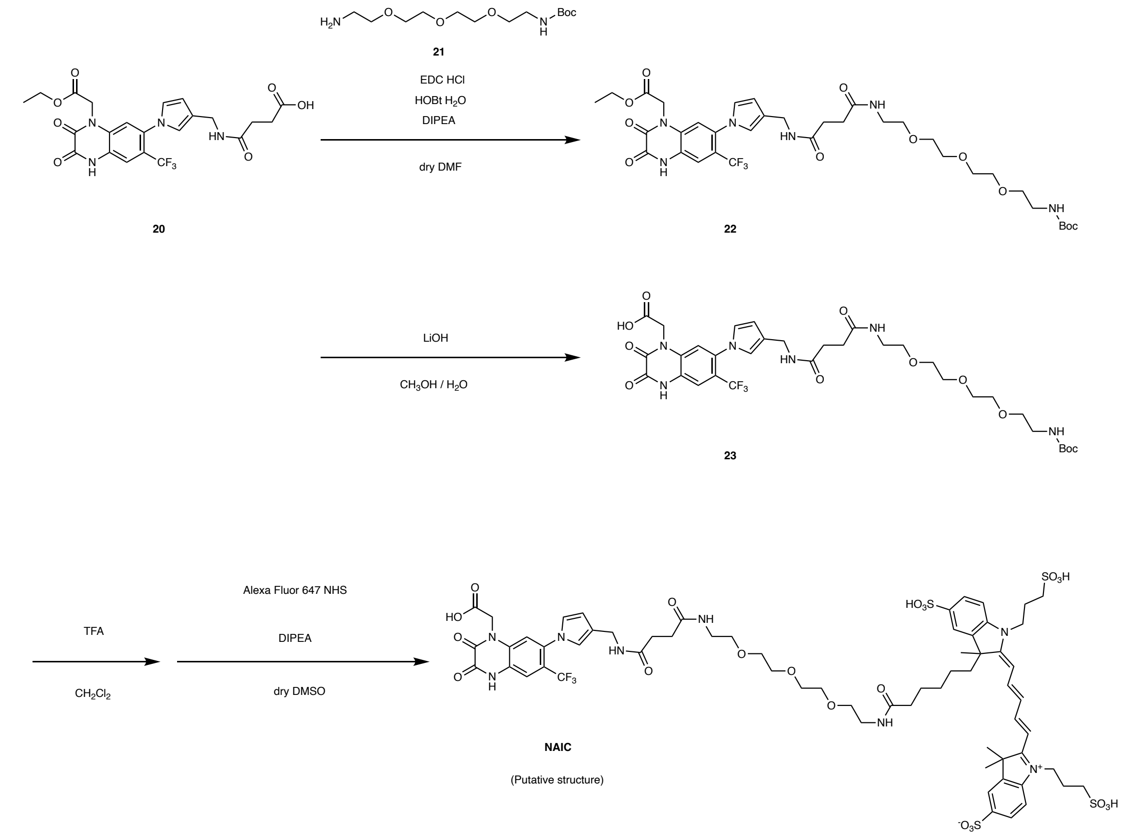


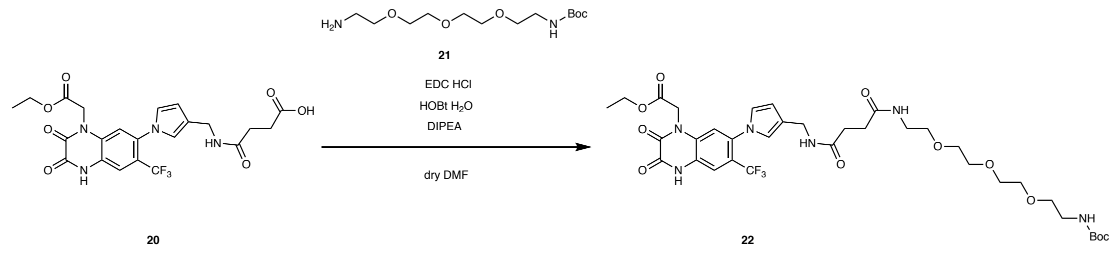


**ethyl 2-(7-(3-(22,22-dimethyl-3,6,20-trioxo-10,13,16,21-tetraoxa-2,7,19-triazatricosyl)-1*H*-pyrrol-1-yl)-2,3-dioxo-6-(trifluoromethyl)-3,4-dihydroquinoxalin-1(2*H*)-yl)acetate (22):** The compound **20** was synthesized according to the literature.[S1] A solution of **20** (120 mg, 0.235 mmol), **21** (89 mg, 0.304 mmol), EDC-HCl (59 mg, 0.31 mmol), HOBt-$H_2O$ (47 mg, 0.31 mmol), and DIEA (205 μL, 1.18 mmol) in dry DMF (1.0 mL) was stirred for overnight at room temperature under argon atmosphere. After removal of the solvent by evaporation, the residue was dissolved in $CHCl_3$ (50 mL) and the organic phase was washed with sat. $NaHCO_3$ aq. (10 mL, ×2), water (10 mL), and brine (10 mL). The resulting organic phase was dried over anhydrous $Na_2SO_4$, and then evaporated. The residue was purified by silicagel chromatography with $CH_2Cl_2$/$CH_3OH$ to give a compound **22** as a white solid (159 mg, 0.203 mmol, 86% yield). $^1$H NMR (400MHz, $CD_3OD$): δ 7.60 (1H, *s*), 7.35 (1H, *s*), 6.79 (1H, *s*), 6.77 (1H, *s*), 6.22–6.20 (1H, *m*), 5.04 (2H, *s*), 4.25 (2H, *s*), 4.22 (2H, *d*, $J$ = 7.2 Hz), 3.63–3.58 (8H, *m*), 3.52–3.48 (4H, *m*), 3.33 (2H, *d*, $J$ = 4.8 Hz), 3.20 (2H, *m*), 2.50 (4H, *s*), 1.42 (9H, *s*), 1.25 (3H, *t*, $J$ = 7.2 Hz). $^{13}$C NMR (100MHz, $CD_3OD$): δ 174.69, 174.21, 168.82, 158.40, 157.28, 155.1, 136.24, 131.12, 126.44, 125.50, 125.14, 123.48, 123.23, 122.94, 117.65, 115.70, 110.47, 80.05, 79.48, 71.54, 71.22, 71.03, 70.50, 63.25, 58.31, 45.70, 41.24, 40.39, 37.21, 32.31, 28.77, 18.38, 14.45. HR ESI-MS: m/z (calc. for $C_{35}H_{47}F_3N_6O_{11}$): $[M+Na]^+$ 807.3141 (calc. 807.3147).

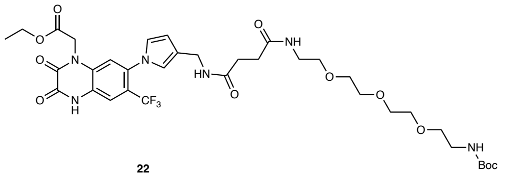

22

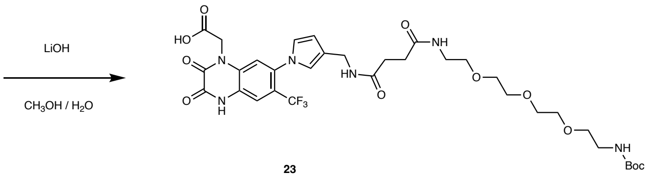


23

**2-(7-(3-(22,22-dimethyl-3,6,20-trioxo-10,13,16,21-tetraoxa-2,7,19-triazatricosyl)-1*H*-pyrrol-1-yl)-2,3-dioxo-6-(trifluoromethyl)-3,4-dihydroquinoxalin-1(2*H*)-yl)acetic acid (23):** To a solution of **21** (115 mg, 0.147 mmol) in methanol (5.0 mL), 0.5 N LiOH aq (2.0 mL, 1.0 mmol) was added. The reaction mixture was stirred at room temperature for 2.5 h. After adjustment of the pH to 4 with 1N HCl, the reaction mixture was concentrated and the residual solvent was azeotropically removed with toluene (×3). The residue was purified by silicagel chromatography with $CH_2Cl_2$/$CH_3OH$ to give a compound **23** as a white solid (100 mg, 0.132 mmol, 90% yield). $^1$H NMR (400MHz, $CD_3OD$): δ 7.61 (1H, *s*), 7.17 (1H, *s*), 6.80 (1H, *s*), 6.76 (1H, *s*), 6.20–6.19 (1H, *m*), 4.79 (2H, *s*), 4.25 (2H, *s*), 3.63–3.60 (8H, *m*), 3.52–3.48 (4H, *m*), 3.34 (2H, *d*, $J$ = 2.0 Hz), 3.21 (2H, *t*, $J$ = 5.6 Hz), 2.52 (4H, *s*), 1.42 (9H, *s*). $^{13}$C NMR (100MHz,

$CD_3OD$): δ 174.80, 174.21, 172.77, 158.38, 157.42, 155.73, 136.01, 131.44, 126.32, 125.70, 125.06, 123.30, 122.89, 122.68, 122.30, 117.74, 115.60, 110.49, 80.04, 71.46, 71.15, 71.02, 70.48, 58.28, 47.50, 41.19, 40.36, 37.19, 32.38, 28.77, 18.38. HR ESI-MS: m/z (calc. for $C_{33}H_{43}F_3N_6O_{11}$): $[M–H]^-$ 755.2867 (calc. 755.2869).

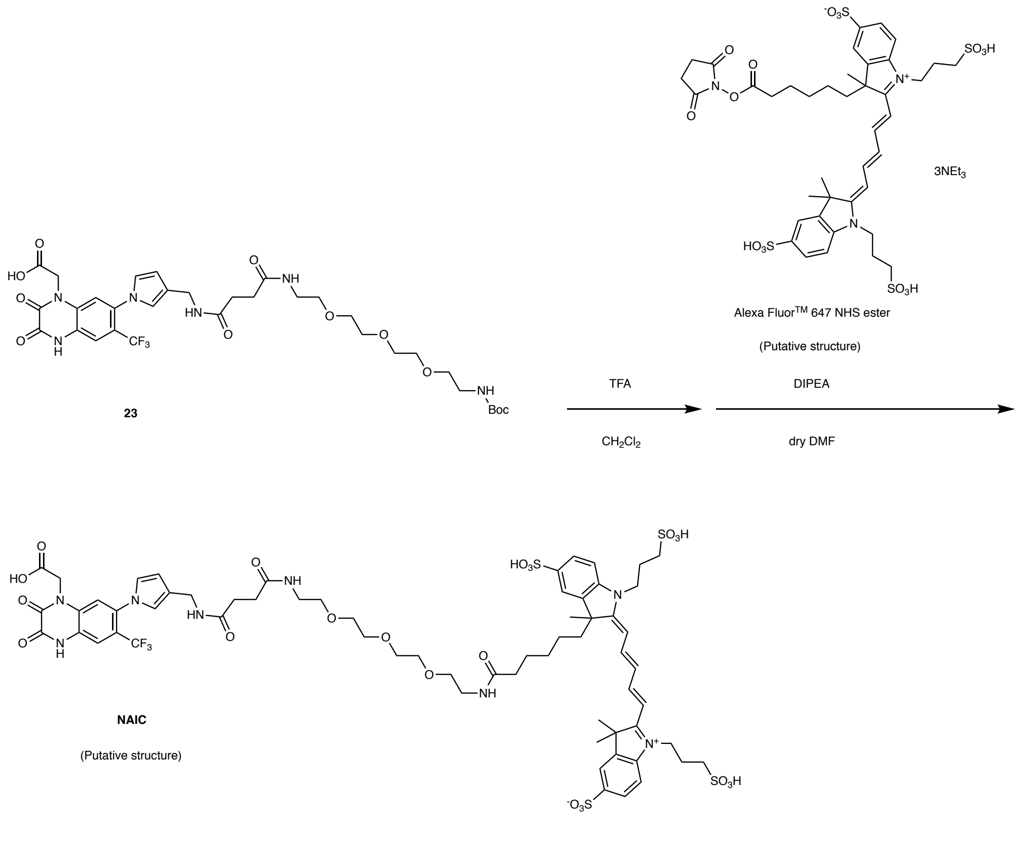


**NAIC:** To a solution of **23** (2.6 mg, 3.4 μmol) in $CH_2Cl_2$ (0.5 mL), TFA (0.10 mL) was added. The reaction mixture was stirred at room temperature for 1 h. After removal of the solvent by evaporation, the residual TFA was azeotropically removed with toluene (×3). The Boc-deprotected of **23** was dissolved in dry DMSO (500 μL). AlexaFluor™647 NHS ester (1.0 mg, 0.81 μmol) and DIEA (4.0 μL, 22 μmol) were added and the mixture was stirred at room temperature for 5 h. The reaction mixture was purified by RP-HPLC (Cosmocil 5C18AR2, 250 x 10 mm, mobile phase; $CH_3CN$ : 10 mM $AcONH_4$ aq. = 0:100 to 45:55 (linear gradient over 45 min), flow rate; 3 mL/min, detection; UV (220 nm)), giving **NAIC** as a blue solid (0.61 μmol, 75% yield, determined by measurement of UV-absorbance). HR ESI-MS: m/z (calc. for $C_{64}H_{79}F_3N_8O_{22}S_4$): $[M–3H]^{3-}$ 497.7973 (calc. 497.7975).

#### Synthesis of Lys-labeled product

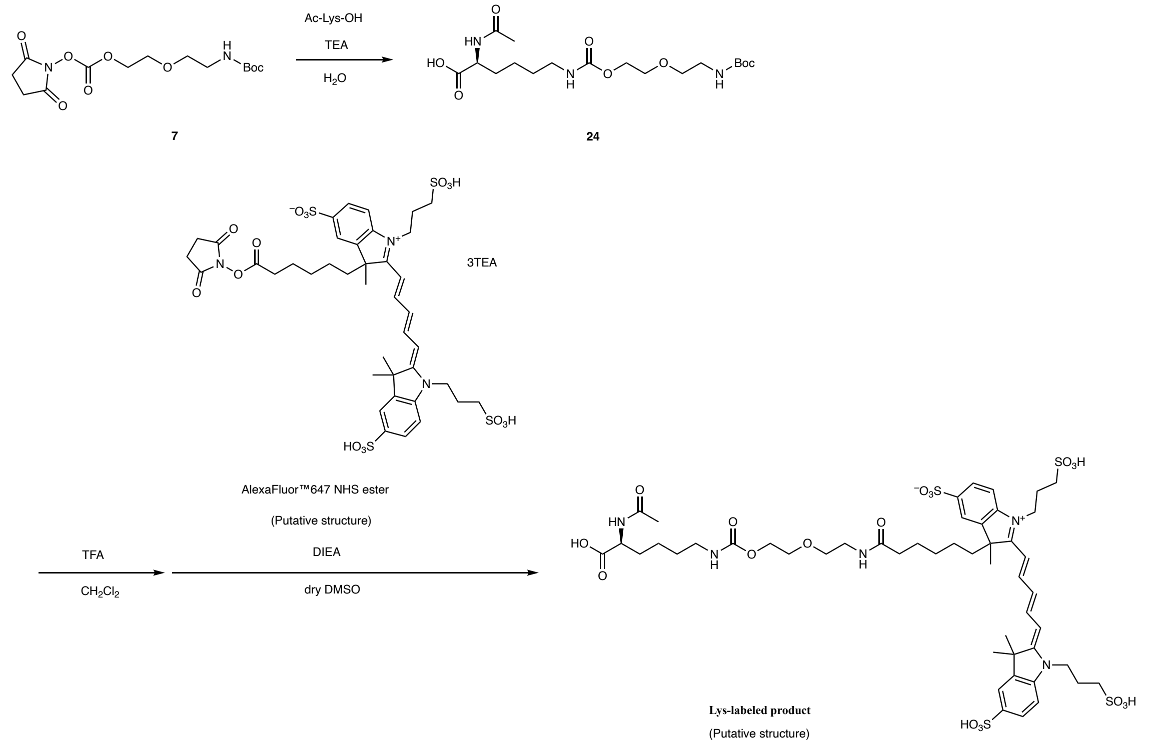


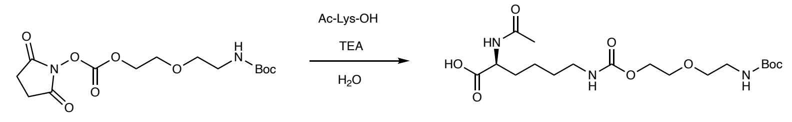


7 24

**(*S*)-18-acetamido-2,2-dimethyl-4,12-dioxo-3,8,11-trioxa-5,13-diazanonadecan-19-oic acid (24) :** To a solution of **7** (43.6 mg, 0.126 mmol) in acetone (1.0 mL), Ac-Lys-OH (47.4 mg, 0.252 mmol) in triethylamine 70 µL and water 500 µL was added. The reaction mixture was stirred at room temperature for 20 h. After removal of the solvent by evaporation, the residue was purified by silicagel chromatography with $CH_2Cl_2$/$CH_3OH$ to give a compound **24** as a white solid (21.8 mg, 0.0520 mmol, 41% yield). $^1$H NMR (400MHz, $CD_3OD$): δ 4.34 (1H, *dd*, *J* = 7.2, 4 Hz), 4.14 (2H, *t*, *J* = 3.6 Hz), 3.63 (2H, *t*, *J* = 4 Hz), 3.49 (2H, *t*, *J* = 4.4 Hz), 3.20 (2H, *t*, *J* = 3.6 Hz), 3.10 (2H, *t*, *J* = 5.6 Hz), 1.98 (3H, *s*), 1.87–1.80 (1H, *m*), 1.72–1.67 (1H, *m*), 1.64–1.37(13H, *m*, *J* = 4 Hz). $^{13}$C NMR (100MHz, $CD_3OD$): δ 175.52, 173.36, 158.99, 158.44, 80.09, 70.99, 70.41, 64.93, 53.65, 41.41, 41.24, 32.25, 30.46, 28.75, 26.27, 24.10, 22.34. HR ESI-MS: m/z (calc. for $C_{18}H_{33}N_3O_8$): $[M+Na]^+$ 442.2159 (calc. 442.2160).

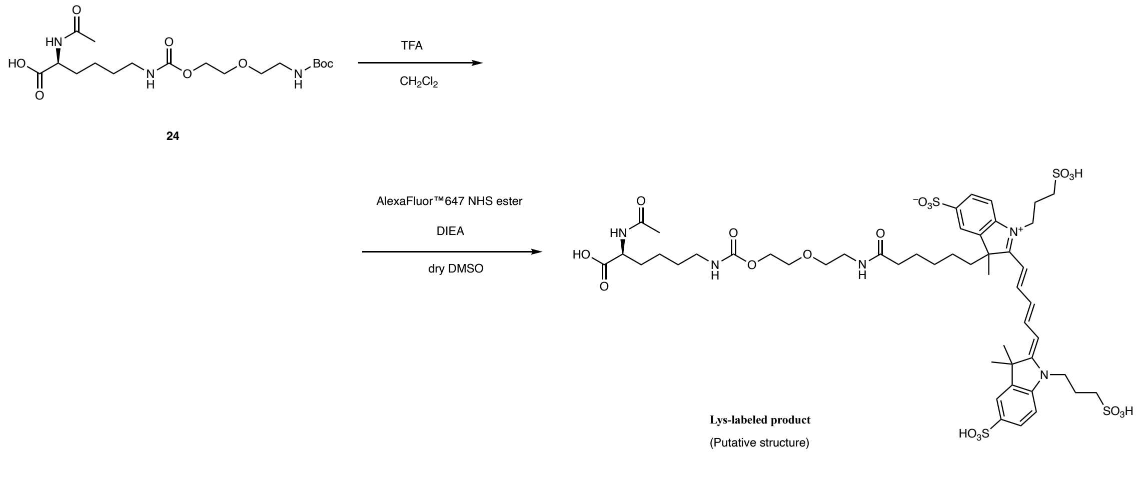


**Lys-labeled product:** To a solution of **24** (1.3 mg, 3.1 μmol) in $CH_2Cl_2$ (1.0 mL), TFA (0.20 mL) was added. The reaction mixture was stirred at room temperature for 2 h. After removal of the solvent by evaporation, the residual TFA was azeotropically removed with toluene (×3). The Boc-deprotected of **24** was dissolved in dry DMSO (500 μL). AlexaFluor™647 NHS ester (1.0 mg, 0.81 μmol) and DIEA (2.0 μL, 11 μmol) were added and the mixture was stirred at room temperature for 19 h. The reaction mixture was purified by RP-HPLC (Cosmocil 5C18AR2, 250 x 10 mm, mobile phase; $CH_3CN$ : 10 mM $AcONH_4$ aq. = 0:100 to 50:50 (linear gradient over 50 min), flow rate; 3 mL/min, detection; UV (220 nm)), giving **Lys-labeled product** as a blue solid (0.60 μmol, 74% yield, determined by measurement of UV-absorbance). HR ESI-MS: m/z (calc. for $C_{49}H_{69}N_5O_{19}S_4$): $[M–3H]^{3–}$ 385.4420 (calc. 385.4417).
